# VNTR prediction on sequence characteristics using long-read annotation and validation by short-read pileup

**DOI:** 10.1101/2020.07.17.185983

**Authors:** Diederik Cames van Batenburg, Jasper Linthorst, Henne Holstege, Marcel Reinders

## Abstract

Tandem repeats (TRs) are contiguously repetitive sequences with a high mutation rate. Several human diseases have been associated with an expansion of TR, a mutation which constitutes a change in their number of repetitions. Nevertheless, these Variable Number Tandem Repeats (VNTRs) have not been included in many genome-wide studies. The reason is that VNTR genotyping is inaccurate using short-read sequencing while new technology like long-read sequencing is expensive and lacks throughput.

Here, we propose a sequence based random forest classifier that is able to predict variable expansion of TR regions, given by incomplete VNTR annotation from long-read sequencing of 5 haplotypes. The classifier mainly predicted VNTRs using the features TR length. The second most used feature is a novel finding: the Mfold predicted likelihood of self-folding for which more stable foldings are correlated with VNTRs. We validated VNTR candidates predicted by this classifier by clustering short-read pileup patterns compared across 17 genomes. TRs labeled VNTR by the classifier showed similar local variance in their pileup profiles.

**Supplementary information:** Supplementary data are available at bioRxiv

## 1 Introduction

While originally deemed insignificant, the importance of Tandem Repeats(TRs, see Box 1) has been progressively established. Past studies have found TRs to be functional and disease causing elements as well as one of the driving forces in human evolution (Liang *et al.*, 2015; Sonay *et al.*, 2015).

#### Box 1: Tandem Repeat definitions

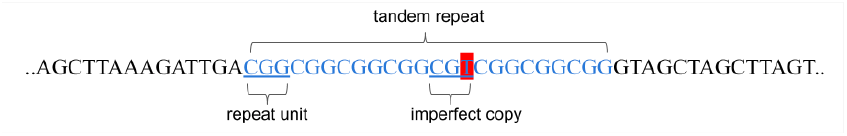

A Tandem Repeat (TR) consists of multiple contiguous occurrences of a motif or Repeat Unit(RU). The RU consists of a set sequence of nucleotides, but TRs may show small imperfections in individual repeats. The number of repeats of the motif is called the copynumber or the RU count of the TR. In many studies, TRs are separated by their motif length into long TRs (> 6 basepairs) and STRs (| 6 basepairs).

A subset of TRs show expansion: a growth or reduction of the RU count. These Variable Number Tandem Repeats (VNTRs) are defined as TRs for which a variant in RU count is observed in the population. In this paper, we refer to TRs for which no expansion has yet been observed static TRs (stTRs). This is a simplification since actually there likely is some degree of variation present within the human population for all TRs. The most recognized theory for the source of TR expansion as described by Fan and Chu, 2007 is DNA slippage events during DNA replication or repair (Box 2). Another correlated event is double strand breakage (Audano *et al.*, 2019) which is enriched in subtelomeric regions and as a side-effect of DNA repair mechanisms (de la Chapelle and Peltomäki, 1995).

Many diseases have been related to an expansion in a specific VNTR to a certain range of copynumbers or are at least partly caused by VNTR variants. Many of these are neurodegenerative diseases like frontotemporal dementia, Huntingtons disease, different types of ataxia and amyotrophic lateral sclerosis(ALS) as reviewed in Hannan, 2018 and Bakhtiari *et al.*, 2018. Furthermore, there are several complex diseases for which VNTRs have been found to be one of the genetic factors, such as bipolar disorder, ADHD and Parkinson’s disease (Benedetti *et al.*, 2008; Franke *et al.*, 2010; Kirchheiner *et al.*, 2007). Nevertheless, as reviewed by Brookes, 2013, VNTRs have been underrepresented in the search for disease causing genes for complex diseases with missing heritability. Family studies show that these diseases are caused by inheritable traits, but regular screenings have not recovered the disease causing genes to explain them fully (Manolio *et al.*, 2009).

#### Box 2: DNA slippage events

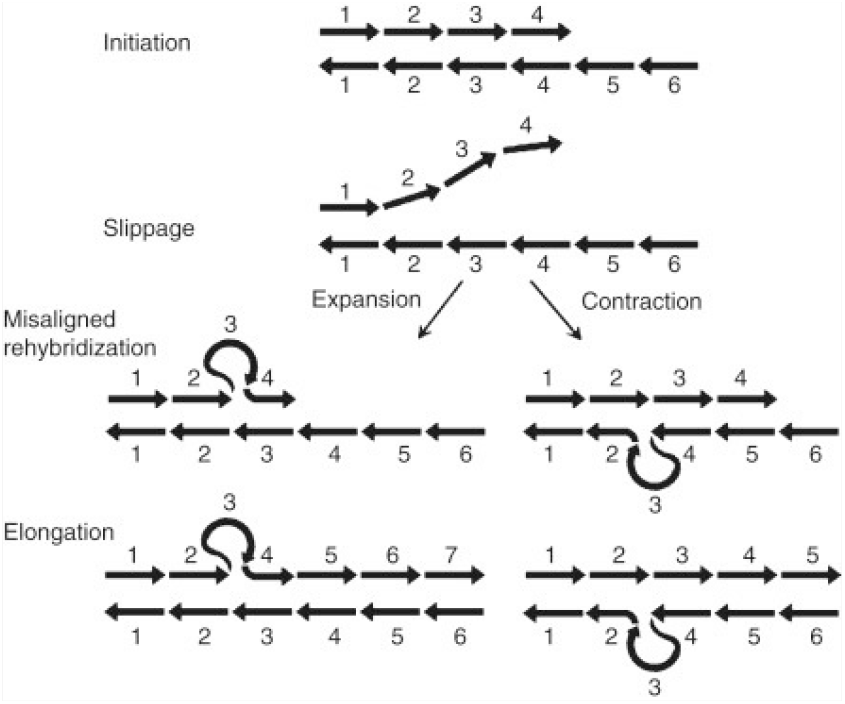

Source: Jorda and Kajava (2010).

In TR expansion the nascent strand may loop and then miss-align back to the already transcribed template. This reference region that is skipped back will then be replicated a second time (region 5 repeats 4). In contraction, the strand that is used as template may loop out, causing the looped part (region 3) to be omitted from replication.

VNTRs have not been tested nearly as much as regular genes because the two most used sequencing techniques are not suitable for determining TR RU count: SNP chips in Genome Wide Association Studies (GWAS) and short-read sequencing. For testing a large amount of genes for partial contribution to complex diseases, GWAS test for common variations in a large set of single nucleotide loci (Hardy and Singleton, 2009). This does not help for genotyping TR length variants because they inherently require sequencing regions. Furthermore, when sequencing many regions, the chosen method is shotgun sequencing for its cost and time efficiency. Applied to VNTR genotyping this poses two problems: 1) at the amplification step, the Polymerase Chain Reaction (PCR) artificially induces in vitro TR expansions, and 2) TRs contain a long repetitive sequence, so that reads that fall within the repetitive region can not be uniquely mapped to any one position.

One way that extra information can be gained for TR read mapping is the paired end set up of common short-read sequencing methods (Box 3), but its use is limited because read pair distances are a distribution instead of a known value. Moreover, this information is of no use when even the complete fragment is shorter than the TR.

These shortcoming are overcome by long-read sequencing, in which reads overlap the complete TR but current costs of long-read sequencing, however, prohibit this technique to be applied in large cohort studies such as current GWASs.

### 1.1 Previous work

Multiple models have been developed to estimate TR copynumber (RU count). One program called ExpansionHunter by Dolzhenko *et al.*, 2017 makes use of paired end reads in PCR-free short-reads. It produces confidence intervals of TR length based on the counts of flanking reads and anchored in-repeat reads. These can also provide lower bounds for TR length. Unanchored in-repeat reads are recovered from off-target positions with low mapping quality, selecting by high sequence similarity to TRs. They formulate a binomial model of expected number of reads mapped on a TR of certain length given the read length and average pileup. The inverse returns the estimated TR length based on read counts.

In the program adVNTR developed by Bakhtiari *et al.*, 2018 a Hidden Markov Machine(HMM) is trained to estimate RU count. Each specific TR is modelled by a unique HMM that includes separate sections for the left and right flanks of the TR as well as a repetitive middle section for RUs. First, all reads overlapping a specific TR are recruited by testing the likelihood its HMM produced it. Secondly, all reads are processed again by the HMM while keeping track of the number of times that the RU HMM section was completed to produce the estimated RU count.

A statistical method based on paired end distance was developed by Cao *et al.*, 2013 in the program STRviper. In case of anchored in-repeat TRs, any aberration in the distance between the range versus expected range can indicate a sequence in between that contains more or less RUs compared to the reference. Through Bayesian inference, a probability and confidence interval for RU count is returned for STRs.

Although other methods like Southern blotting and repeat-primed PCR perform well in genotyping TRs, these require high amounts of time and effort for each TR (Dolzhenko *et al.*, 2017).

Ideally VNTRs can become part of the large scale GWAS along with regular gene variants. This would require a method to detect TR copynumber in a cohort-like fashion.

An easier problem would be to detect only variation of TR length. This requires no estimate of the length, but only significant differences of TR properties between genome samples. This might still be accomplished by short-read sequencing data, as differences in the amount of reads mapped to a TR. A pre-requisite is then that the VNTR regions are known beforehand, so that read counts to these regions can be established. This can be realized by predicting whether a TR region is variable or not based on sequence properties of the TR region.

This approach is used by Näslund *et al.*, 2005, who created a linear regression classifier on sequence properties returned by the TRfinder program. They found the best result using the four predictors: RU count, GC dinucleotide bias, entropy and match percentage between repeats. Especially TRs with high copynumber and pure repeats were associated with expansion probability. Since our current level understanding of TRs is fairly limited by current analysis techniques, we are not in a position to test hypotheses on TR emergence or TR/VNTR transitioning. We therefore aim to create a predictor for expansion-prone TRs first.

Our study makes use of a small set of longread data as ground truth to develop an indicator on only short-read data for TR expansion. Here, we follow a two-way approach with a sequence based classifier to find candidate regions of VNTRs followed by validation by analysis of short-read pileup. Our classifier aims to expand on the feature set used by Näslund *et al.*, 2005 towards more general pattern features such as k-mers and folding tendency of single strand TRs as well as genomic context.

TR short-reads are analyzed in terms of local pileup along the TR through unsupervised machine learning. We aim to find patterns in single or multi genome comparative pileup profiles that are typical of disparaging TR lengths.

## 2 Methods

Detailed TR coordinates and detailed TR information as generated by the TandemRepeatFinder(TRfinder, Benson, 1999) program was obtained from the ucsc table simple repeats track. TRfinder by default restricts motif length above 500 and a minimum alignment score equivalent of 25 perfect matches. TRs that are fully contained in other TR regions are considered redundant and left out from the dataset, leaving 610. 685 unique TRs in chromosomes 1 through 22. Summarizing TRfinder data for merged cases was done by selecting a representative sub TR with the best motif alignment score as given by TRfinder.

To produce initial VNTR labeling, the HG38 (the most recent human assembly reference genome) known TRs in 4 haplotype Pac-Bio longread human genomes were compared. A subset of TRs were labeled VNTR if sufficient structural variant level (>50bp) of difference in TR length was observed in the samples. The condition for VNTR labeling was overlap of a region labeled as SV with a TR, including any partial overlap of any size. Of the remaining TR set, 11.873 are labeled VNTRs, giving us a baseline VNTR ratio of 1.9%.

This long-read derived labeling is taken as TR expansion ground truth. Keep in mind that the TR variation, studied here, is between just 5 haplotypes and thus constitutes a heavy underestimation of TR variability. With respect to the paper by Audano *et al.*, 2019 on novel structural variant detection using long-read sequencing, we estimate a conservative 45% of VNTRs are covered by the number of samples.

### 2.1 Classifier

We constructed a classifier to detect the VNTR class in TRs, with the other class being static length TRs. This is done mainly on sequence based characteristics and some genomic context.

A featureset from multiple datasources is extracted, making use of TR genomic position as index. Table 1 shows the features that were extracted with their sources.

**Table 1.** Features considered for VNTR classifier and their usage by the final random forest classifier.

| Name | Description | source | usage |
| --- | --- | --- | --- |
| seqlen | TR length in bp | simple repeats* | 0.276 |
| Mfold | Free energy change of folding | Mfold | 0.158 |
| copy_num | Repeat unit count | simple repeats* | 0.095 |
| motiflen | ConsensusSize of repeat unit | simple repeats* | 0.068 |
| cgcon | Count of C or G over sequence length | new | 0.052 |
| centro | Distance to centromeric region | centromeres* | 0.049 |
| matches | Percentage of matches between adjacent copies overall | simple repeats* | 0.049 |
| telo | Distance to nearest telomeric region | cytoBand* | 0.035 |
| pcT | Count of T nucleotides over sequence length | new | 0.033 |
| pcC | Count of C nucleotides over sequence length | new | 0.033 |
| pcG | Count of G nucleotides over sequence length | new | 0.028 |
| transcr_up | Distance to nearest transcription start site, upstream TR position | GENCODEv29* | 0.028 |
| transcr_dwn | Distance to nearest transcription start site, downstream TR position | GENCODEv29* | 0.028 |
| pcA | Count of A nucleotides over sequence length | new | 0.028 |
| indels | Percentage of indels between adjacent copies overall | simple repeats* | 0.019 |
| chrom | Source chromosome | cytoBand* | 0.019 |
| genic_pos | Genomic context: intergenic, intronic, non-coding exon, coding exon | GENCODEv29* | 0.003 |
| mfoldNaN | Marks undefined mfold value | Mfold | unused |
| 3-mer | Counts of all 3 nucleotide combinations | new | unused |
| 2-mer | Counts of all 2 nucleotide combinations | new | unused |

\*from the ucsc database, GRCh38/hg38 assembly

#### General rationale

Features were chosen to reflect sequence characteristics (length, nucleotide ratios and k-mers), TR properties (motif length, copyNumber and motif imperfections) and genomic context (chromosomal position, gene proximity and self-folding).

#### Feature description

Sequence characteristics were considered because the TR expansion mechanism may be driven or inhibited by nucleotide-encoded patterns. TR features such as motif length and total length are determined by their sequences as well but these patterns manifest themselves at a higher order.

###### Box 3: Paired end read mapping of TRs

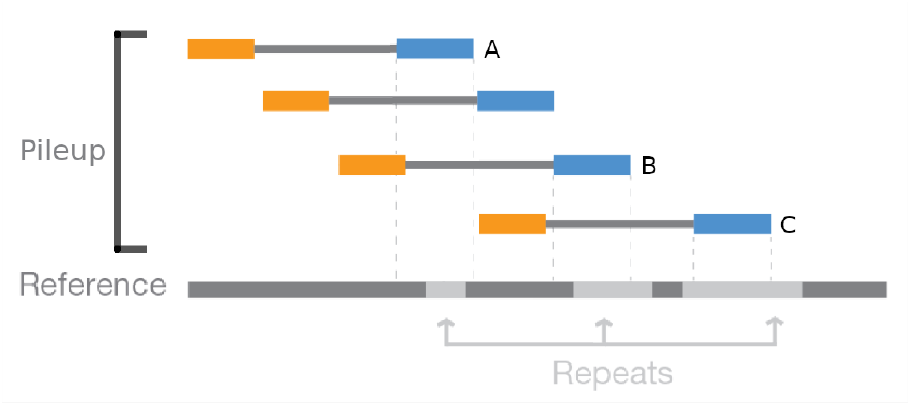

Reads are sequenced in pairs, located at the ends of a fragment. There is a known average length of unsequenced region between them, commonly modeled as a normal distribution (Cao *et al.*, 2014). Many of these are mapped on a reference genome based on maximum similarity with redundancy, so that multiple reads will overlap. The count of overlapping reads mapped on any location is called its read pileup.

Paired end reads contain extra information because of the expected distance between them. If the complete fragment with both ends falls within a TR, there is no additional information gained from the paired ends. Otherwise, three variants exist where information in pairs can help to uniquely map repetitive TR regions.

In the example we see A: Spanning read; just one suffices for unique mapping and TR size is known. B: Flanking reads; uniquely mapped using the non TR flanks. TR size can be estimated from them. C: In-repeat reads; can be anchored by the read mate (orange here) if it falls outside of repetitive regions. Its within-TR mapped location is estimated from the distance between the read pairs.

Furthermore, TR patterns are expected to have a relation to TR expansion. Indeed, the presence of TR patterns alone causes in vitro TR expansion behavior during PCR as described by Dover, 1995. Repeat unit purity (Percent Matches between adjacent repeat copies) may be witness to TR age or expansion events that were interceded by point mutations. They could also disrupt the other TR features by introducing noise. The k-mer frequencies present in motifs form a general representation of sequence patterns for stretches of contiguous nucleotides of length *k*. We tested k-mers up to *k* = 3 to limit the number of features generated. The k-mer feature was represented as the ratio of a k-mer count compared to all counted k-mers in the particular TR.

The genomic context features are expected to mostly reflect genetic pressure against expansion, such as proximity to transcription start sites. Moreover, whatever mechanism is directly causing TR expansion is likely modulated by other interactions with the TR sequence and flanking regions as well. It is known for example that SVs and TRs are more concentrated in telomeric regions where double strand breakages are more prevalent Audano *et al.*, 2019, although there has been no indication of increased expansion probability. The theory of DNA slippage as mechanism directly causing TR expansion assumes one of the strands is looping out of the DNA polymerase during replication. The stability of this loop and the likelihood of this loop forming in the first place may be strongly correlated to the nucleotide makeup. Furthermore, TR folding may be the mechanism underlying modulation of nearby gene translation attributed to TRs, where the folded structure may restrict promotor accessibility. For this, the Mfold program was used to estimate the largest free energy change resulting from optimally folding the single strand TR on the level of secondary structure. Generally, TR expansion can be disruptive or beneficial and is under evolutionary selective pressure. Therefore, we include the ordinal functional context of the TR, scaling in relevance from low relevance in intergenic, to intronic, non-coding exonic and finally coding exonic regions. To confirm previous reports of uniform distribution of expansion rate over chromosomes we also included this categorical feature.

#### Mfold feature

The likelihood of the sequence to engage in auto-folding was assessed using the Linux application Mfold version 3.6 by Zuker, 2003 on single strand DNA using default settings. The auto-folding predicted free energy change, Δ*G*, which we considered as feature. However, this showed a large ratio of missing data (28.7%) that requires special interpretation. The Mfold program may return an undefined value for an input sequence if it was unable to find a stable folding. Since a strong negative Δ*G* corresponds to a more stable folding, it may be sensible to replace missing values with a high > 10000 value as suggested in Zuker, 2003. However, this shows a strong outlier-like behaviour which will dominate the scale of the numeric feature, because there are no other positive energy changes. This has lead us to prefer a value of 0, which indicates no inclination to change its unfolded structure.

In exploratory case studies, we trained multiple classifiers with: 1) zero-imputed Mfold values, 2) excludig TRs for which Mfold has missing values, 3) adding a categorial feature with one-hot encoding whether Mfold feature was missing or not, together with a feature of zero-imputed Mfold values.

#### Data variants

Additional variants of the classifier were learned for handling categorical data and motif length subgroups. Methods for integration of categorical and continuous features were compared using simple label replacement by integer or one-hot encoding. Data variants were divided into the categories of STRs and long tandem repeats. Comparison between the two categories appears customary in TR literature and it may uncover unknown differences between them. For choosing the final representation of the data, we performed exploratory training and testing on a simple 100 tree classifier for a few different feature subsets as well as different solutions for missing mfold data and representation of categorical features.

#### Classifier design

The chosen classification algorithm is an ensemble method of Random Forest Classification (RFC) with 3000 estimators. We chose this classifier after a brief exploration the RFC, K-nearest neighbors and support vector machine classification algorithms which showed that the RFC outperformed these classifiers on F1 performance. Because the data is strongly unbalanced towards static TRs, the training scheme applies a stronger weight on the minority class, giving it equal representation of the two classes. All features in all datagroups are scaled by subtracting the mean and setting scaling to unit variance giving each feature equal priority to the classifier. In addition, the telomeric and centromeric distance is scaled by the distance to the closest telomere on the same chromosome arm in bp as a fraction of the length of the corresponding arm.

#### Implementation details

TR dataset generation and VNTR labeling was done using BEDtools (Quinlan and Hall, 2010). Implementation of the classifier was done in Python making use of the popular scikit-learn module (Pedregosa *et al.*, 2011) for the classifier pipeline as well as imbalanced-learn (Lemaître *et al.*, 2017) for dealing with unbalanced data.

#### Classifier description

A random forest classifier bootstraps the data over multiple uncorrelated simple decision tree classifiers and decides by majority vote which leads to improved generalization (Breiman, 2001). Apart from a most likely label it also allows to estimate the label probability by taking the fraction of trees that support the decision. In our configuration, each tree also uses a random subset of features with size 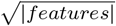, to further decouple the subtrees. Hyperparameters to be set were class weights, which act effectively as randomly duplicating VNTR samples in the training for the ratio given by the class weight. Weights are set in each tree so that it counters the local imbalance in its bootstrap subsample by applying an inversely proportional weighting to the class ratio in the subsample: 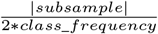. The number of samples in a node at which a tree will stop splitting it for purification any further is set at 35. We set the number of trees at 3000 although the training and evaluating of the classifier still increased slightly.

#### Classifier training scheme

A combined training and optimization set of 90% of samples was selected using random stratified subsampling without replacement. This was first used to select the best classifier with the best hyperparameters using 5 fold cross-validation for each configuration. The chosen classifier with the best found hyperparameters is then trained on the full 90% training data to produce predicted labels and tested on the remaining 10% test samples. Hyper-parameter optimalization confirmed better performance for increased weighting on the minority class instead of undersampling the majority class.

To produce more reliable classifier predictions of VNTR probability, calibration by isotonic regression is applied to the training set. This uses a 5 fold cross-validation procedure, effectively training on 80% of the training set and applying isotonic regression on the remaining 20% iteratively. Isotonic regression first divides the sorted predicted probability into intervals by linear regression enforcing a non-decreasing piece-wise linear approximation. The predicted probability of each linear interval is then mapped to the fraction of VNTRs in the corresponding subset of training samples (Boström, 2008). Skewed probability estimates are often observed for random forest classifiers which have a tendency to predict close to 0.2 and 0.9 but rarely close to 0 and 1 as explained by Niculescu-Mizil and Caruana, 2005. Calibrated probabilities as predicted by all 5 folds are averaged to produce the final predicted probabilities as the final classifier.

#### Evaluation measures

Classifier performances are evaluated by F1-score, precision, recall and AUC (area under curve) of ROC (Receiver Operator Characteristic) and AUC of Precision-Recall Curve. For evaluation, accuracy is the least relevant metric, because the majority of TR samples is static, and easily distinguished as such which leads to a naturally high accuracy. Other metrics which focus on the relevant samples, being the VNTRs, are more relevant and represents cases which are harder to classify correctly. As we are interested in detecting VNTRs we are primarily interested in a good recall of VNTRs, while at the same time having a high precision to avoid needless validation efforts. Specifically because of resource limitations and statistical power diminishing with the number of multiple candidate testing the first priority remains presenting a limited set of regions of interest. Therefore, we consider a composite performance metric of recall and precision in the form of F1-score the most suitable. However, the classifier is expected to heavily overestimate the number of false positives (FP) that should be allocated to true positives (TP), given that the classifier misses out on some 45% of VNTRs (due to our initial annotation of TRs). The same holds true for a subset of true negatives (TN), that in actuality should be part of false negatives (FN).

Given that the total set of positive cases is underestimated by a factor of 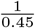 we can estimate the impact on some of the metrics used. Because the unlabeled fraction is estimated based on the positive cases, it has the most impact on the precision score 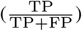. We assume the number of missing positive cases would become spread over FN and TP cases in the same proportion as their prevalence in the currently labeled cases. In that case the TP number can be corrected by the correction factor 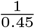 to generate a corrected precision score: 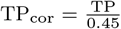. This will result in a precision score that is more optimistic and moves in the direction of the real performance under completely labeled data (given the assumption). A corrected FP number is expected to drop because the missing TP cases will be transitioning from either FP or TN cases. We choose not to correct the FP number because the number of TNs is much bigger than the number of FPs. However, correcting negative cases would only further improve the corrected precision score. The recall score 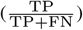, could also be calculated using corrected parameters. We choose to leave it unchanged, because our knowledge of the fraction of unlabeled VNTRs has no influence on the correction.

The chosen metric, F1-score, which is a harmonic mean over precision and recall, can similarly be corrected by substituting the corrected precision score. This corrected F1-score is used as evaluation scorer in setting the classifier hyperparameters.

#### Classifier evaluation

Features were first related to TR expansion individually by Pearson’s *r* correlation in the case of continuous variables and Cramér’s V for categorical variables, both of which represent correlation strength.

### 2.2 Validation by pileup analysis

This section describes how we validated detected VNTRs using short-read sequencing data. We used unsupervised learning to cluster TRs with similar pileup characteristics. Subsequently, we analyzed the position of sequence classifier predictions within the pileup clustering. In particular, we aimed to find evidence of FP cases that are actually new candidates by their proximity to TP cases in the clustering.

#### Short-read sequencing data

Raw bwa-mem aligned short-read data was retrieved from the 1000 Genomes Project Consortium *et al.*, 2015. Specifically, 17 human PCR-free read aligned genomes with high coverage (>40x) were selected: HG00096, HG00268, HG00419, HG00759, HG01051, HG01112, HG01500, HG01565, HG01583, HG01595, HG01879, HG02568, HG02922, HG03006, HG03052, HG03642 and HG03742. PCR-free seuencing data is not hampered by sequencing errors due to artificial in vitro PCR-induced TR expansion.

From the long-read test set we selected a sample of 10, 000 loci each corresponding to static TRs (stTRs) and VNTRs to create a labeled pileup dataset. Furthermore a random selection of similar regions was taken which acts as a control set. This was done by taking stTR coordinates and apply translation by a random integer between 1, 000 and 10, 000 with a random sign. To select read data for regions of interest, the Samtools application by Li *et al.*, 2009 was used. For detailed analysis of local read pileup, the Python module Pysam was used, which is a wrapper around the Samtools package.

#### Pileup Preprocessing

The Pysam data represents the number of reads that show any overlap in their alignment with a genomic nucleotide position for every TR in each human genome. On this data, a few data cleaning and stratification steps are applied. First off, TRs shorter than 30 nucleotides are omitted for a sufficient pileup analysis resolution. Secondly, TR pileup with a number of positions that did not match the nucleotide range of the TR coordinates were considered faulty data and omitted. Finally, TRs with positions that showed very low coverage (<10) on any one position were deemed too unreliable and are left out.

#### TR pileup profiles

TR pileups are made into comparable TR pileup profiles (TRPPs) by a set of normalization steps. First, the raw pileup data was normalized by the median coverage of its genome sample (division by read depth). Next, to compare TRs of different length, TR coordinates are normalized by min-max scaling and then divided into 29 bins. The pileup value in each bin is assigned the average value over coordinates that fall within it. In further steps, a two-way approach is used: 1) absolute pileup profile (TPP) which is unchanged and 2) a relative pileup profile (RPP) which captures any signature profile shape along the TR. An RPP is created by normalizing the sum of read counts across the TR profile to 1.

To detect any different outcomes of readmapping given TR length we split TR pileup into subgroups by sequence length: this was based on their relation with the read size of 250 nucleotides and the median fragment length of 500 from the frequency distribution of paired read end to end distance on the reference genome. Small reads are of nucleotide length (*l* < 250), medium reads (250 ≤ *l* ≤ 500) and large reads (*l* > 500), with read length *l*. These were expected to have a strong influence on pileup profiles: fragment and read length determine at what distance from the side of the TRs there is still a reliable mapping possible using unique flanking regions. Because the TR coordinate range is normalized to 1 for binning, this information is initially lost but TR length stratification aims to regain insight in read and fragment length related patterns.

Summarized TRPPs are created by integrating the TRPPs over every human sample, creating an 1) average TRPP, 2) spread TRPP, 3) variance TRPP, 4) Median Absolute Difference (MAD) TRPP, across the 17 human samples (for each TR seperately). Study of summarized TRPPs entailed comparison between each of the three TR types (static, variable length and control) stratified on TR length and count normalization.

#### TR profile analysis

TRPPs were hierarchically clustered using standard euclidean distance between TRPPs and, before clustering, Z-score normalization was applied for each bin across TRPPs. Finally, for analysis of total pileup, the average sum of pileup of TRs over the genomes, its variance, MAD and spread were calculated over absolute TRPPs.

#### Evaluation

We expect VNTRs to have similar TRPPs. To test this we evaluate whether TRPPs of VNTRs cluster together based on their pileup profile. As our ground truth is underannotated for 45%, we also expect that the predicted False Positives (FP’s, i.e. positives according to the VNTR predictor, but not annotated in our intial annotation set) for a large part will also be true VNTRs (TP’s). Consequently, we expect the TRPPs of FP VNTRs to cluster together with the TRPPs of TP VNTRs. Finally, in an effort to formulate a simple VNTR indicator from TRPPs, a set of binary variables is derived from variance, MAD and spread TRPPs. We expected that TRPP’s that show high variability in a sufficient number bins are correlated with VNTRs. First, for each TRPP type, a threshold is defined for the bin value and a second bin threshold represents the fraction of bins that need to pass the value threshold. Settings for the two thresholds were briefly explored over a range of values searching for a high correlation with classifier predicted VNTR probability. This correlation is quantified by the maximum absolute coefficient of Pearson’s *r* correlation. To relate total pileup statistics, we visualized the relation between predicted probability and total pileup and its variation and calculated Pearson’s *r* correlation.

## 3 Results

### 3.1 Overview

We are interested predicting whether a TR is variably expanding (a VNTR) or not (an stTR) based on information hidden in the genomic sequence. Hereto, we build a VNTR predictor based on features derived from the genome sequence of the TR using a training set of observed VNTRs in 4 haplotypes using long-read sequencing data able to span the VNTRs (see Materials). We expect to predict that TRs not observed as VNTRs to be variable in a larger population, i.e. false postives in the training are expected to be unlabeled true positives. We validate these predictions on a set of 17 other human samples being measured with short-read sequencing data. As the short-read data is not able to span the TR, we inspect whether the read pileup profiles of predicted VNTRs are indeed similar to observed VNTRs in the long-read data.

#### Training data

To train our VNTR predictor we make use of structural variations called from comparing 5 human haplotypes: 2 haplotype human data, one diploid human genome, and the reference genome (see Methods). Structural variations are annotated using Tandem Repeat Finder, resulting in 610.685 TRs (see Methods). TRs are annotated as being variable if the length of the TR varies more than 50 nucleotides between the 5 haplotypes. This results in 11.873 VNTRs, 1.9% of the total TRs. This serves as our training data for the VNTR predictor. According a recent paper looking at 17 long-read based haplotypes, we have an underestimate of 45% of VNTRs. We validate the predicted VNTRs using short-read sequencing data from 17 human samples of the 1000 genomes project that are not PCR amplified (see Methods).

#### Initial experiments for choosing feature representation

We use a Random Forrest Classifier(RFC) to predict VNTRs (see Methods). Before training this VNTR predictor, we first need to find an appropriate feature representation of TRs derived from the genomic sequence of the TR. We use a set of derived features based on their perceived predictive power. One of the features predicts the secondary structure (looping) of the sequence (which is one of the guiding principles on how TRs are being created). This feature can, however, not been calculated for 25% of the TRs due to the instability filter enforced by the Mfold program. Mfold will omit any stability calculations if it leads to an isolated base pair: matched base pairs that are neighbored by two mismatched base pairs which is deemed extremely unstable (Zuker, 2003). Indeed, all of inspected TRs with no returned mfold value were short (30 bases) TRs of mono- or di-nucleotide repeats with non-complimentary base types which would cause isolated base pairs. We initially experimented with three different ways to cope with this missing information, either by 1) zero-imputation, 2) dropping TRs, and 3) including a feature that indicates mfold missingness (see Methods). Secondly, two methods were tested to integrate categorical features with continuous features: 1) integer encoding and 2)one-hot encoding. For this analysis we compared the performance of 100 tree RFC performances on the VNTR training data.

The best data representation appeared to be omitting any samples with undefined mfold value and using a one-hot encoding of categorical variables (see Figure 1). However, due to the increased feature dimensionality from one-hot encoding, training time rose significantly. Since there was only very little difference in performance, we chose integer replacement over one-hot encoding. This also simplified relating results to the underlying categorical variable.

**Fig. 1:**
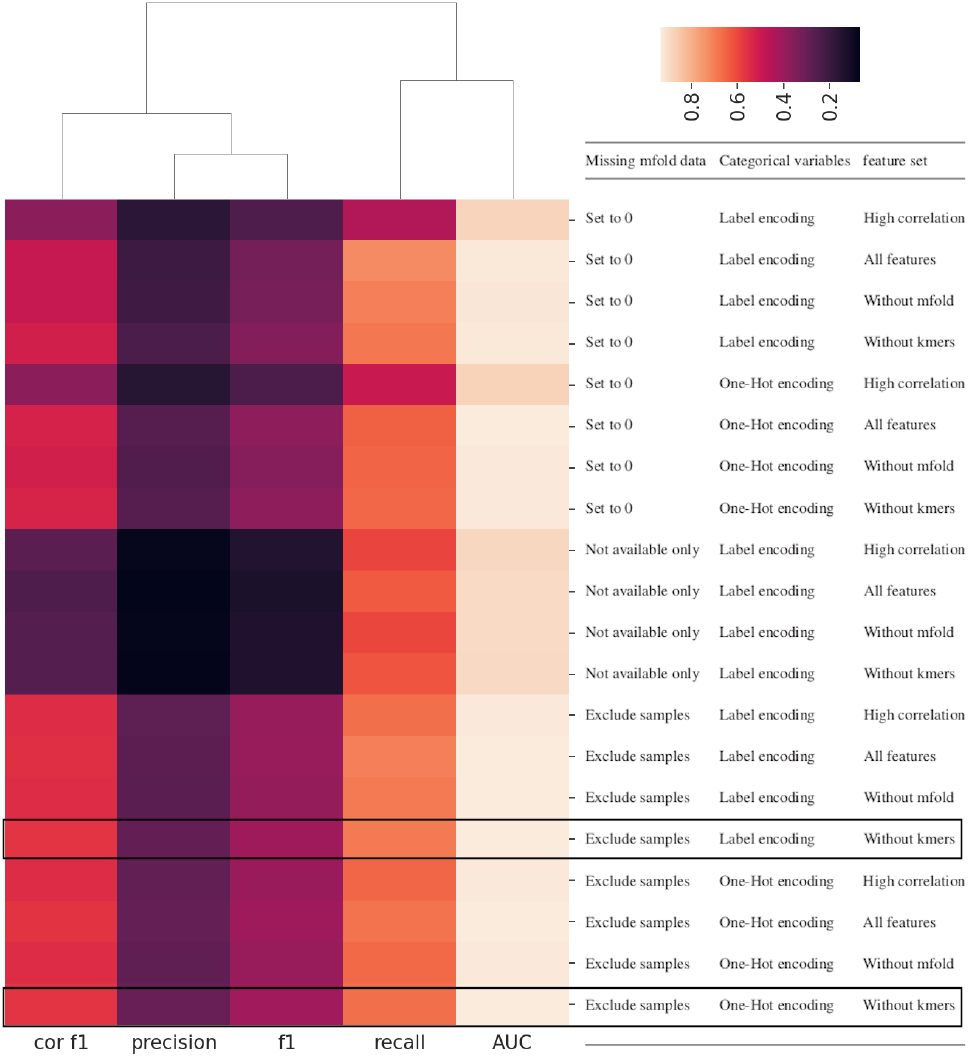
Classification performance(columns) on an RFC containing 100 trees trained with each dataset (rows) generated by a combination of different feature sets, categorical data handling and handling missing data. Mfold data was missing for 25% of TR samples, which can be handled by 1) zero-imputation or 2) excluding TRs. Also included is the performance for only using the missing samples (while omitting the mfold feature). Categorical data was combined with continuous data by 1) Label encoding (integer replacement) and 2) One-hot encoding. A small number of feature sets was tested: 1) High correlation features, which is a selection of the 7 features with the highest absolute VNTR correlation (see Figure 2). The best corrected F1 performance is achieved by excluding samples with missing mfold data, one-hot encoding for categorical features and using all features except k-mers. The selected method uses Label encoding instead one-hot encoding, for convenience reasons.

Omitting TRs with missing mfold resulted in a data set of 435, 481 TRs with reasonably unaffected VNTR ratio of 2.44%. The strongest corrected F1 performance was achieved by training on all features except k-mers, which is consequently left out of the feature set for the final classifier.

Furthermore, when splitting TRs in STRs and long TRs categories(Methods), some difference in performance could be observed. VNTR annotation seemed to be less prevalent for STRs (3, 872 of 306, 096, 1.26%) and more for long TRs(6, 774 of 129, 385.00, 5.24%). Moreover, over all TR types, the classifier performance was highest in long TRs and lowest in STRs and average for generic TRs (see Figure S3). In further analysis we focus on the generic TR dataset, to pursue a generalized application for novel VNTR discovery.

### 3.2 Statistical analysis

Figure 2 shows the correlation between each feature. Sequence length shows a high negative correlation with mfold (positive with stability of auto-folding). Features related to TR length show a small positive correlation with CG-content. Many features show a clear (non-linear) interaction in pairwise scatter or density plots as is seen in Figure 3 for nucleotide composition features and centromeric distance. Although there is a clear dominance of the stTRs in almost all of feature space, there are local hotspots for VNTRs in some of the two dimensional feature spaces. There is increased VNTR ratio in CG rich sequences in regions far from the centromere. Furthermore, separate C and G percentages do not show as strong a local increased VNTR ratio when paired with centromeric distance. In the plot relating A and G percentages, it appears that extremes in A percentage show a low VNTR ratio. There is a single high VNTR ratio region around 50% A coupled with low G percentage. A roughly inverse pattern is observed for A and T percentages, where equal percentages for A and T seem to be related with higher VNTR ratio. See Figure S1 for other feature-feature plots.

**Fig. 2:**
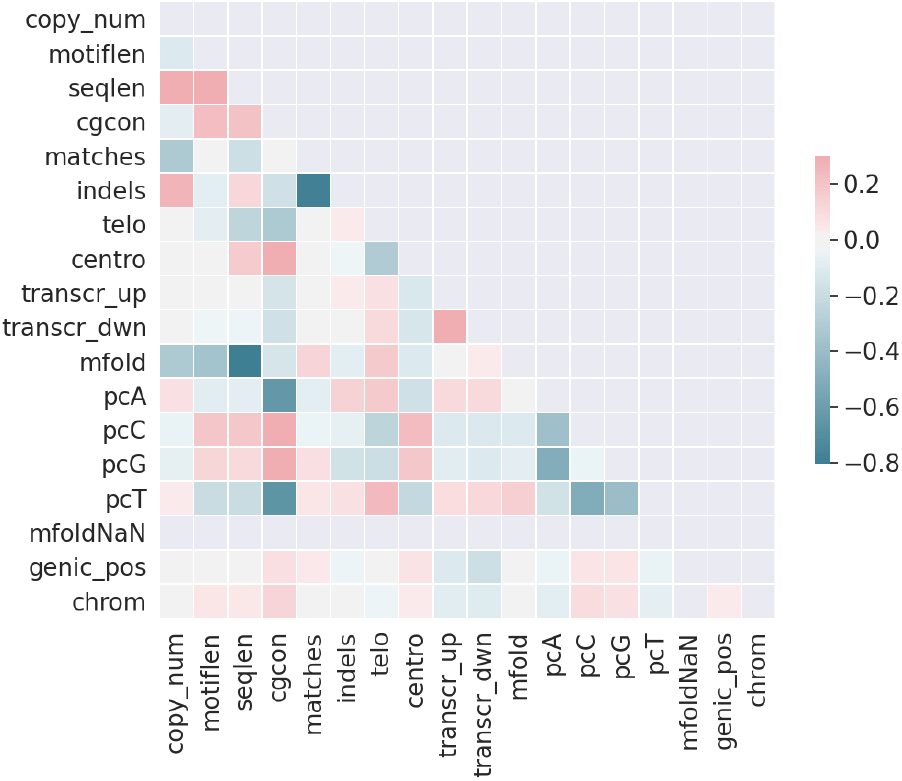
Pearson’s *r* correlations among features. Strong negative correlation exists between (negative) mfold free energy change and sequence length. MfoldNaN is constant because all NaN values are removed from the dataset.

**Fig. 3:**
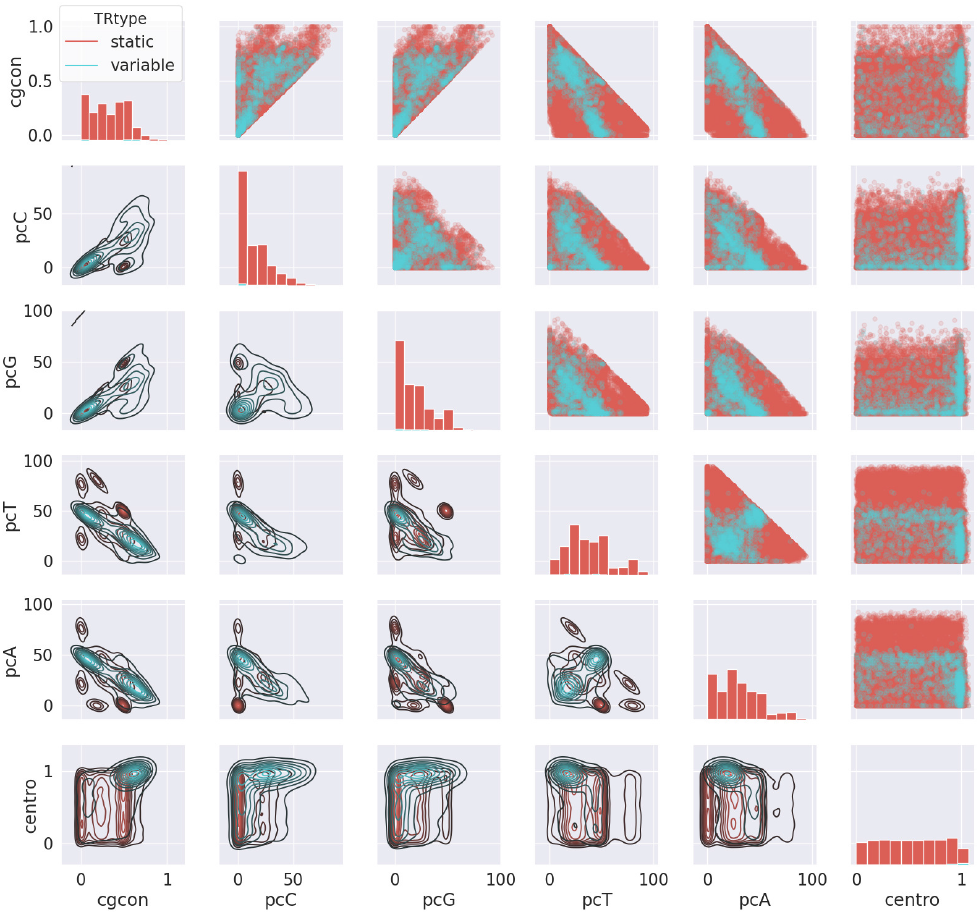
Feature interactions for nucleotide composition features and centromeric distance. Kernel density plot (lower triangle) is performed separately on each class with a stratified sample of 25 000 TRs. Diagonals plots show the density distribution with where VNTR dominates stTRs (indiscernable) in all of single feature space. The upper triangle depicts the simple semi transparent scatter plots, with VNTR cases plotted in the forefront.

Correlations of the features with VNTRs appeared to be different for the STR/long TR categories and handling NaN mfold data (Figure 4). For example, the correlations in long TRs are stronger than in STRs for mfold, matches, indels and weaker in copynumber and G nucleotide ratio. For the chosen data representation, there is a VNTR correlation with seq_len (0.42, *p*<0.01)and an opposite correlation (−0.39, *p* <0.01) with mfold, in line with the negative correlation between these two features. There is also a low negative correlation found for telomeric and an opposite positive centromeric distance (−0.25 and 0.25 respectively, *p*<0.01). To clarify, this equates to a weak positive VNTR association in regions close to telomeres. The strongest k-mer correlations are purified for either the pair of A and T nucleotides or otherwise the pair of C and G nucleotides. These k-mers generally show weaker correlations than the nucleotide composition features (CG ratio and pcA,pcT,pcG,pcC). There are two exceptions to this: 1) a weak negative VNTR correlation for the AAA k-mer, in contrast to almost no correlation for pcA and 2) a full GGG k-mer that surpasses the pcG slightly in VNTR correlation.

**Fig. 4:**
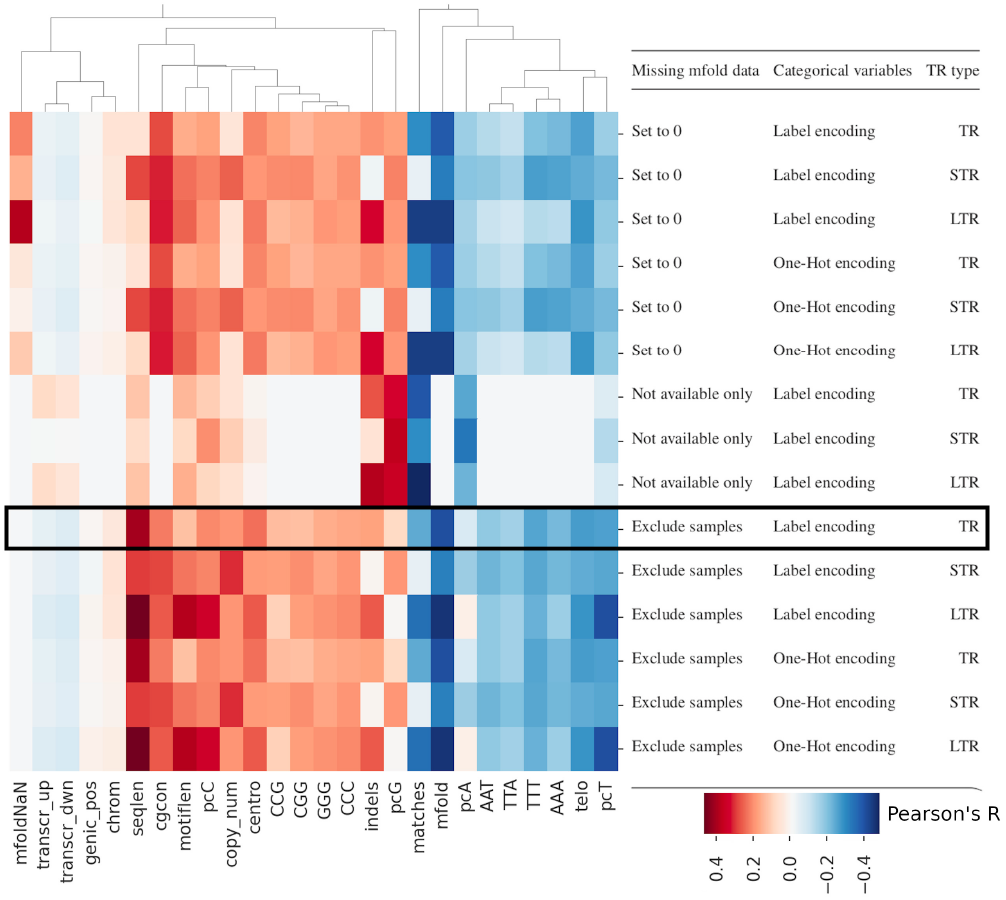
Pearson’s *r* correlations with VNTR annotation of individual features(columns), for different combinations of data representation and TR categories (rows). Mfold data was missing for 25% of TR samples, which can be handled by 1) zero-imputation or 2) excluding TRs. Also included is the feature-VNTR correlation for only the missing samples (while omitting the mfold feature). Categorical data was combined with continuous data by 1) Label encoding (integer replacement) and 2) One-hot encoding. The variant chosen in the final classifier is outlined. To represent k-mers, only the top 4 positive and negative k-mer correlations are visualized. The strongests k-mer correlations do not mix A,T with C,G nucleotides and do not significantly exceed their respective nucleotide ratio features (CG ratio and pcA,pcT,pcG,pcC).

### 3.3 Sequence classifier

The final classifier showed a test performance of 0.62 for precision, 0.25 for recall, and 0.36 for F1 with a high ROC AUC of 0.94. As our training data is underannotated with 45%, implying that the expected number of VNTR should be larger than is annotated in the training set (Methods), we also calculated corrected performances assuming that there is an equal (prior) chance for a TR to be actually labeled VNTR (Methods). These corrected performances are: 0.78 corrected precision and 0.38 for corrected F1. The high 0.94 AUC of ROC is not very descriptive of imbalanced binary classifier performance. A more suitable description is the Precision-Recall Curve (see Figure 5) which is comparable to the F1-score under different probability thresholds. The AUC of the Precision-Recall curve was 0.45.

**Fig. 5:**
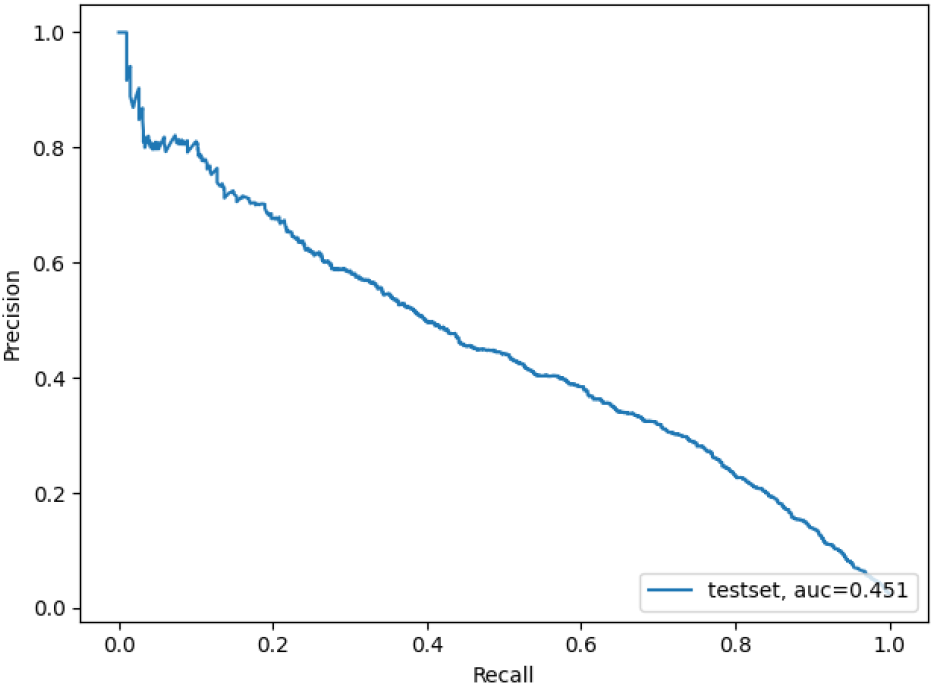
Precision-Recall curve of the final VNTR classifier on 10% test data with the corresponding area under the curve.

The reported feature importances (Table 1) are calculated by the ratio of splits on that feature in all of the trees in the trained RF classifier. The most used features are sequence length, mfold, copy number and motif length. Then follow, with similar usage, CG-content, centromeric distance, match percentage, nucleotide ratios and distance to nearest transcription start site. The lowest usage is seen for indels, chromosome, genetic functional region. In our 100 tree RFC exploration phase, the RFC trained on the chosen dataset but using all features including k-mers also showed lower feature importance for all k-mers than any nucleotide composition features, even ranking below the genomic position.

The final classifier predicts 1.36% VNTRs on the complete dataset and 0.99% VNTRs on the test set, which are both lower than the VNTR ratio in the annotated set which is 2.44%. On the total set, a total of 915 unannotated candidates are generated as FP cases. In the test set these are predicted as VNTR while not annotated like that in long read data. Applying the classifier on the test set generated 163 unannotated VNTR predictions. Since the precision is quite high, we expect that these unannotated VNTRs are actually true VNTRs.

### 3.4 Validating predicted VNTRs

Next, we set out to find evidence for the predicted VNTRs to be true VNTRs. For that we analyzed short-read data across 17 human samples. The short-read data cannot span the TR, but we expect that the read-mapping profile across the reference genome-based TR (denoted as TR pileup profile, or TRPP) is uniquely different between VNTRs and stTRs. Moreover, we expect that there is a varying percentage of reads mapped to a VNTR when there are expansions within the 17 human samples as compared no varying percentage for stTRs.

#### Single genome

For each TR we created a TR read-mapping profile (TRPP) that shows the count of mapped reads over the TR region. To be able to compare the variable length TR regions, we binned the TRPPs into 29 bins (Methods). TRPPs vary considerably within the between TR category (see Figure S8 for some examples) but also within the same TR category. Therefore, Figures 6–8 show the average TRPPs across all VNTRs and across all stTRs for three different categories of TRs; small, middle and large TRs (Methods). It clearly shows that these average TRPPs are different for both VNTRs and stTRs over the different categories.

**Fig. 6:**
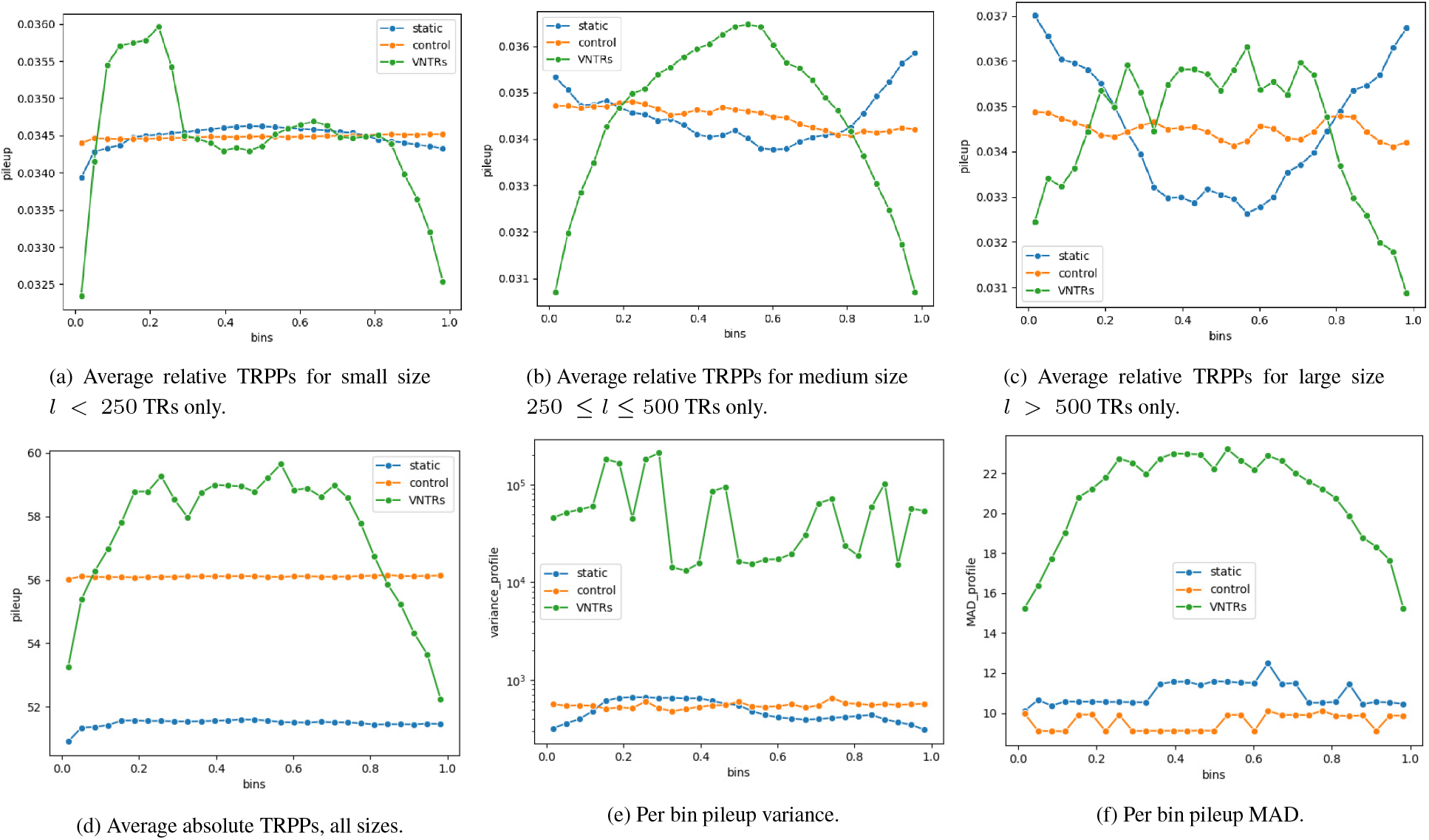
Pileup profiles and derived pileup profiles compared for different TR categories and control regions in the single genome HG00096.

To study the effect of the tandem repeat pattern, we compared these TRPPs with pileup profiles of randomly selected parts of the genome in the neighborhood of a tandem repeat (ensuring a similar genomic context). Those control regions are assumed not to be tandem repeat regions and reads can thus uniquely match in contrast to TR regions. The resulting control pileup profiles (denotes as ctlPPs) are more stable, as can be seen in Figures 6–8.

Looking into more detail, we see that TRPPs of VNTRs fall off sharply near the borders of the TR region. In contrast, average stTRPPs show a pattern with increased pileup near the borders of TR regions. Short TRs show different TRPPs than the medium and long TRs for both stTRs and VNTRs (Figure 6a-c). Short stTRPPs are somewhat constant and fall off slightly to the borders. TRPPs of short VNTRs include an additional dip in the middle region with an asymmetric sharp rise at the 20% region.

When averaging TRPPs over all three TR categories (Figure 6d) we do not see any difference in profile shape between an stTR and a control region. Although the control group does show a higher absolute pileup (explainable by the mappability difference between unique and repetitive sequences), VNTRs have the highest amount of total pileup, but the level drops below the level of the control group pileup at the borders, outside of the 10% to 80% range.

TRPP variances and Mean Absolute Difference (MAD) per bin across TRs are shown for the static and variable TRs as well as the control regions in Figures 6e,f, respectively. On average there is a clear increased variability over the complete length of VNTRs, showing volatile levels between bins. The stTRs show an order of magnitude in variability that is similar to ctlPPs (Figure 6e). The stTRPPs show lower variability near the borders and only show higher than control MAD between the 15% to 50% marks. The MAD values shows a less outlier sensitive profile of bin pileup variability (Figure 6f). Profile shapes of MAD variability closely follow the average absolute TRPPs (in Figure 6d). Values are of similar order of magnitude but the VNTR variability is clearly larger over the complete TR range. Even though stTRPPs show lower average pileup values than ctlPPs, stTRPP MAD values are slightly increased with respect to ctlPPs. The MAD at each bin in ctlPPs surpasses any average differences in the absolute pileup between the TR types in Figure 6d.

To investigate individial TRPPs in more depth, we next clustered TRPPs using a 200 randomly chosen regions from each TR category and control regions (all sizes). We find a clear cluster of TRPPs belonging to VNTRs (cluster marked *V*_*a*_ in Figure 7) and TRPPs from TRs mixed with ctlPPs (cluster *S*_*a*_ in Figure 7). Most notable in VNTRPPs compared to ctlPPs or stTRPPs is the increased extreme values, both locally within the profile and compared between profiles.

**Fig. 7:**
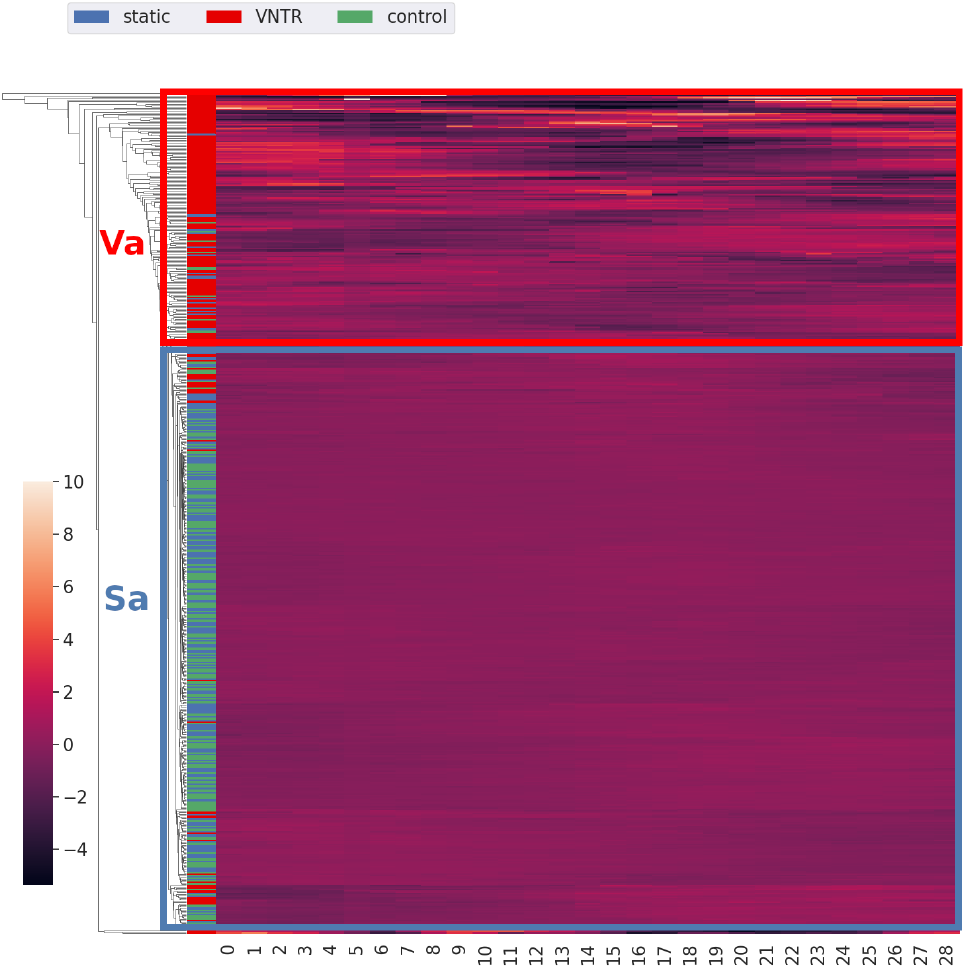
Clustering of individual relative stTRPPs, VNTRPPs and ctlPPs in the single genome HG00096. Rows are pileup profiles and columns are one of one of 29 bins. Z-score normalization is applied to columns.

#### Inter-genomic

Next we set out to investigate the variability of the TRPPs across the human samples in more detail. For that we looked into the variability of the TRPPs across the different samples for 400 randomly selected TRs evenly distributed over stTRs and VNTRs. Furthermore, each TR category has equal representation of agreement and disagreement in labeling between the sequence classifier and annotation. This is equivalent of 200 random selections from each classifier predicted TP,FP,TN, FN truthgroup. We then clustered the variance-based TRPPs and found 3 clusters, see Figure 8a. For each TR, we compared the original annotation and predicted TR type, and looked whether saw that several clusterings showed a clear separation of TR types that both labelings agreed on. The best clustering (in the sense of enrichment with respect to TR category) was achieved by clustering based on the variance of the relative TRPPs. (Figure 8a). It shows a clearly separated subcluster with stTR label agreement for cluster *S*_*a*_ with a 77% agreement ratio. This region does contain sporadic VNTRs as indicated by the annotations but consists almost fully of stTRs based on the classifier output. In contrast, the other two clusters (*V*_*a*_,*V*_*b*_) show agreement mostly for predicted VNTRs labeling although there is less agreement here with 36% and 12% agreement respectively. For these clusters, when the annotation indicates an stTR, the classifier predicts a VNTR indicating a false positive. These two clusters are almost completely labeled VNTR by either one or the other datasource.

**Fig. 8:**
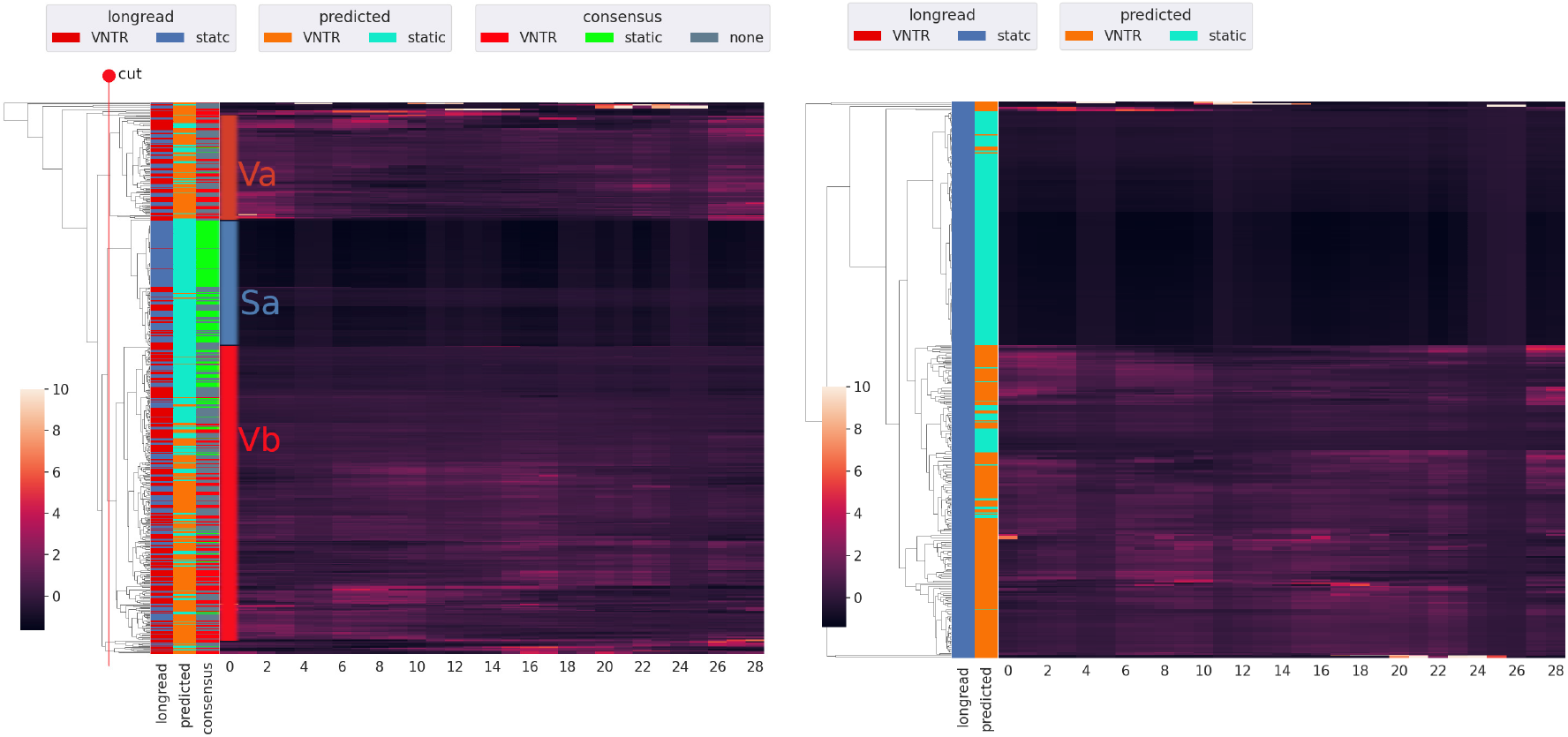
Clustering of per bin inter-genomic variance of relative TRPPs. Row labels show in left to right column order: training annotations (red=VNTR,blue=stTR), classifier predicted labels (orange=VNTR,magenta=stTR), 8a only: label indicating whether there is consensus between annotaton and prediction (red=consensus VNTR, green=consensus stTR, blue=no consensus). Rows are variance TRPPs and columns represent TR positions (across 29 bins) and are Z-score normalized. (a) Clustering all. Rows are 200 randomly selected TRs for each combination of predicted label and long-read annotated label(TP,FP,TN, FN). Cluster labelings *V*_*a*_,*V*_*b*_ and *S*_*a*_ result from the cutoff position as shown. Both the annotations as well as the prediction by the classifier show that *V*_*a*_ and *V*_*b*_ contain predominantly VNTR and *S*_*a*_ almost exclusively stTRs. (b) Clustering only TRPPs with stTR annotation. Rows are 200 randomly selected TRs for each combination of predicted label and TRs that are long-read annotated as stTR (FP and TN predictions).

The three clusters are assigning a label according to majority vote the annotation labeling (which coincides with majority vote of agreed labeling) so that the clusters *V*_*a*_ and *V*_*b*_ are VNTR clusters and *S*_*a*_ an stTR cluster. Label counts can be reviewed in Table S1. Even though disagreement is increased for both these VNTR clusters *V*_*a*_ and *V*_*b*_, they show a high FP ratio of 0.430 and 0.283 compared to 0.017 in the stTR group. In contrast, FN cases only show an increase to 0.212 and 0.284 in *V*_*a*_ and *V*_*b*_ compared to 0.209 in the stTR group. Clustering only the annotated stTRs, we can further observe the difference between agreed and disagreed static cases of Figure 8a in Figure 7b. Static annotated TRs that the classifier predicts as VNTRs are for the majority clustered. A close second best label agreement in subclusters was achieved using normalized individual profiles from only the single genome HG00096. Clustering variants can be found in Figures S9–32. Inspecting predicted probability of clustered TRPPs showed continuous behaviour in neighboring profiles (Figures S10,12,14,16,18,20,22,24,26,28,30,32). Variants with per TR pileup normalization showed better long-read label separation than raw data counterparts.

#### TRPP variability indicators

We next investigated whether a single variability measure across TRPPs can summarize their differences. We tested if they correlate with the predicted probabilities for being a VNTR as produced by the classifier (Methods). To summarize the variability of the TRPP we defined binary (high or low) indicator variables that require a certain number of bins to exceed a variability threshold in variance-based TRPPs (Methods). We found three moderate correlations with classifier predicted probability with *p* < 0.01 for TRPP variability indicators, after testing different thresholds (see Figure 9: Profiles with any bin showing a variance above 0.09 show a 0.47 Pearson’s *r* correlation with classifier predicted probability. Spread shows a similar result for at least one bin above 0.9 with a correlation of 0.40. MAD shows a negative correlation of −0.23 with predicted probability if all bins are above 0.6.

**Fig. 9:**
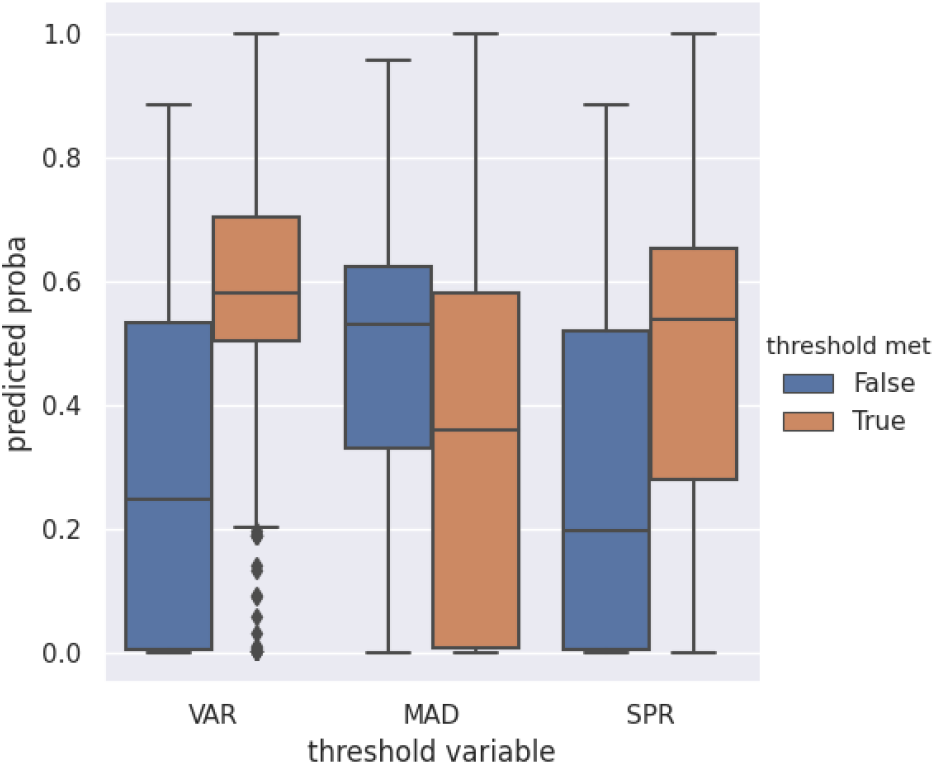
Boxplot of classifier predicted probability against several pileup binary threshold variables. Per bin thresholds and number of bins thresholds: variance 0.09 for 1 bin; MAD: 0.6 for all 29 bins; spread: 0.9 for 1 bin.

#### Total pileup indicator variables

Finally, we explored whether the total count of reads mapped or the variability of that number across TRPPs correlated with predicted VNTR probabilities from the classifier. We tested average total pileup and variance, MAD and spread of total pileup. These showed negligible correlations with classifier predicted probability (Figures S33-40).

## 4 Discussion

### 4.1 Classifier

The Random Forest Classifier (RFC) managed to predict VNTRs with a weak F1 performance of 0.36 and a precision of 0.62 with a recall of 0.25. The AUC of the Precision-Recall curve (Figure 5) reaches a value of 0.45, with a precision that drops sharply followed by a short plateau as recall grows from 0 to 0.10 after which precision drops linearly. For the chosen threshold, the classifier sacrifices in the number of VNTR candidates produced for a higher reliability.

The classifier uses only a few features extensively in deciding the TR class (Table 1). There is a clear priority for sequence length related features where total sequence length is used the most. Even though the combination of copynumber and motif length combined gives the same information, sequence length itself is preferred. Still, copynumber and motif length are more used than expected from their individual correlation with VNTR status, suggesting that their interaction is important or that these are VNTR related in discontinuous fashion.

Mfold is the second most used feature, which may be caused by its high negative correlation with sequence length. A high mfold (close to 0 energy change) could effectively be used as a proxy variable for detecting low sequence length. However, analyzing the interaction plot, we see that there is sufficient unique information for VNTR separation to warrant the inclusion of folding as feature for the VNTR classifier (see Figure S2). A special subgroup appears in long TRs with around 0 folding stability that show a relatively high VNTR ratio. The rest of mfold values clearly show a correlation with the length, where the length determines the maximum amount of negative free energy change. When TR cases lie outside of the this main trend, they appear enriched for VNTRs. This is also reflected in a stronger divergence of classifier predicted VNTR probabilities for mfold values lower than −200 (see Figure S7). Finally, since features representing nucleotide composition are less important (Table 11), it is unlikely that the mfold feature is entirely determined by it. The importance of mfold is also illustrated by a (slight) decrease in classifier performance when omitting the mfold feature (Figure 1).

Other features all show a feature importance of 5% or lower. Most notably, the gene element type feature, here represented by a integer encoded ordinal variable for each type of genomic context, shows almost no usage. This was also the case when using a categorical one-hot encoded data representation, where each of the binary variables represents whether it belongs to a certain group of gene elements or not. Each of these markers showed similarly low importance.

Telomeric regions have been shown in literature to be enriched both for TRs as well as VNTRs, with varying reports of their ratio (Audano *et al.*, 2019; Näslund *et al.*, 2005). Here, we find a weak 0.25 positive correlation for VNTR status of TRs with proximity to telomeres. Telomeric distance correlation is properly negated in VNTR correlation coefficient for proximity to centromeric regions. Furthermore, the impact of telomeric distance consists of a highly localized probability increase close to the telomeres. The increased predicted probability quickly drops to a constant beyond the first 5% of the chromatide arm (see Figures S5,6).

K-mers have not been included in the final dataset, but if used, the performance dropped slightly. Moreover, using that feature set, the k-mers showed less usage than any other nucleotide based feature. Also when considering correlations with VNTR status, the most relevant k-mers are those that are predominantly single nucleotide, or separated in *{A, T}* only or *{C, G}* only k-mers, which would make them an indicator of the CG content feature. It seems, that at least for *k* = 3, specific nucleotide patterns represented in k-mer ratios do not hold strong additional information for VNTR prediction outside of representing the already present CG content and single nucleotide features.

From the feature interaction plots the most comprehensive conclusion is that VNTRs are enriched in CG rich telomeric regions. Furthermore, VNTRs rarely comprise of more than 60% just A or just T nucleotides, but do in some cases contain mostly C or mostly G nucleotides.

Mfold NaN cases were omitted by the classifier performance which leaves out 28.7% of the data. Therefore, it is important to note that our classifier and following pileup analysis represent only TRs with a defined Mfold value. From the exploratory experiments for variants of handling NaN data, we learned that the omitted Mfold cases proved to be harder to classify (Figure 1). Nevertheless, these omitted cases showed difference in VNTR correlations, where sequence length and CG content correlation is relatively weak while G counts show increase in correlation (Figure 4). We therefore leave these mfold omitted to be handled in a tailored way as future work. In general, we do conclude that there is sufficient reason to use multiple features in a classifier context over simple correlations or additive effects.

### 4.2 Profile analysis

#### Single genome

The general overview of average pileup profiles for TRs in a single genome (Figures 6a-c) showed a clear difference for VNTRs and static TRs (stTRs) versus control sequences. VNTRs on average show a clear drop in TR pileup near the borders of the TR, while stTRs show the opposite shape (except for stTRs below the read fragment length). A relative drop of VNTR pileup to the borders may be caused by a TR variant that is longer than the reference TR. This causes reads containing additional inner TR repeats to be mapped on a smaller space on the reference genome, reaching a higher pileup density. The outer regions of the TR are not suitable for mapping the surplus of reads. Reads near the flanks would require a good mapping on flanking regions as well, which would be different from the elongated inner TR region.

The average pileup pattern of stTRs which shows decreased relative pileup in the inner regions may represent the reduced mapping quality of reads. These reads may be lost to different similar regions outside of the TR or may even be unmapped due to a cutoff for minimum mapping quality. Mapping quality is expected to be low in repetitive regions, because of the large amount of mapping positions with similar match scoring. This is also reflected in average absolute pileup (Figure 6d), where VNTRs exceed control sequences in the center region and stTRs consistently show fewer reads mapped than control sequences.

There seems to be a large within-group between-TR variation in absolute pileup for both TR types as well as the control group (Figures 6e,f). Furthermore, the MAD variation exceeds the difference in average absolute pileup between the TRtypes. Therefore, it seems that a single TRPP can not be classified by maximum similarity to one of the average absolute TRPPs. This suggests that we need the multigenome TR comparison data to find distinguishing indicators. VNTR pileup Variance and MAD profiles themselves clearly exceed those of stTRs and control sequences, which are similar to eachother (Figures 6e,f). However, the VNTR MAD profile shows a shape that resembles the shape of the average absolute TRPPs, suggesting that there is a consistent ratio of deviation along the TR and the shape difference is a natural result from this. In contrast, the variance profile shows a highly irregular pattern that does not follow the average absolute TRPP. It is therefore expected that most information can be gained from comparing TRs between genomes on the basis of pileup variance, including its shape.

Although above results suggest that identifying VNTRs by pileup analysis from a single sample is difficult, a simple clustering of individual pileup profiles did show a clear separation into VNTR and stTR together with control sequence cases (Figure 7). This is interesting because there is no multi-genome comparison used to detect this VNTR similarity. Arguably, there is already an implicit comparative step in mapping reads from the sample to the reference genome, where a variant in TR length would cause detectable side-effects. However, the reference is expected to represent the dominant variant for each locus, which would cause the majority of VNTRs to be of equal length compared to the reference. Note that, because the provided long-read labels only detect the most variable VNTRs, there is a stronger than expected separation from single sample analysis. From the clustering it appears that there is a large within-profile variability of pileup for neighboring bins as well as more extreme values in VNTRs. Between VNTRs, the positions of local pileup highs and lows vary wildly which may underlie the relatively low information gain from multi genome average profiles per TR type in the following section.

#### Inter-genomic

From per TR inter-genomic comparison, the strongest separating clusterings were obtained for normalized pileup variance profiles. These clusterings were applied to a subsample consisting of equal parts of four truthgroups of the sequence classifier prediction. This resulted in worse long-read label separation than random subsamples but it shows promising results in linking long read annotation labels to classifier predicted labels. The clustering produces a strong separation of consensus stTRs. Furthermore, potentially unlabeled VNTR candidates in the form of FP cases are enriched in clusters with high VNTR consensus with an 18.8 times higher rate, whereas the FN is enriched slightly, with a rate increase by a factor of 1.3. Moreover, normalized variance profiles did well in separating FP cases from TN cases in a separate clustering of longread static annotated TRs (Figure 7b). This confirms the different nature of consensus stTRs versus these FP labeled TRs according to the pileup similarity. It reinforces the credibility of their status as unlabeled VNTR candidates.

Summarizing inter-genomic pileup data into single variables shows a moderate 0.47 correlation with VNTR probability as predicted by the classifier. It represents intergenomic pileup profiles that contain a variance above 0.09 in any of the 29 bins. This is a very low value compared to the average variance for TRs, suggesting it mostly separates extremely stable TRs from VNTRs. It must be noted that the thresholds for the variability and the number of bins were found by searching a grid of possible values and selected for the highest absolute correlation for three variables, which is inherently biased towards finding a correlation. The TR total pileup shows little correlation with VNTR probability as predicted by the classifier, suggesting again that there is little predictive value in absolute pileup values.

MAD clustering performed worse than variance clustering and single genome profiles (Supplementary Figures 8,16,24). It is worth noting that the single genome data with normalized individual profiles was a close second in separating consensus labels. It seems that relations between bins within a single TRPP, hold almost as much VNTR predictive information as comparing 17 genomes and combining them into a variability based TRPP. For all profile variants, a better separation in clusters was observed by applying TR profile normalization first, suggesting that profile shape holds more information than its magnitude.

### 4.3 Suggestions

To improve the classification performance, it would be interesting to focus more on gene related properties. The genomic related features in this article seemed to be of relatively low value for the classifier, despite previous results showing clear evolutionary pressure against expansion in gene related regions at least for STRs (Willems *et al.*, 2014). Furthermore, the classifiers low feature usage of k-mer features suggests a weak role for short base pair patterns. A different approach of capturing gene related information may be required, such as involving gene function, RNA expression, conservation and epigenetics factors.

Although the classifier only performed slightly better by omitting NaN mfold samples than imputing with zero values, the nature of NaN mfold status requires more close inspection. For now we accept this choice because of its low correlation with expansion, although for future application a separate classifier needs to be build to handle this subset of data to be able to predict every TR. We decided to handle STRs and long TRs together in a single classifier, but the exploratory classifiers showed that long TRs are easier to classify than STRs. For long TRs compared to STRs, the correlation of features with VNTR status showed no extreme differences outside of the number of mismatches, which loses all of its negative VNTR correlation in long TRs. It seems that if a TR with a short motif has some relative amount of mismatch, this is correlated with stability while long motifs with the same percentage of mismatch are not. In conclusion, there is some reason to separate the analyses of STRs and long TRs to gain deeper insight in their differences. However, as long as motif length is present as a feature, there is no need to build a separate classifier.

The strength of our analysis would be greatly enhanced by expanding the ground truth data of long-read labeling. Simply using more samples would increase the coverage of VNTR labeling. In the current state, due to the unlabeled VNTRs, we expect that actual performance of the classifier in precision is higher and in recall is lower than reported here. More complete VNTR labeling would in the first place enhance precision and prevent learning from false information, but it will also benefit the recall by the possibility of setting a lower VNTR decision threshold, which balances recall and precision performance.

To increase the resolution of the inter-genomic pileup comparison, future studies could allow data from non-PCR-free short-read sequencing samples. There are many more samples available without this restriction. This would decrease the reliability of TR copynumber in short-read data in favor of a more stable statistic, as well as a higher chance to capture rare copynumber variations of VNTRs.

Despite testing several statistical properties on pileup variability of each TR over multiple genomes, it still appeared that single genome data showed similarly strong VNTR separation with 17 times less information. Apparently, the sheer variability of VNTR pileup patterns does not easily summarize into descriptive single indicator values. The most promising future direction is therefore either in investigating the sources of the variability itself or single genome profiles. For the latter, what has not been tried is to summarize individual genome profiles into a single value. Therefore, it would be interesting to consider single TR within pileup fluctuation of neighboring positions as a feature. Since the pileup clustering already showed some promise in exploratory VNTR separation in an unsupervised manner, the next step would be to make this more concrete with a supervised classifier based on pileup profiles. A convolutional neural network can process local information from pileup profiles in one dimension, over the different genome samples in a second dimension, to decide the TR type. This would integrate any comparison within TR and between samples. Since both approaches show some consensus, it is a sensible step to combine them into a single classifier. If a strong classification performance is achieved in detecting VNTRs among stTRs already using single genomes, a first move can be made towards VNTR copynumber detection. One could shift labeling from VNTR versus stTRs towards the case of copynumber growth or shrinkage compared to the reference genome.

## 5 Conclusion

We have shown that a random forest classifier is able to predict the variable expansion of a tandem repeat region, as given by a longread based labeling of regions across 5 haplotypes. These predictions have been validated by variances in the pileup profile of short-read data mapped to the TR regions across 17 genomes. We have shown that the Mfold predicted free energy change of self-folding proved to be one of the most useful features for detecting VNTRs. Furthermore, we have shown a positive correlation for VNTR probability as predicted by the classifier for the following condition on variance pileup profiles: any one region inside the profile needs to show increased pileup variance over the 17 genomes. Finally, pileup information from single genomes seemed to be nearly as descriptive of VNTRs as pileup variance profiles over the 17 genomes.

## Acknowledgements

I would like to thank Marcel Reinders for an incredible amount of patience, kindness, and support, even in times that I may not have deserved it. Thank you Jasper for a great introduction into the world of tandem repeats and setting up for this research. My family, Marieke, Carel and Wouter, for believing in me and pushing me on. Sacha, for finding time to look at my work once again in turbulent times. Last but not least, I thank AnneMijn who has come into my world during this research, helped me in countless ways and made me a better person.

## Notes

### Competing Interest Statement

The authors have declared no competing interest.

