## Supplementary material for "VNTR prediction on sequence characteristics using long-read annotation and validation by short-read pileup"

**Diederik Cames van Batenburg [1,*], Alexander Gultyaev [1,*] and Marcel Reinders [2]**

*To whom correspondence should be addressed.

### This PDF file includes:

Figs. S1 to S41
Table S1

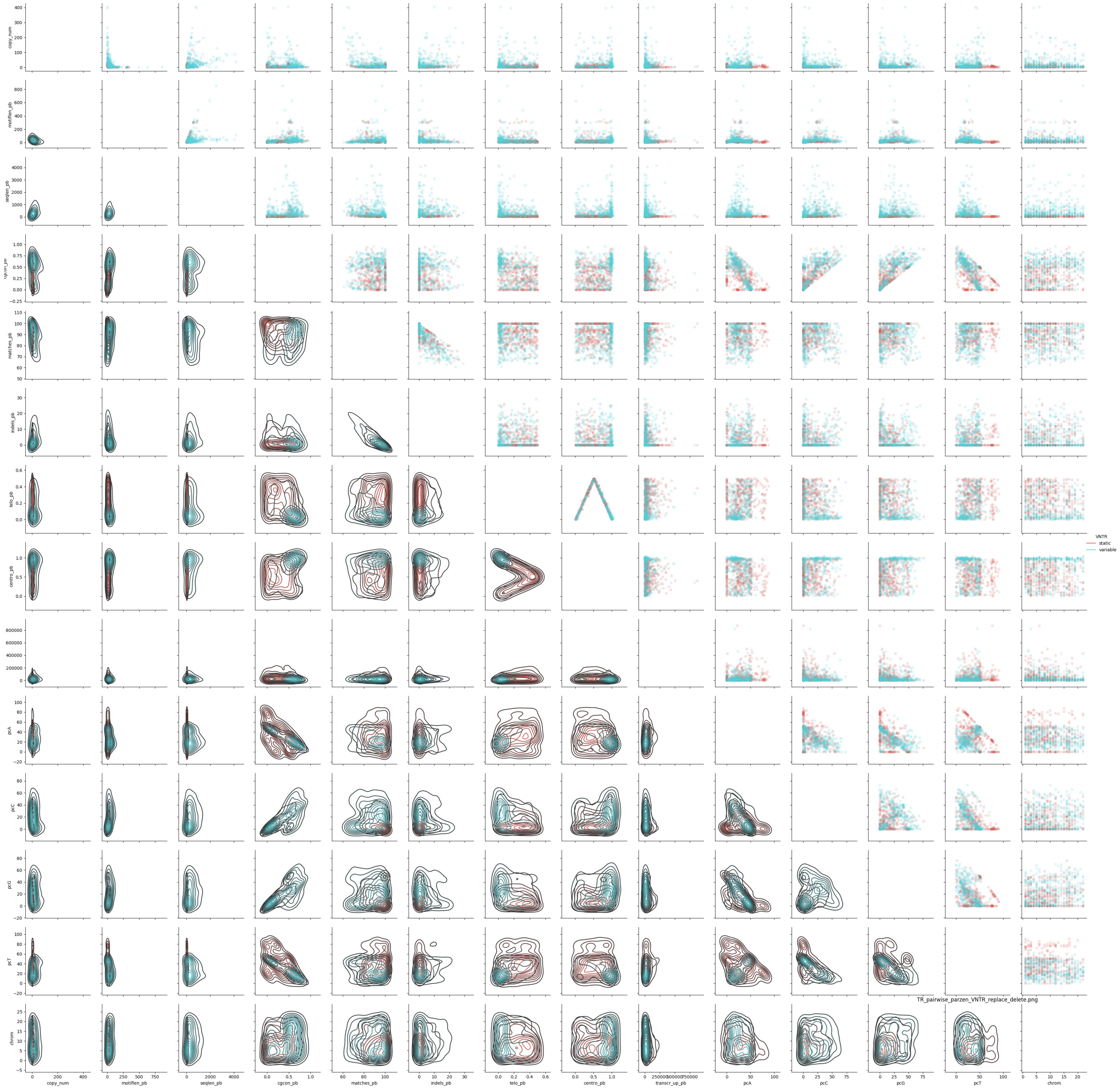

**Fig. S1.** Pairwise feature-feature scatterplots on the upper triangle and kernel density plots on the lower triangle. Shows only numerical and non-kmer features.

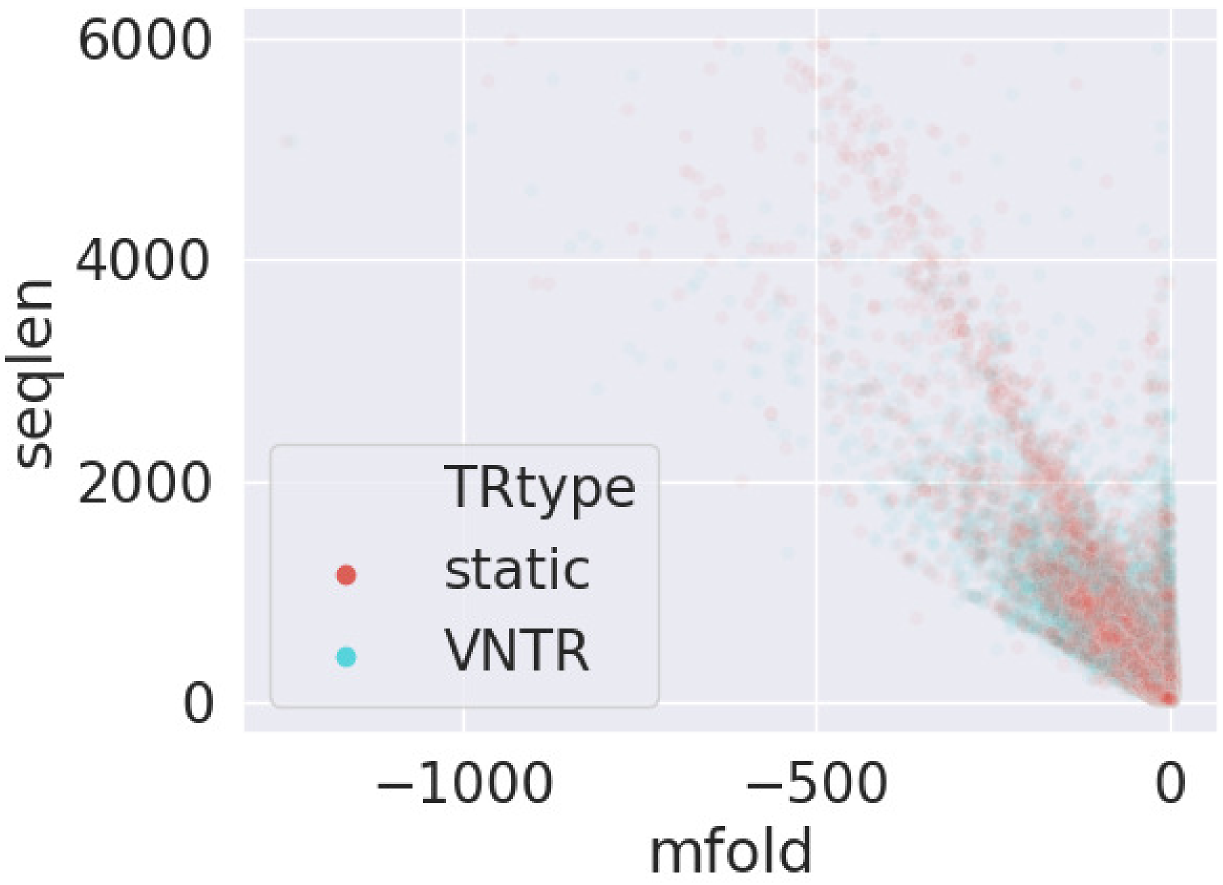


**Fig. S2.** Scatterplot of mfold against sequence length, labeled by longread labeling. Presented ratio reflects the true ratio.

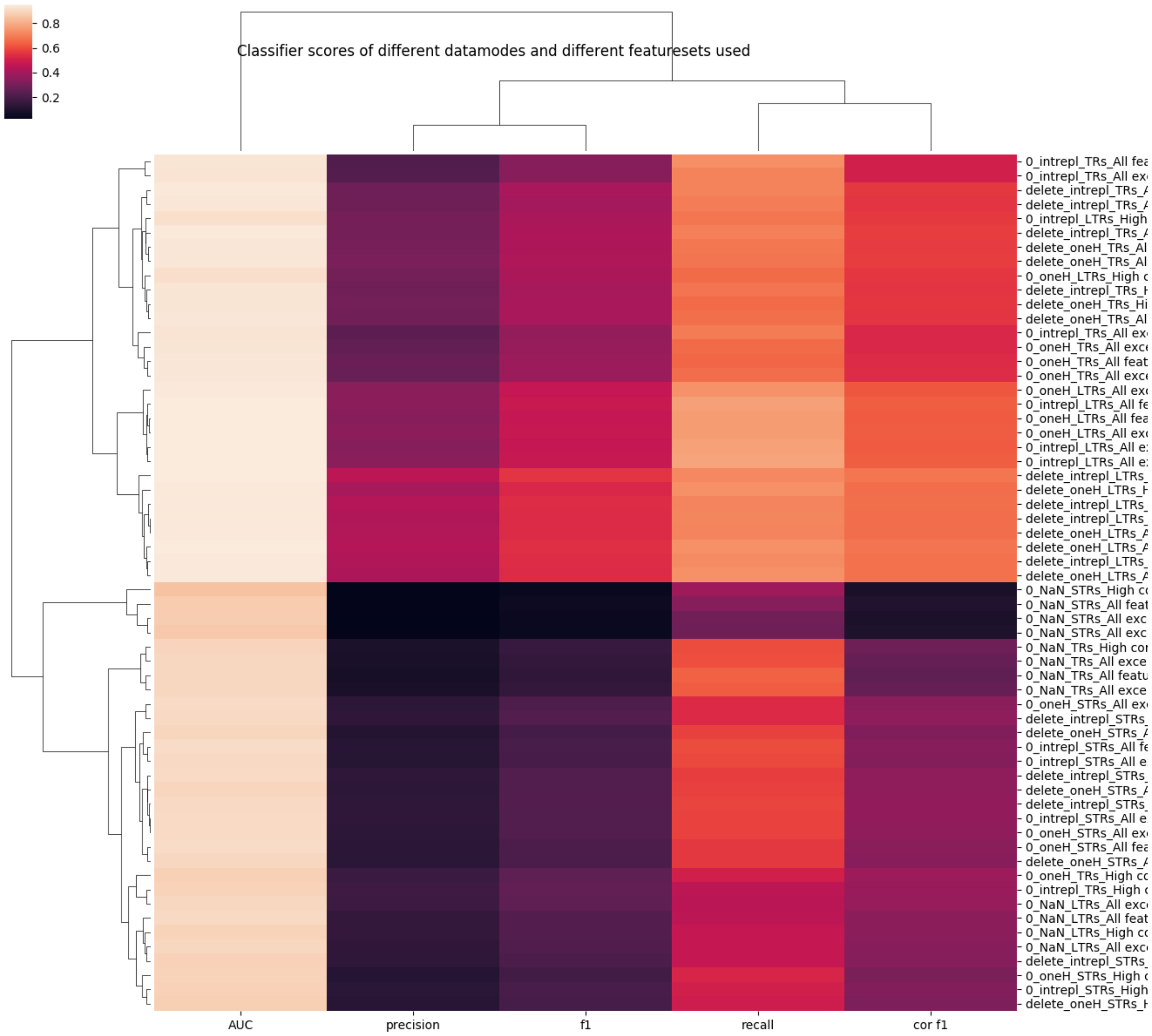


**Fig. S3.** Classifier performance for different featuresets, different handling of categorical data and missing mfold data and different subgroups of TR motiflength. Classifiers are $100$ tree Random Forest Classifiers.


Diederik Cames van Batenburg [1,*], Alexander Gultyaev [1,*] and Marcel Reinders [2]

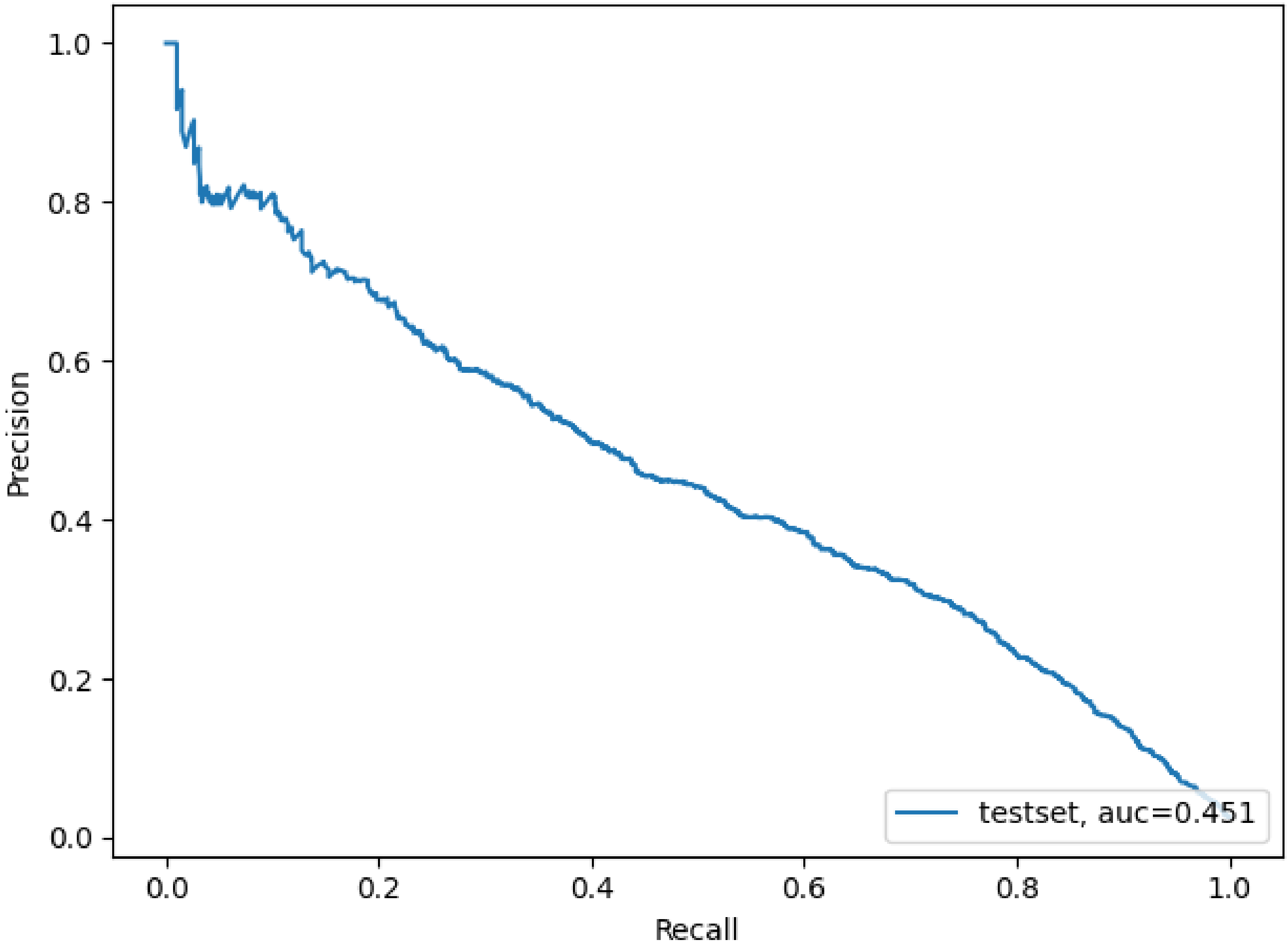


**Fig. S4.** Precision-Recall curve of the final classifier on $10\%$ test data with its area under the curve.

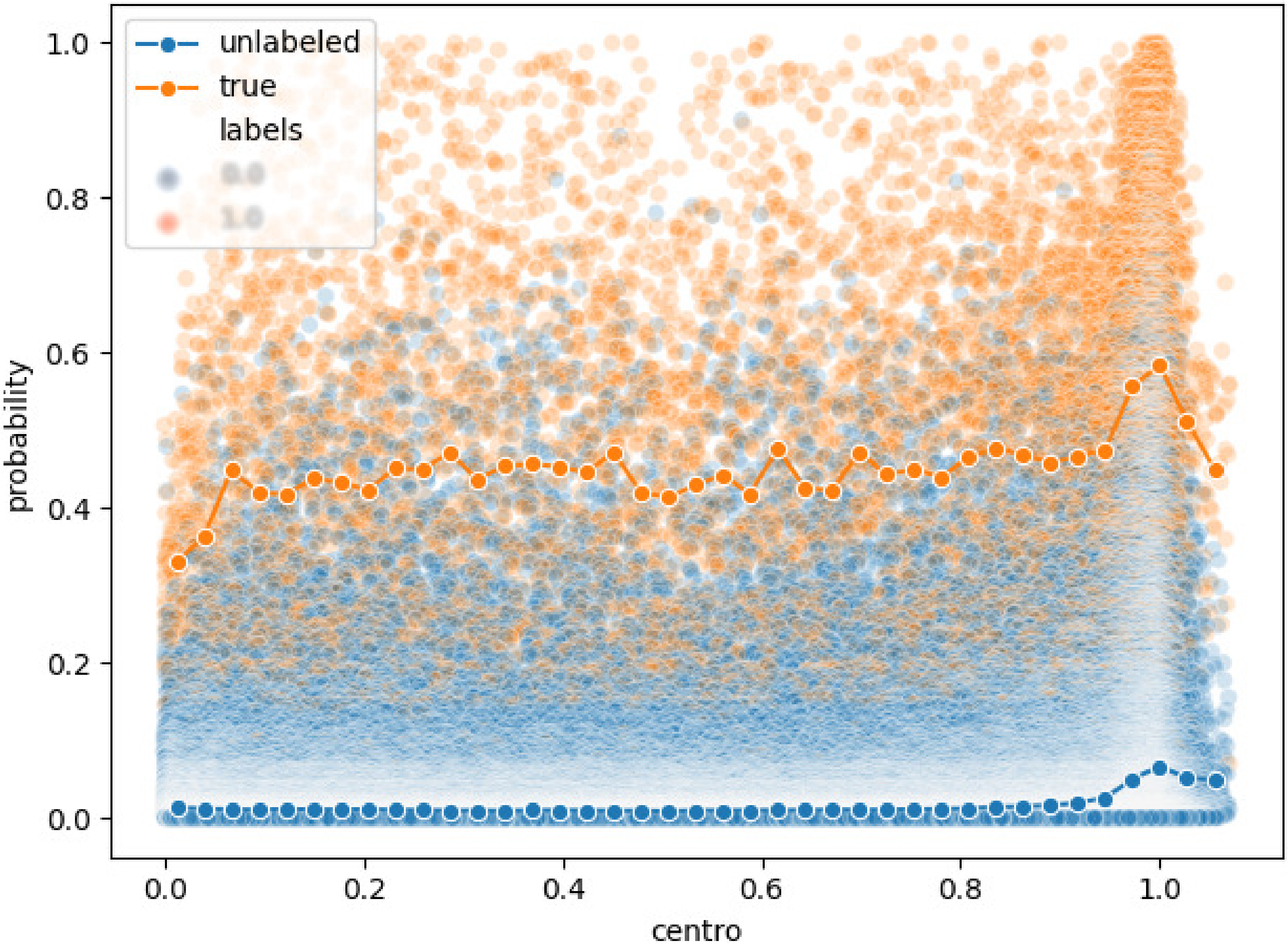


**Fig. S5.** Scatter of classifier predicted VNTR probability over centromeric distance split by long read labeling. Sliding window mean values of classifier predicted probability for both labelings are included.

 Diederik Cames van Batenburg [1,*], Alexander Gultyaev [1,*] and Marcel Reinders [2]

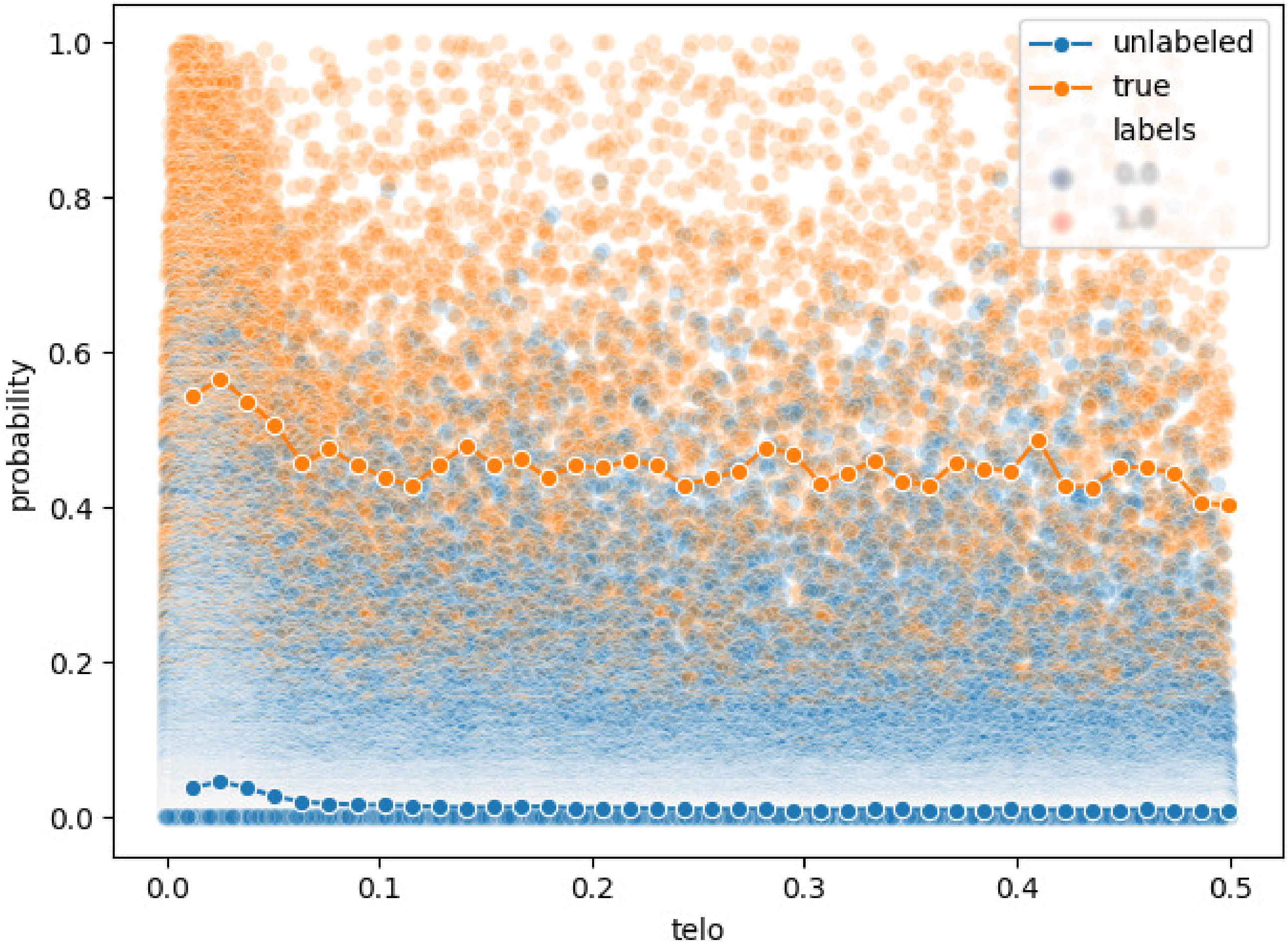


**Fig. S6.** Scatter of classifier predicted VNTR probability over telomeric distance split by long read labeling. Sliding window mean values of classifier predicted probability for both labelings are included.

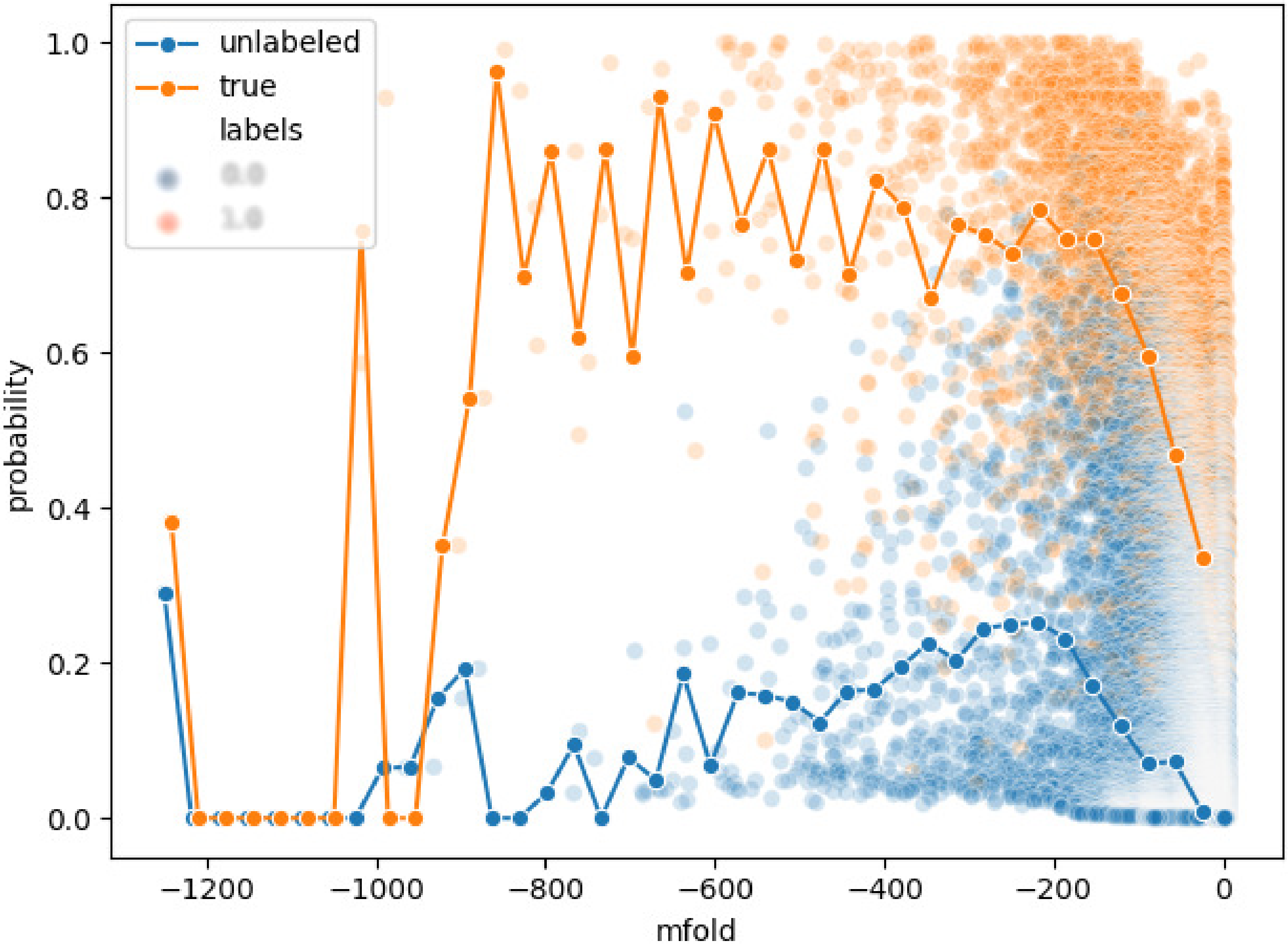


**Fig. S7.** Scatter of classifier predicted VNTR probability over mfold free energy change, split by long read labeling. Sliding window mean values of classifier predicted probability for both labelings are included.

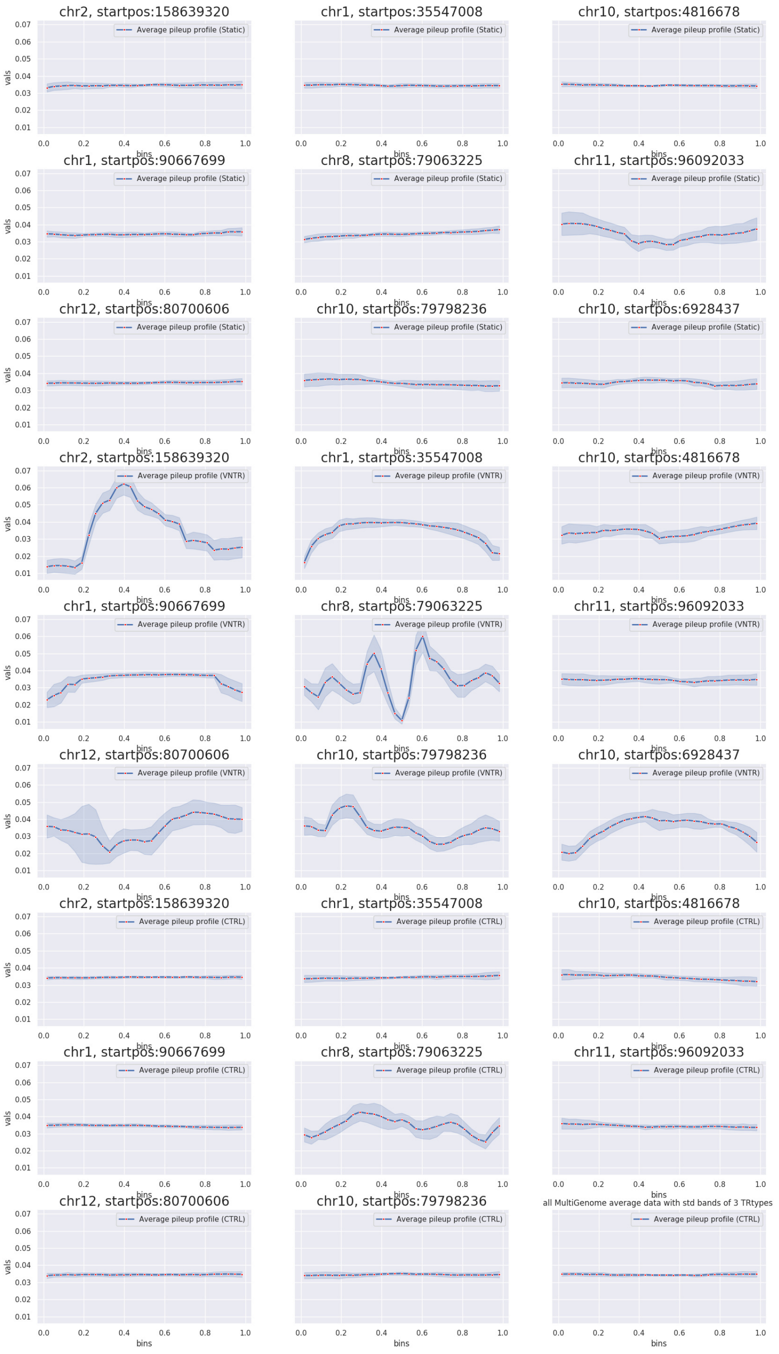


**Fig. S8.** Examples of average pileup for VNTRs, static TRs and control sequences with standard deviation bands. VNTRs show large volatility in pileup and variability along the TR sequence. There is also much variation in patterns between individual VNTRs. TRs are more stable and more similar to control sequences.

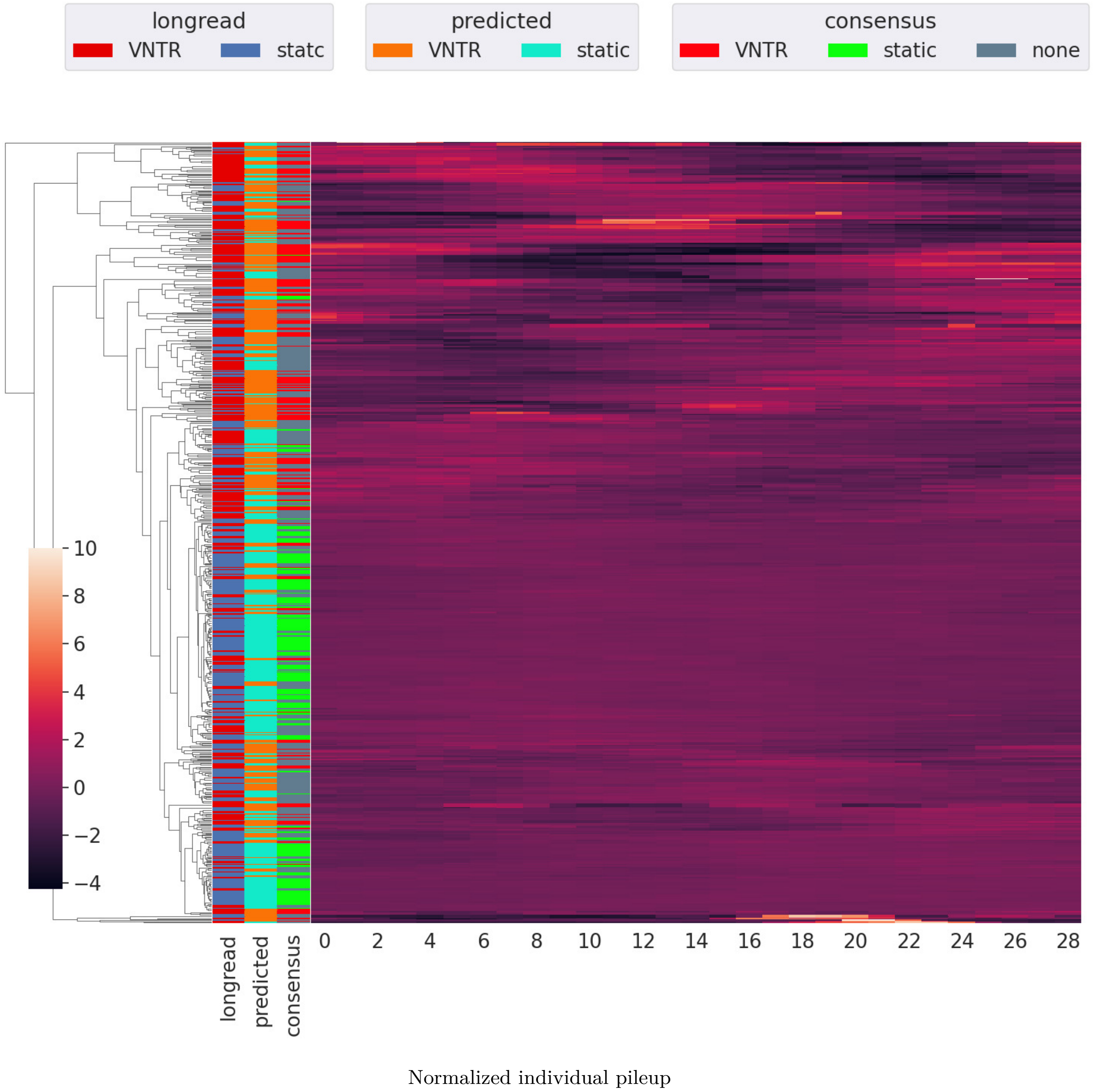


**Fig. S9.** Full linkage hierarchical clustering map of normalized individual pileup profiles from the HG0096 genome, sample of 200 per classifier prediction truthgroup. Normalization is per TR through division by its total pileup.

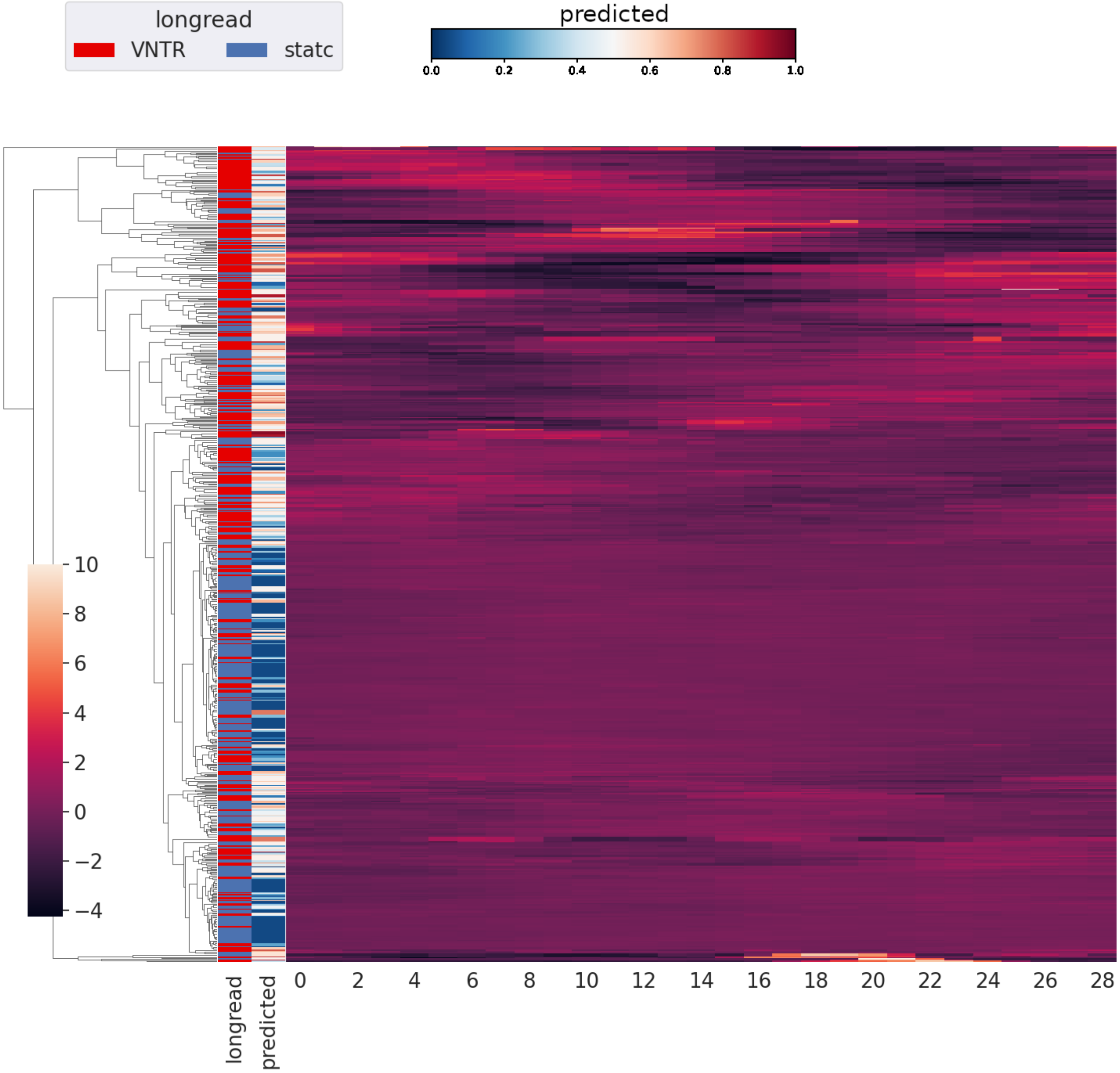


Normalized individual pileup, continuous

**Fig. S10.** Continuous predicted probability labeling of full linkage hierarchical clustering map of normalized individual pileup profiles from the HG0096 genome, sample of 200 per classifier prediction truthgroup. Normalization is done per TR through division by its total pileup.

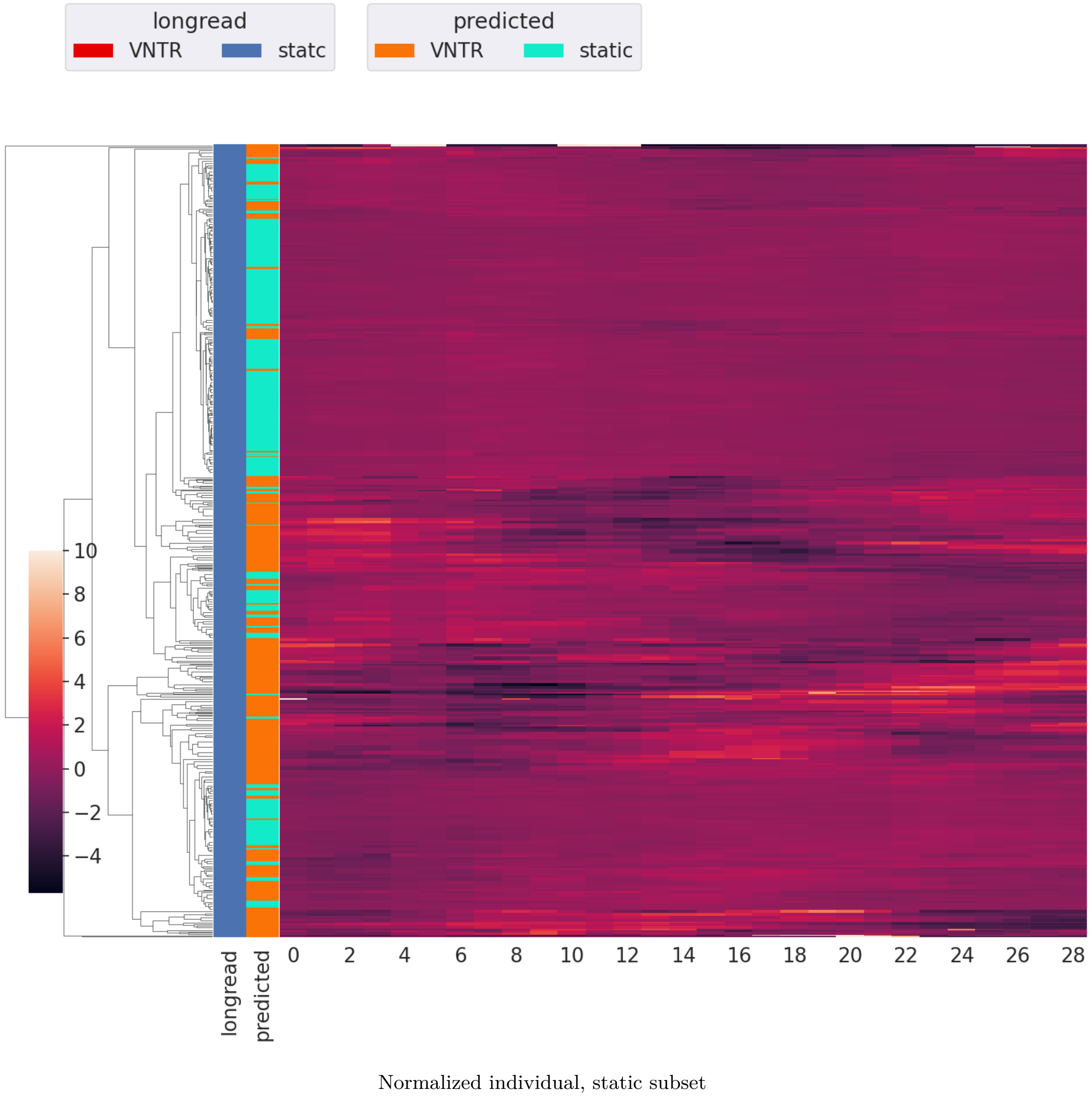


**Fig. S11.** Longread labeled static TRs in full linkage hierarchical clustering map of normalized individual pileup profiles from the HG00096 genome, sample of $400$ per classifier prediction truthgroup. Normalization is per TR through division by its total pileup.

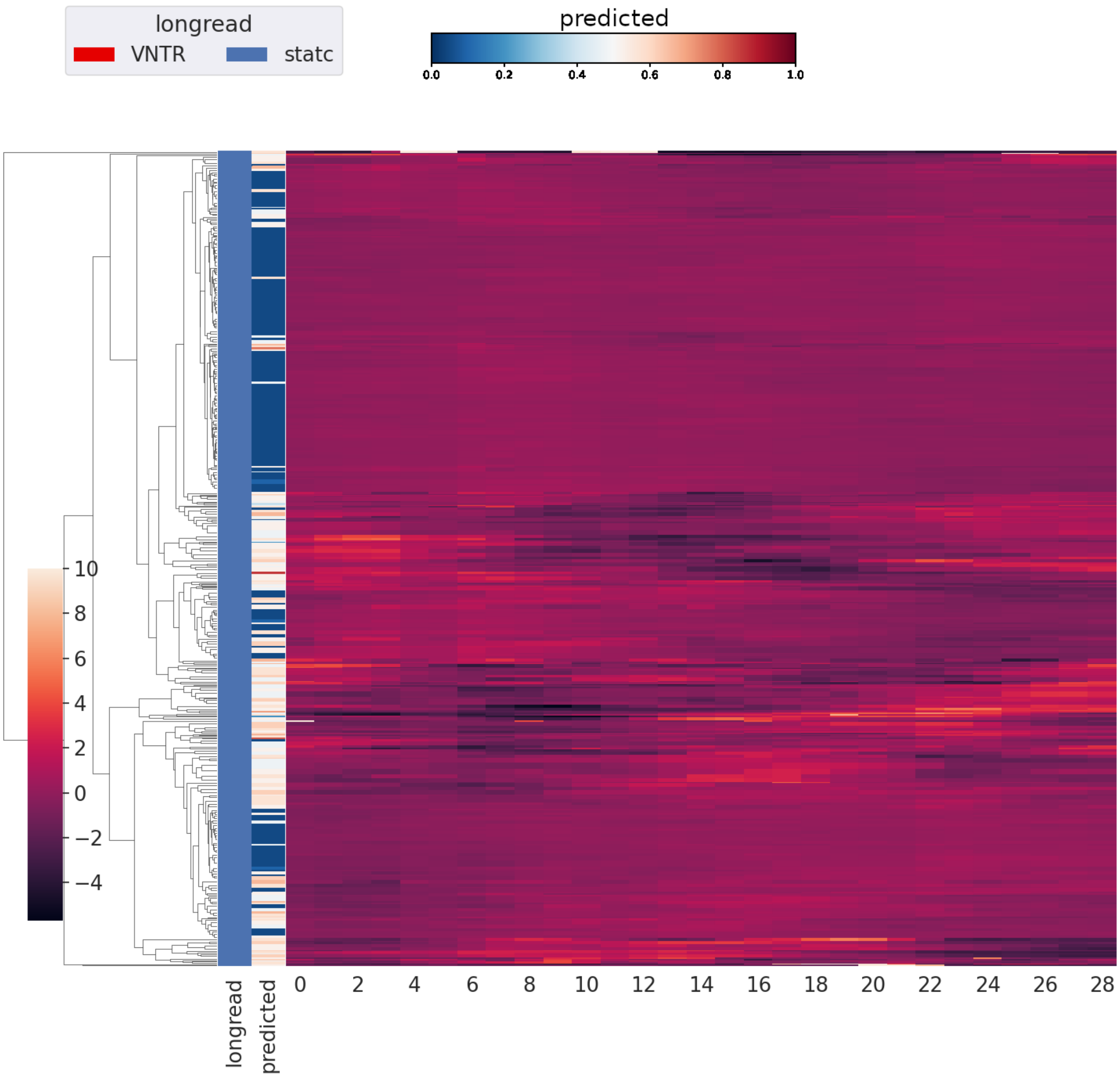


Normalized individual, static subset, continuous

**Fig. S12.** Continuous predicted probability labeling of longread labeled static TRs in full linkage hierarchical clustering map of normalized individual pileup profiles from the HG00096 genome, sample of $400$ per classifier prediction truthgroup. Normalization is done per TR through division by its total pileup.

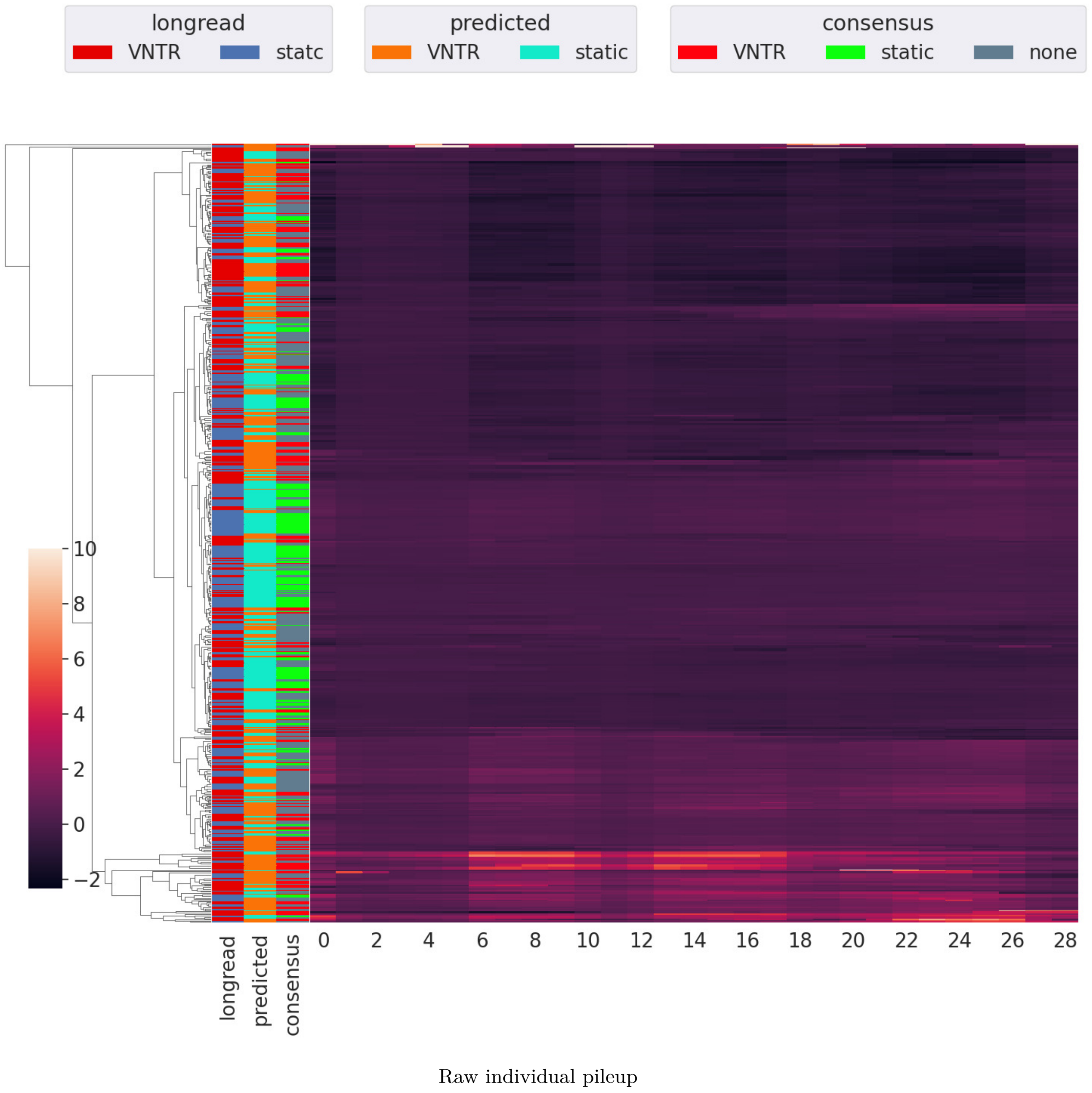


**Fig. S13.** Full linkage hierarchical clustering map of raw individual pileup profiles from the HG0096 genome, sample of 200 per classifier prediction truthgroup.

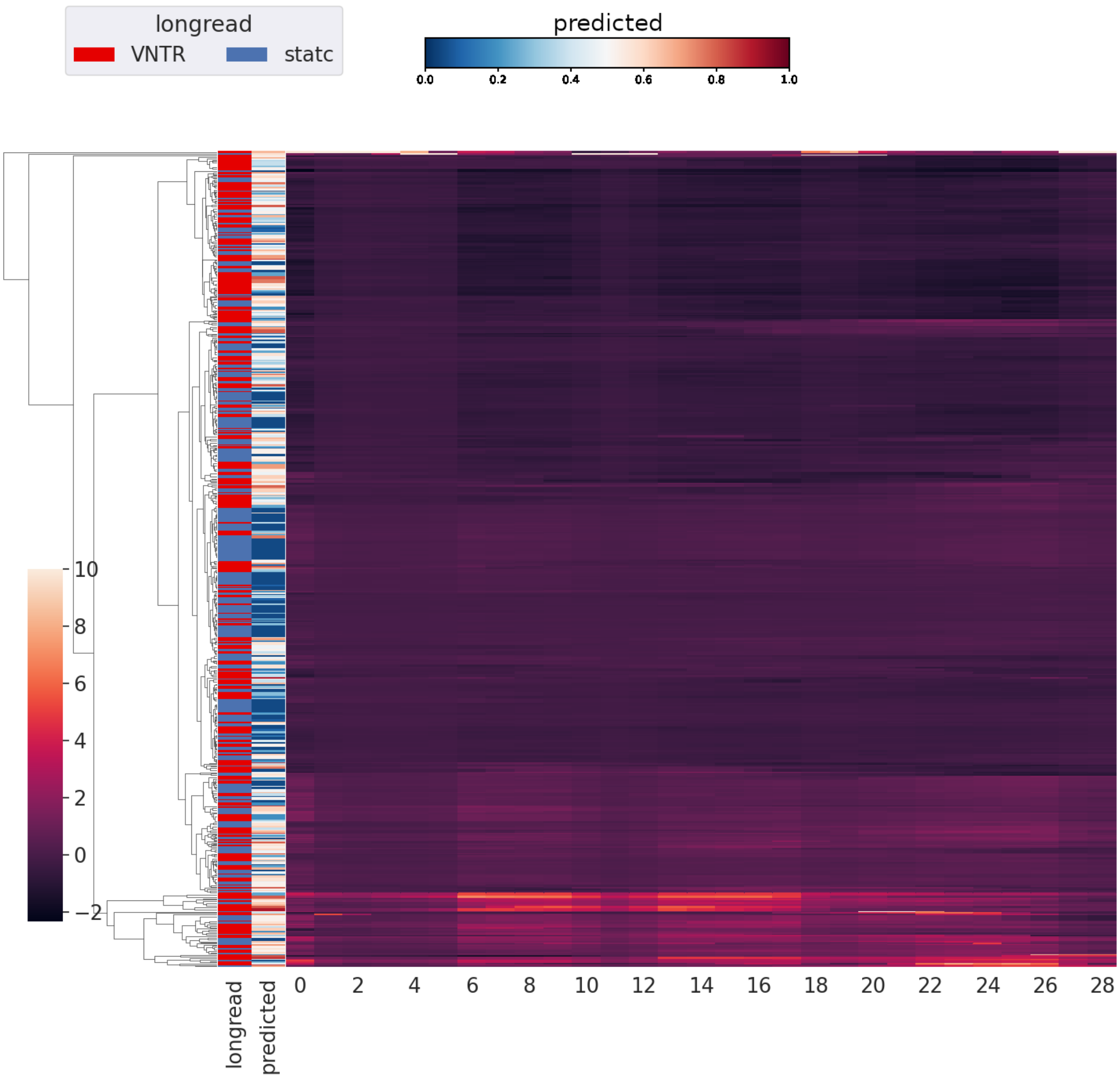


Raw individual pileup, continuous

**Fig. S14.** Continuous predicted probability labeling of full linkage hierarchical clustering map of raw individual pileup profiles from the HG0096 genome, sample of $200$ per classifier prediction truthgroup.

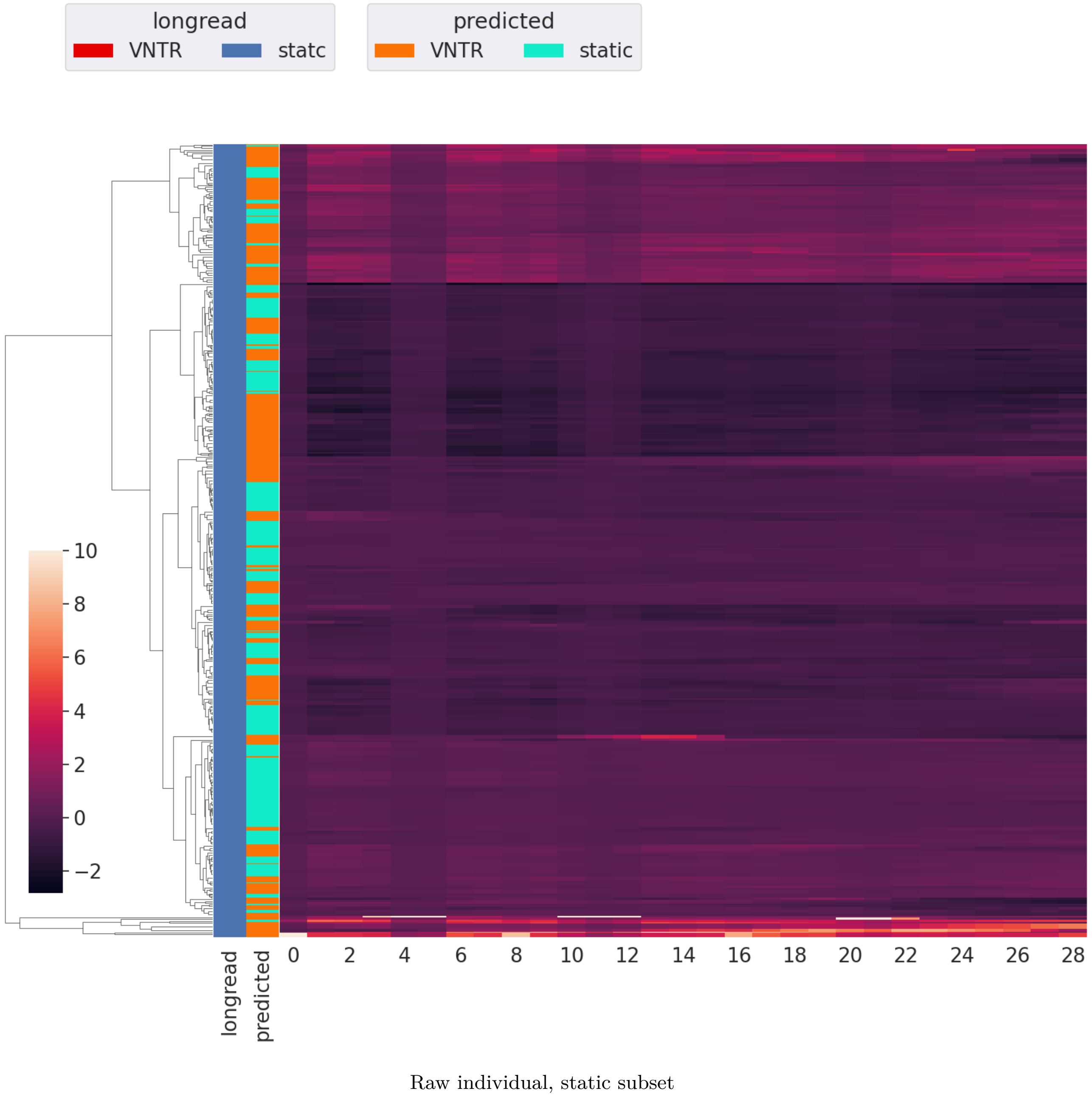


**Fig. S15.** Longread labeled static TRs in full linkage hierarchical clustering map of raw individual pileup profiles from the HG00096 genome, sample of $400$ per classifier prediction truthgroup.

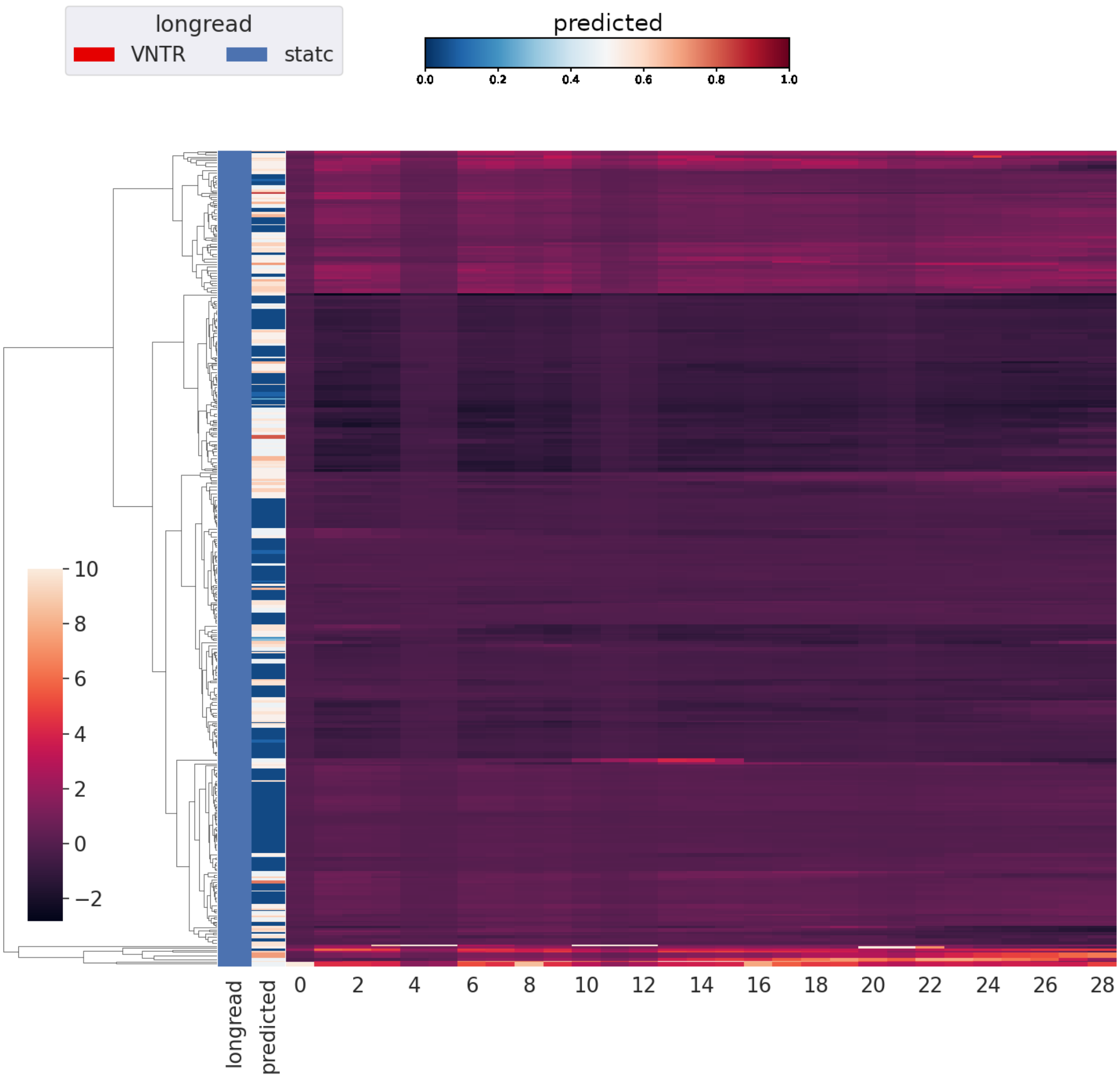


Raw individual, static subset, continuous

**Fig. S16.** Continuous predicted probability labeling of longread labeled static TRs in full linkage hierarchical clustering map of raw individual pileup profiles from the HG00096 genome, sample of $400$ per classifier prediction truthgroup.

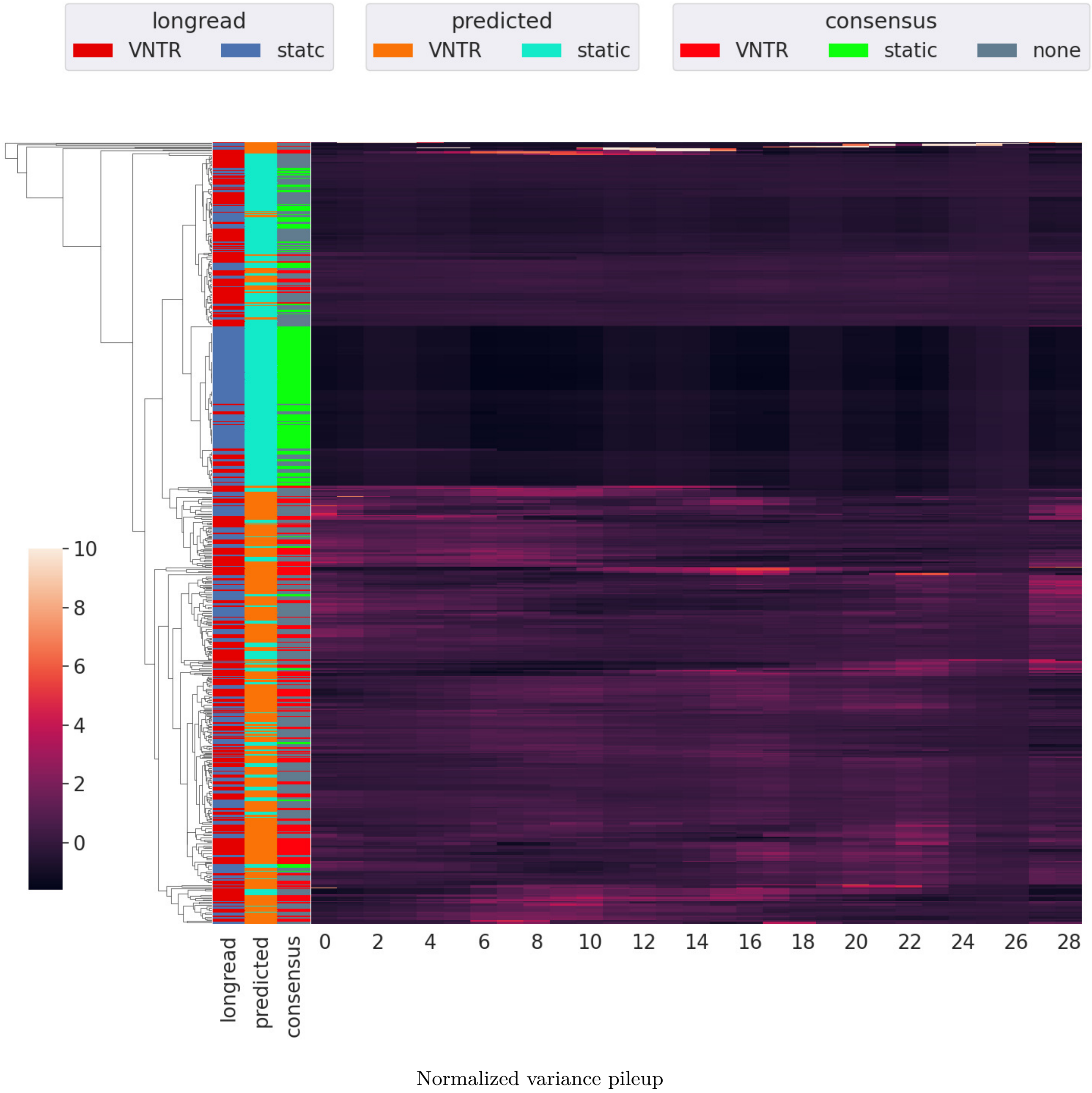


**Fig. S17.** Full linkage hierarchical clustering map of variance profiles over normalized pileup in $17$ genomes, sample of $200$ per classifier prediction truthgroup. Normalization is per TR through division by its total pileup.

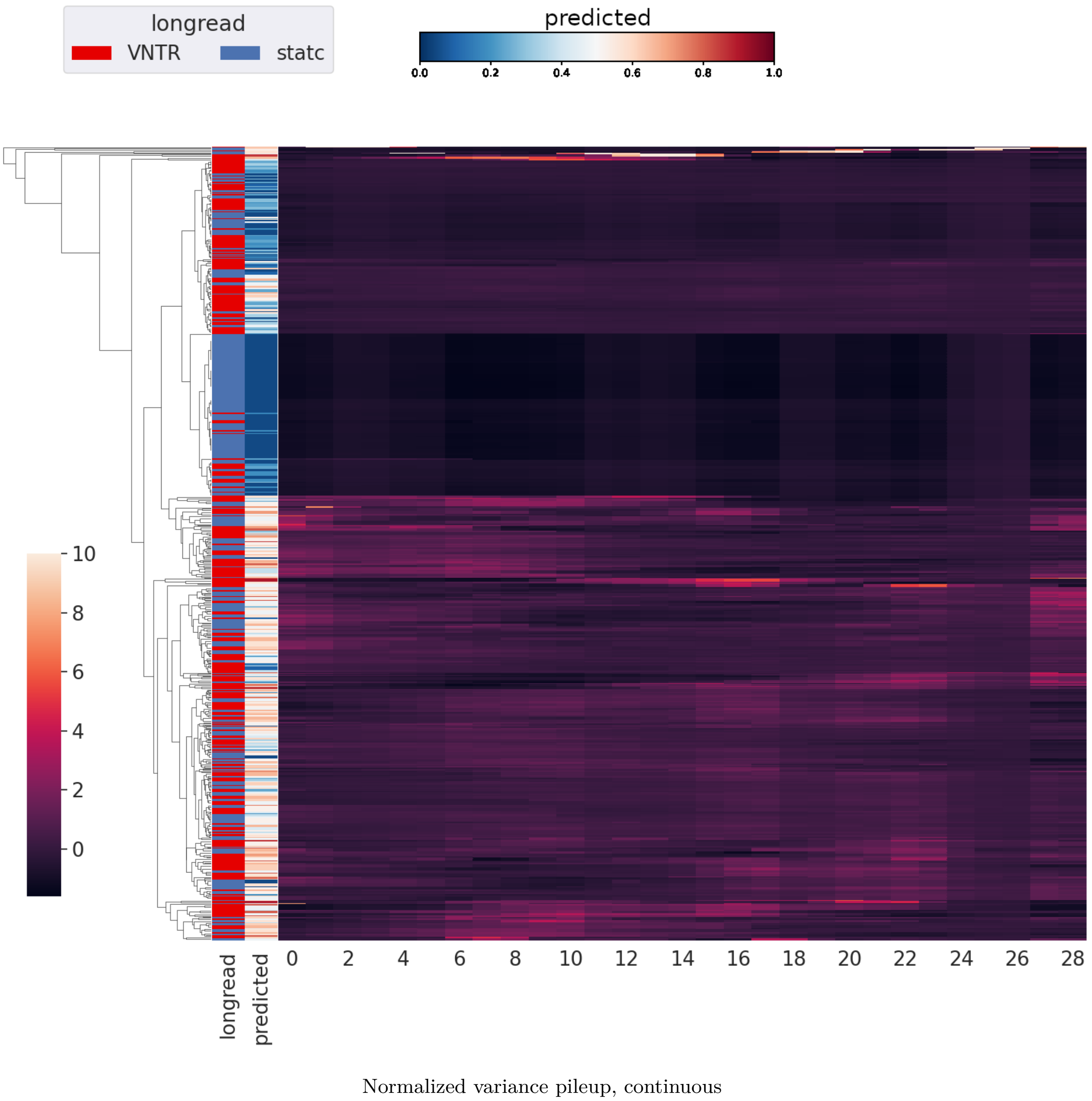


**Fig. S18.** Continuous predicted probability labeling of gull linkage hierarchical clustering map of variance profiles over normalized pileup in $17$ genomes, sample of $200$ per classifier prediction truthgroup. Normalization is per TR through division by its total pileup.

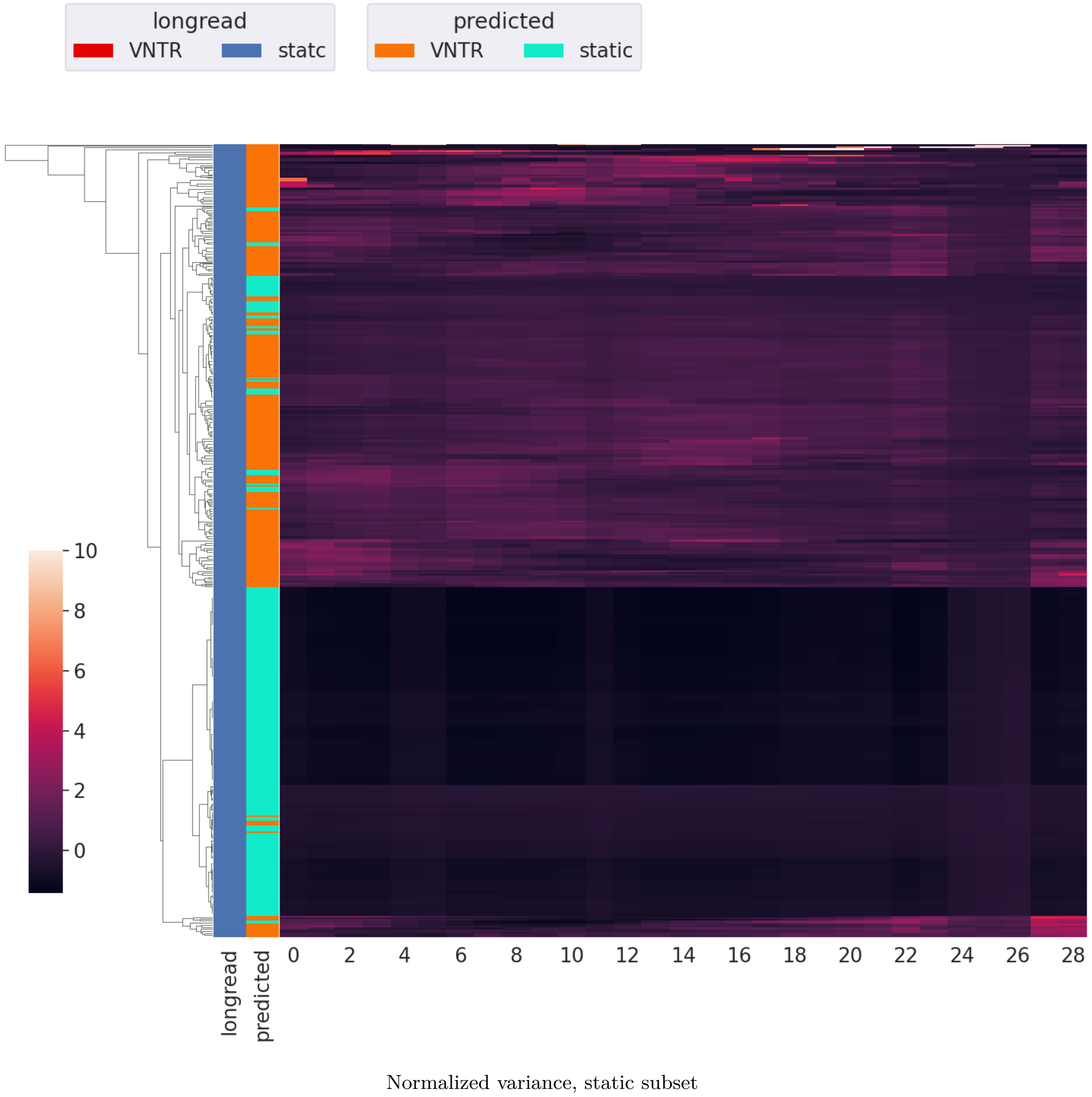


**Fig. S19.** Longread labeled static TRs in full linkage hierarchical clustering map of variance profiles over normalized pileup in $17$ genomes, sample of $200$ per classifier prediction truthgroup. Normalization is per TR through division by its total pileup.

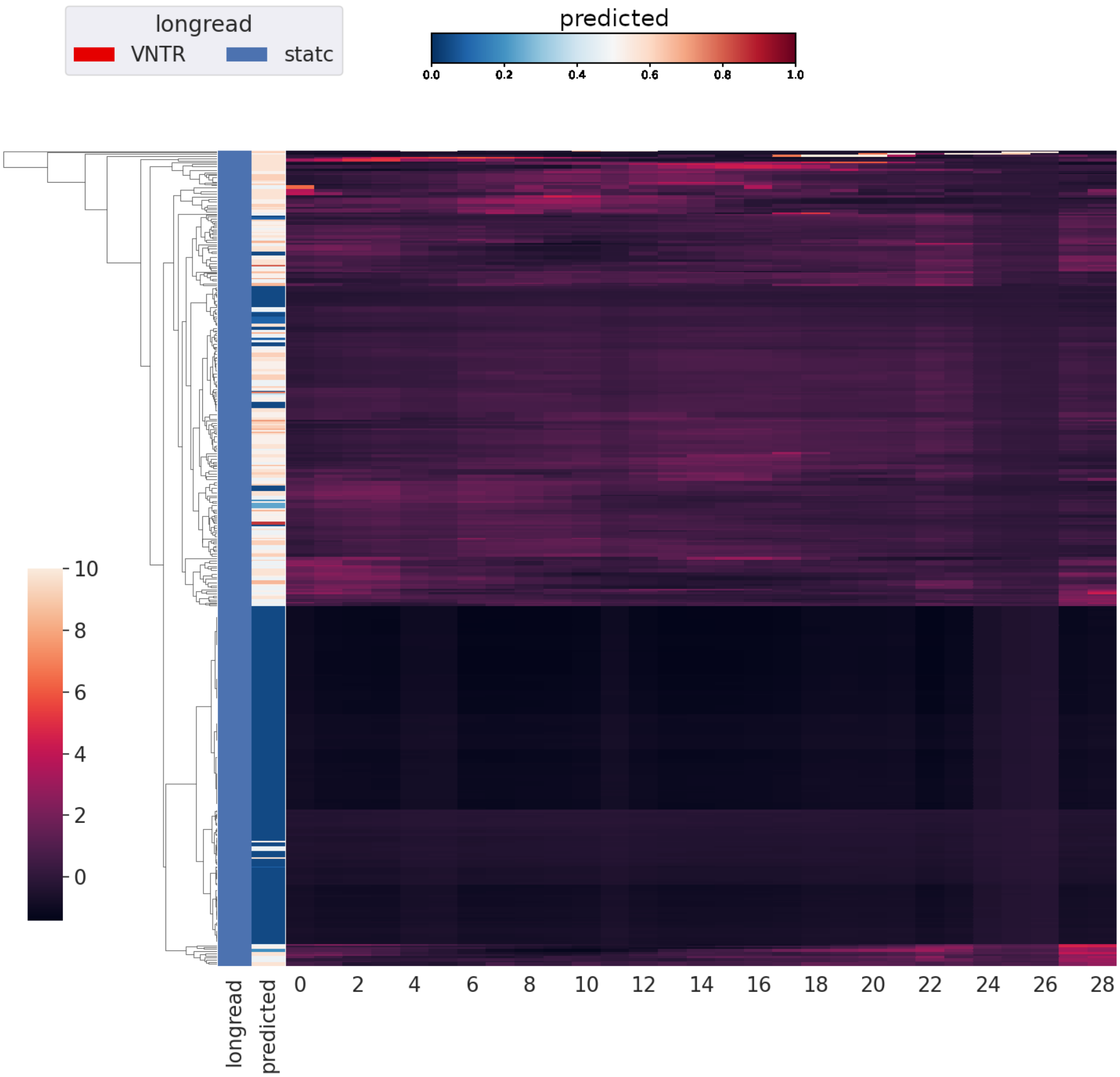


Normalized variance, static subset, continuous

**Fig. S20.** Continuous predicted probability labeling of longread labeled static TRs in full linkage hierarchical clustering map of variance profiles over normalized pileup in $17$ genomes, sample of $200$ per classifier prediction truthgroup. Normalization is per TR through division by its total pileup.

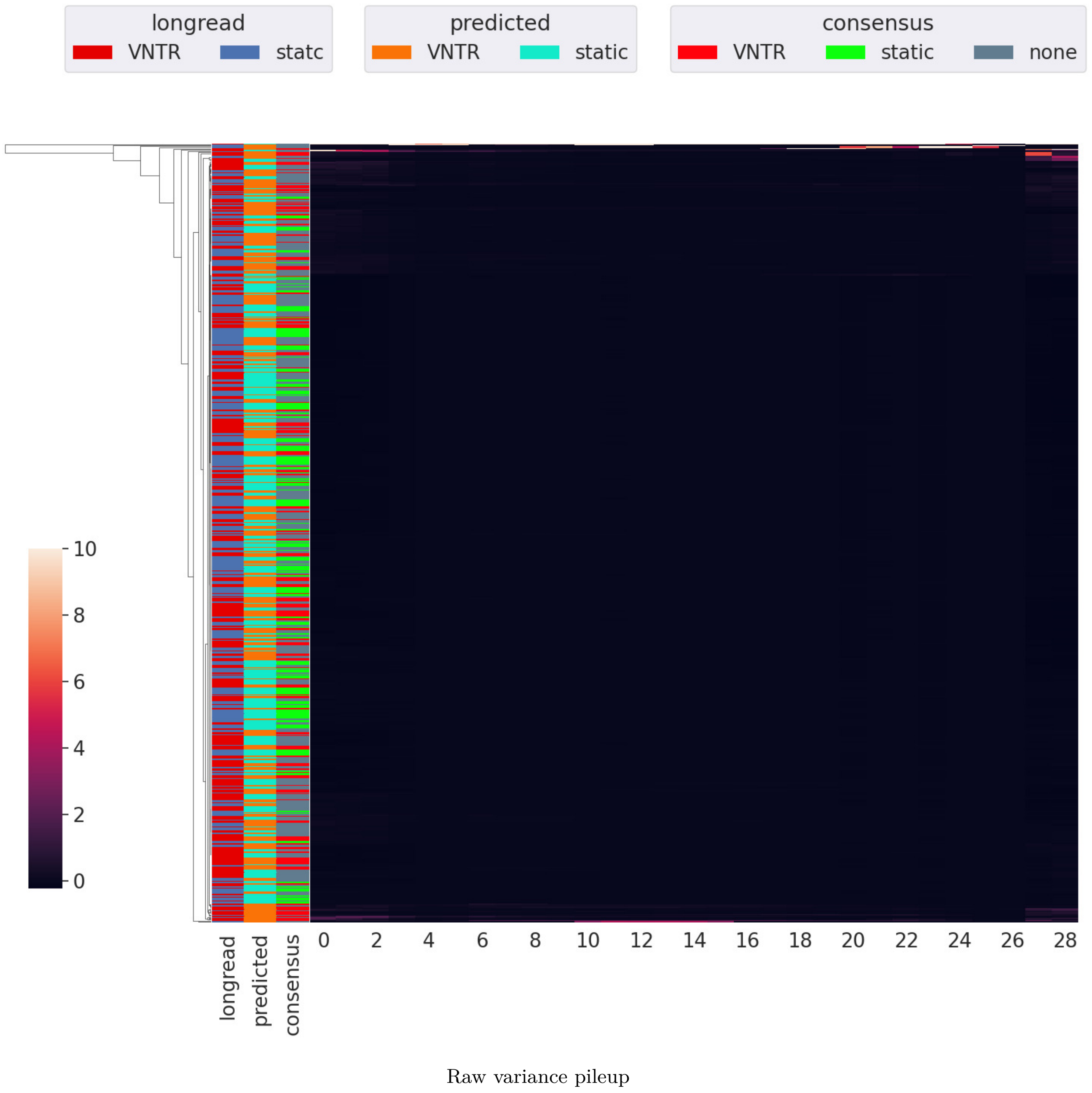


**Fig. S21.** Full linkage hierarchical clustering map of variance profiles over raw pileup in $17$ genomes, sample of $200$ per classifier prediction truthgroup.

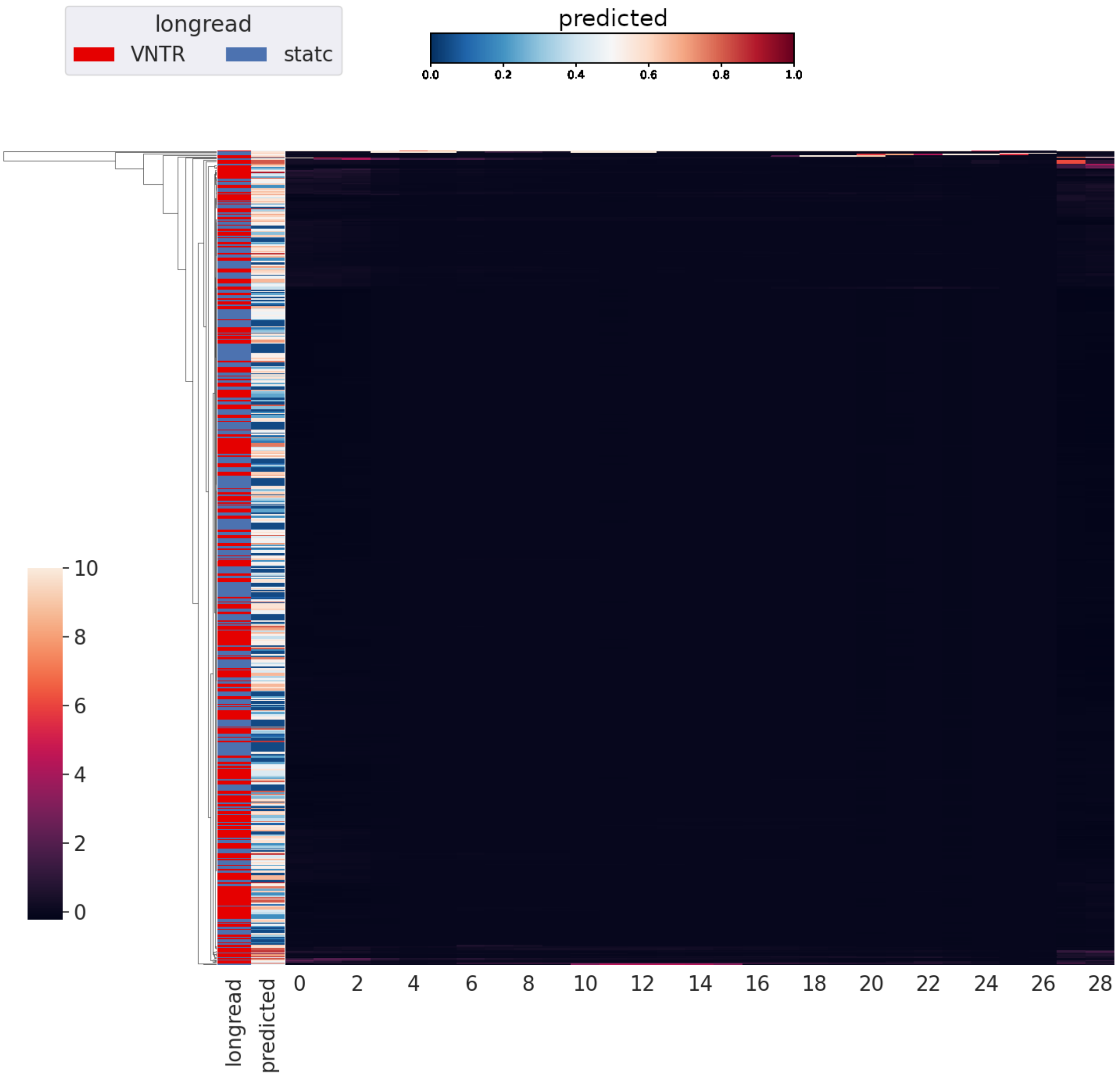


Raw variance pileup, continuous

**Fig. S22.** Continuous predicted probability labeling of gull linkage hierarchical clustering map of variance profiles over raw pileup in $17$ genomes, sample of $200$ per classifier prediction truthgroup.

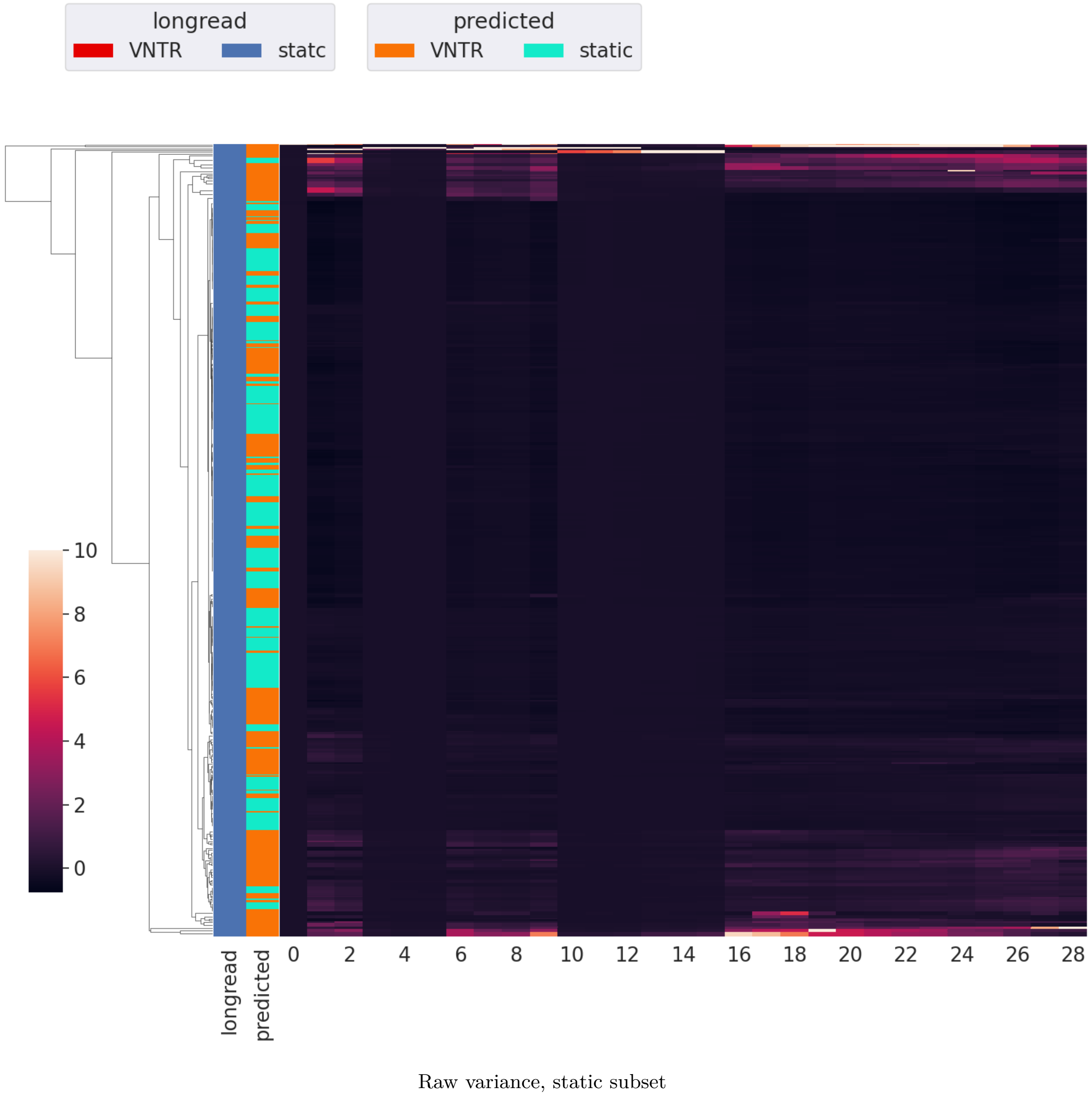


**Fig. S23.** Longread labeled static TRs in full linkage hierarchical clustering map of variance profiles over raw pileup in $17$ genomes, sample of $200$ per classifier prediction truthgroup.

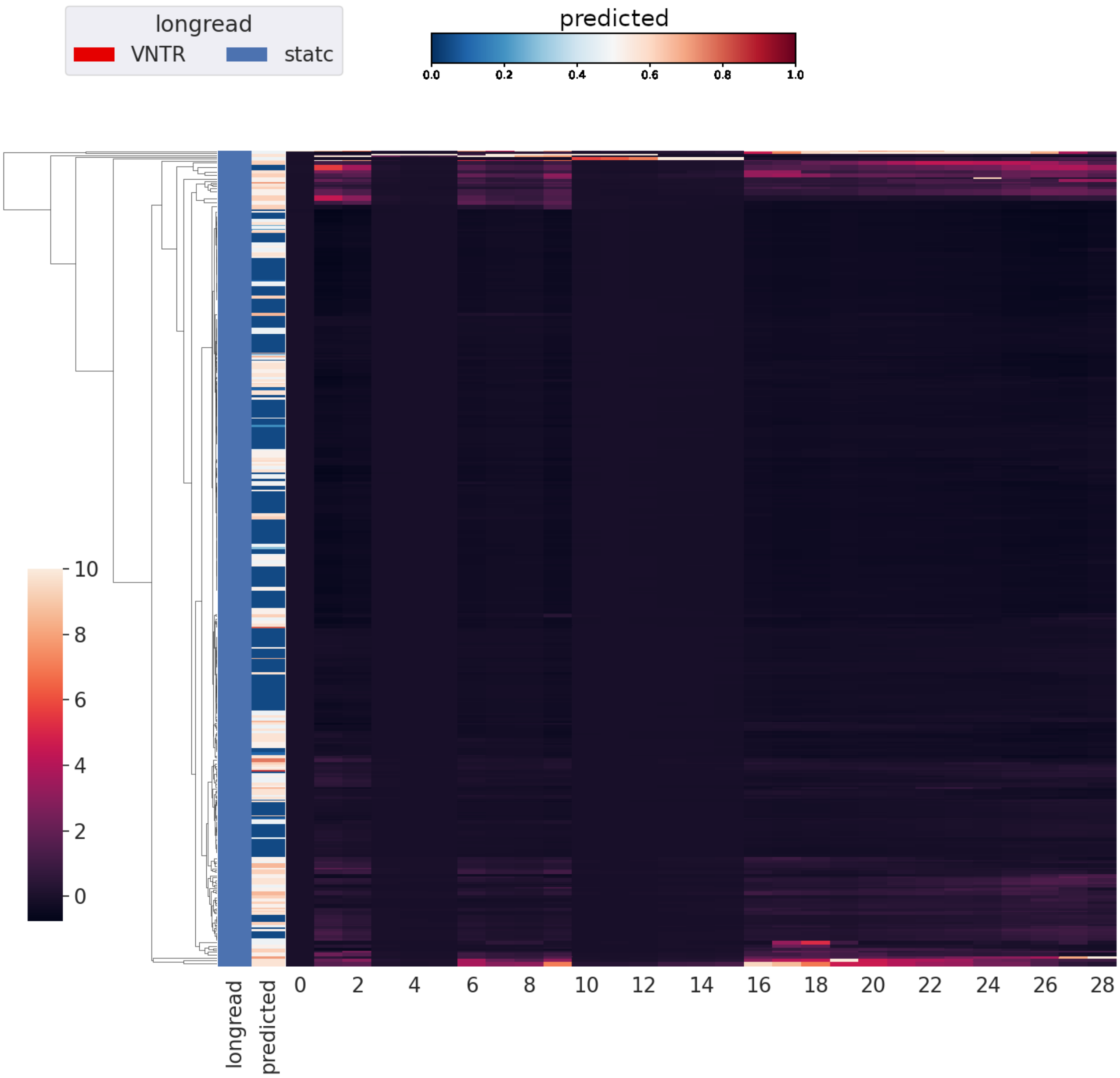


Raw variance, static subset, continuous

**Fig. S24.** Continuous predicted probability labeling of longread labeled static TRs in full linkage hierarchical clustering map of variance profiles over raw pileup in $17$ genomes, sample of $200$ per classifier prediction truthgroup.

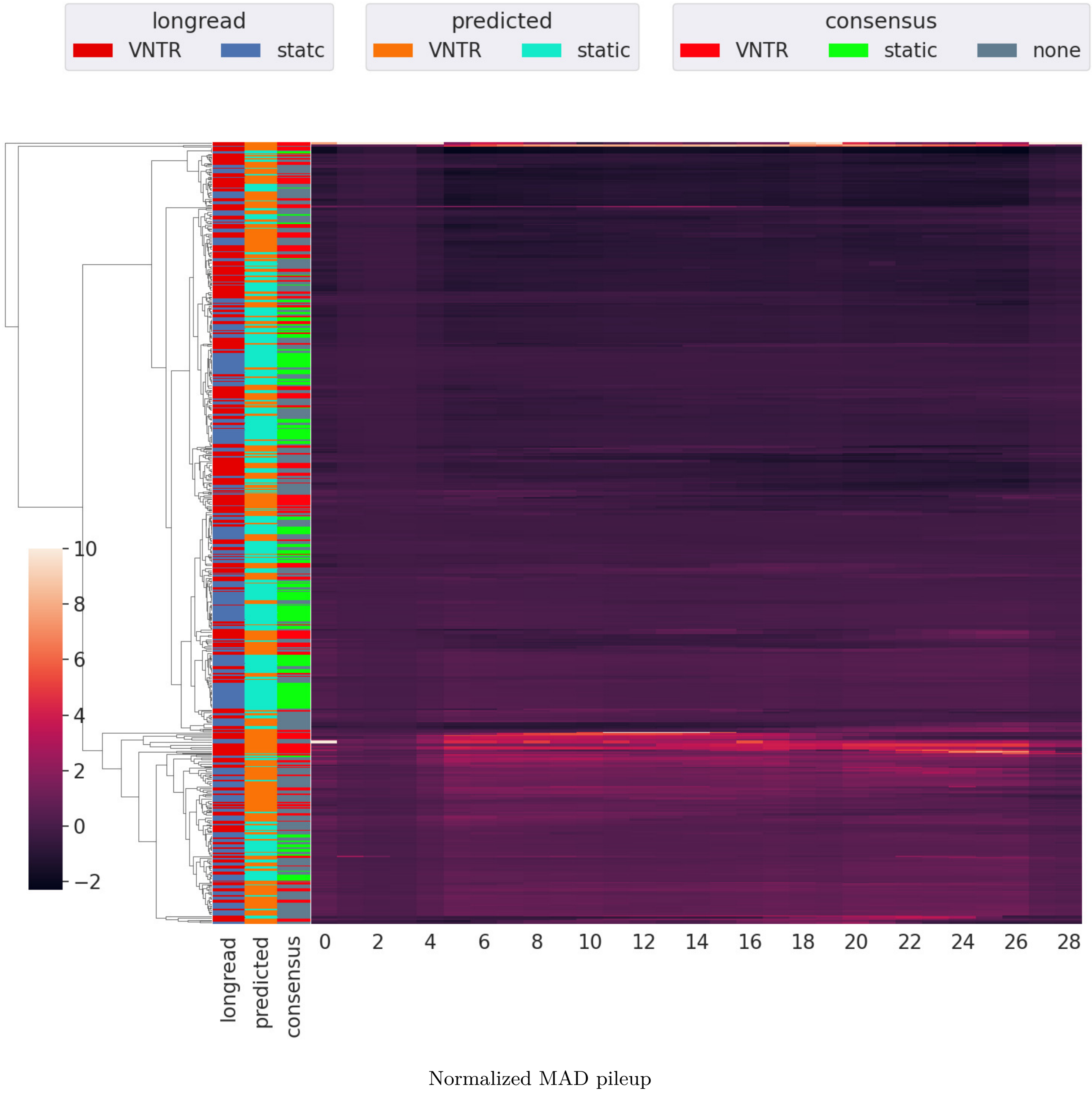


**Fig. S25.** Full linkage hierarchical clustering map of MAD profiles over normalized pileup in $17$ genomes, sample of $200$ per classifier prediction truthgroup. Normalization is per TR through division by its total pileup.

 

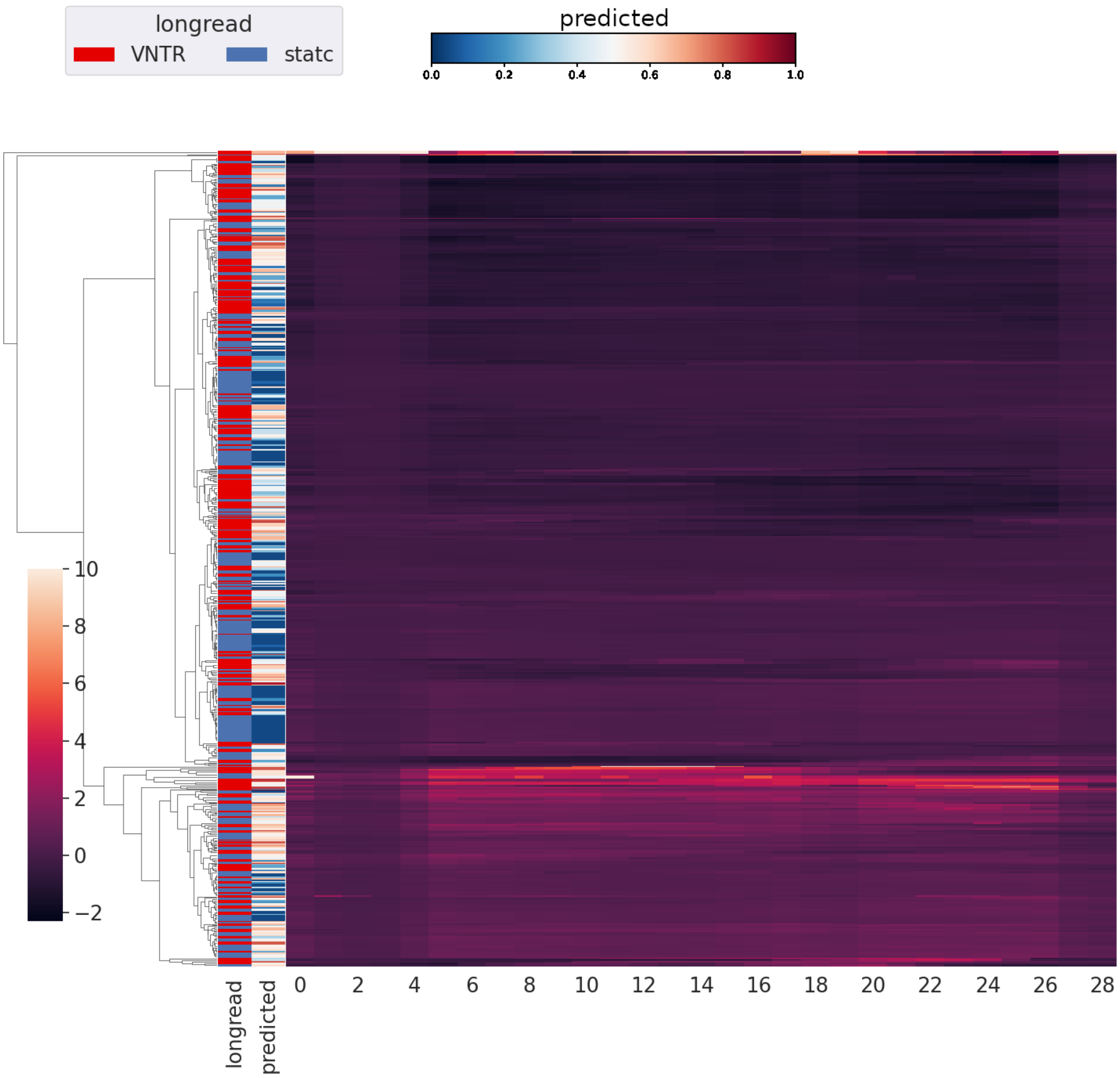


Normalized MAD pileup, continuous

**Fig. S26.** Continuous predicted probability labeling of gull linkage hierarchical clustering map of MAD profiles over normalized pileup in $17$ genomes, sample of $200$ per classifier prediction truthgroup. Normalization is per TR through division by its total pileup.

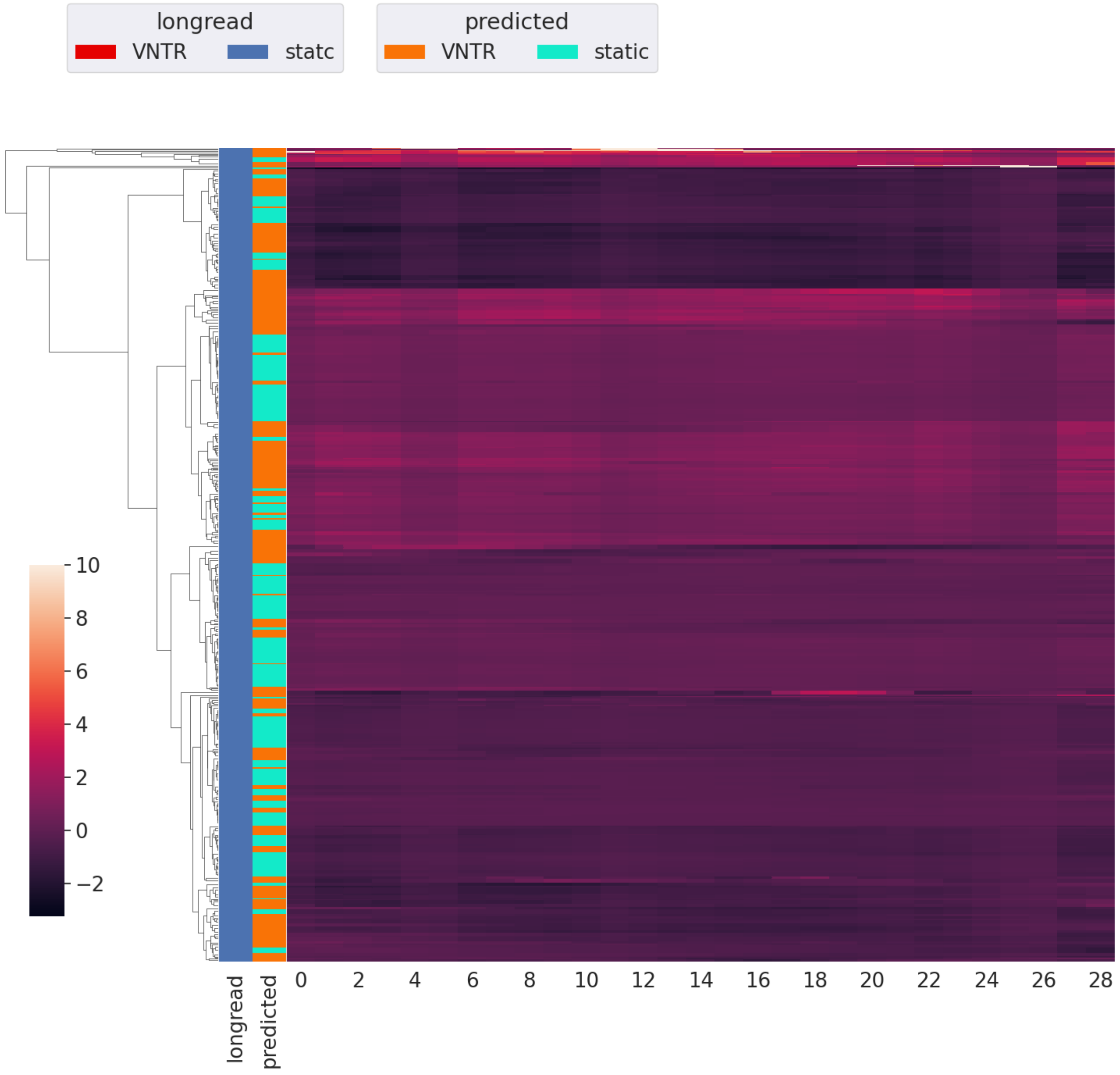


Normalized MAD, static subset

**Fig. S27.** Longread labeled static TRs in full linkage hierarchical clustering map of MAD profiles over normalized pileup in $17$ genomes, sample of $200$ per classifier prediction truthgroup. Normalization is per TR through division by its total pileup.

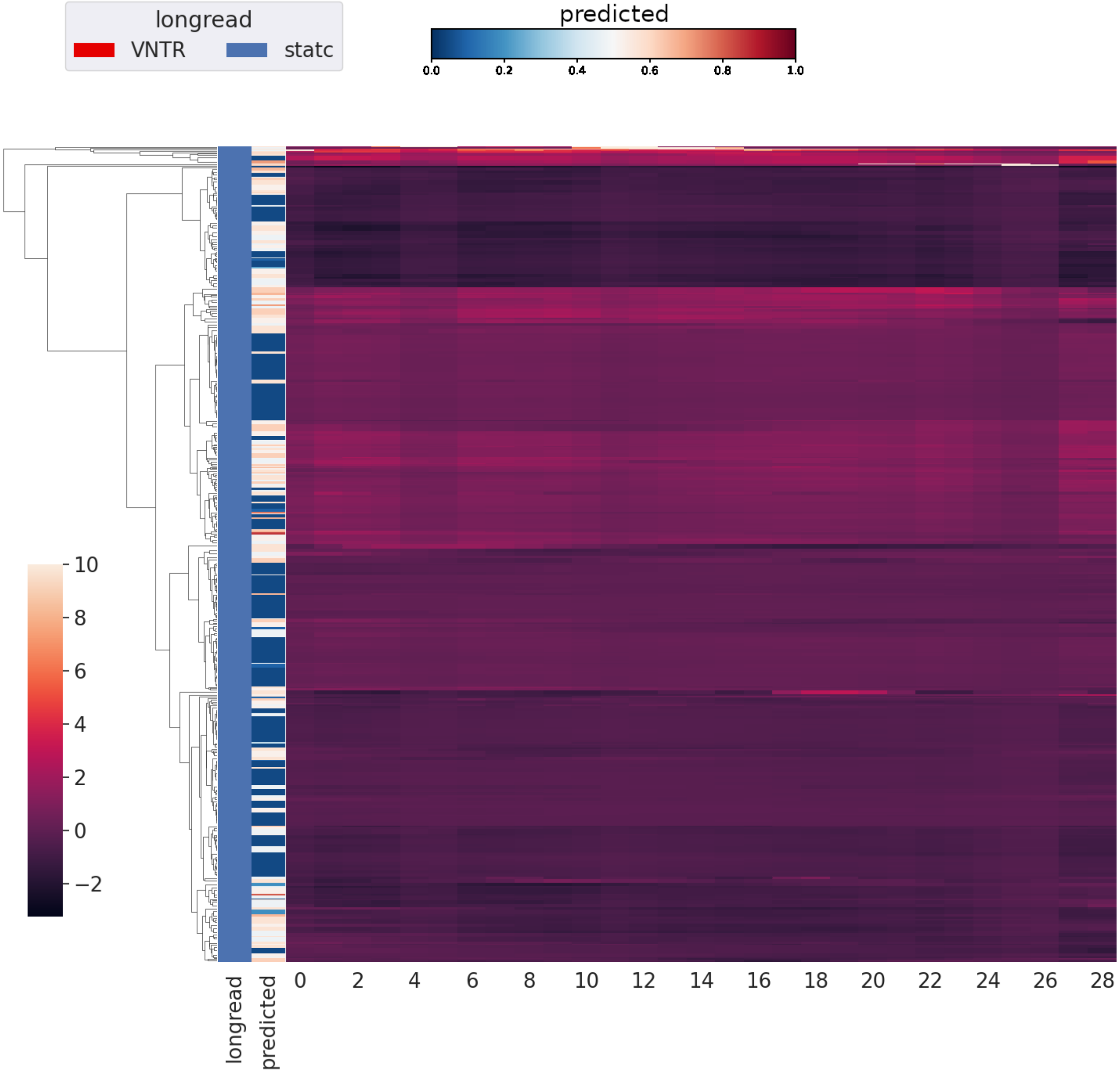


Normalized MAD, static subset, continuous

**Fig. S28.** Continuous predicted probability labeling of longread labeled static TRs in full linkage hierarchical clustering map of MAD profiles over normalized pileup in $17$ genomes, sample of $200$ per classifier prediction truthgroup. Normalization is per TR through division by its total pileup.

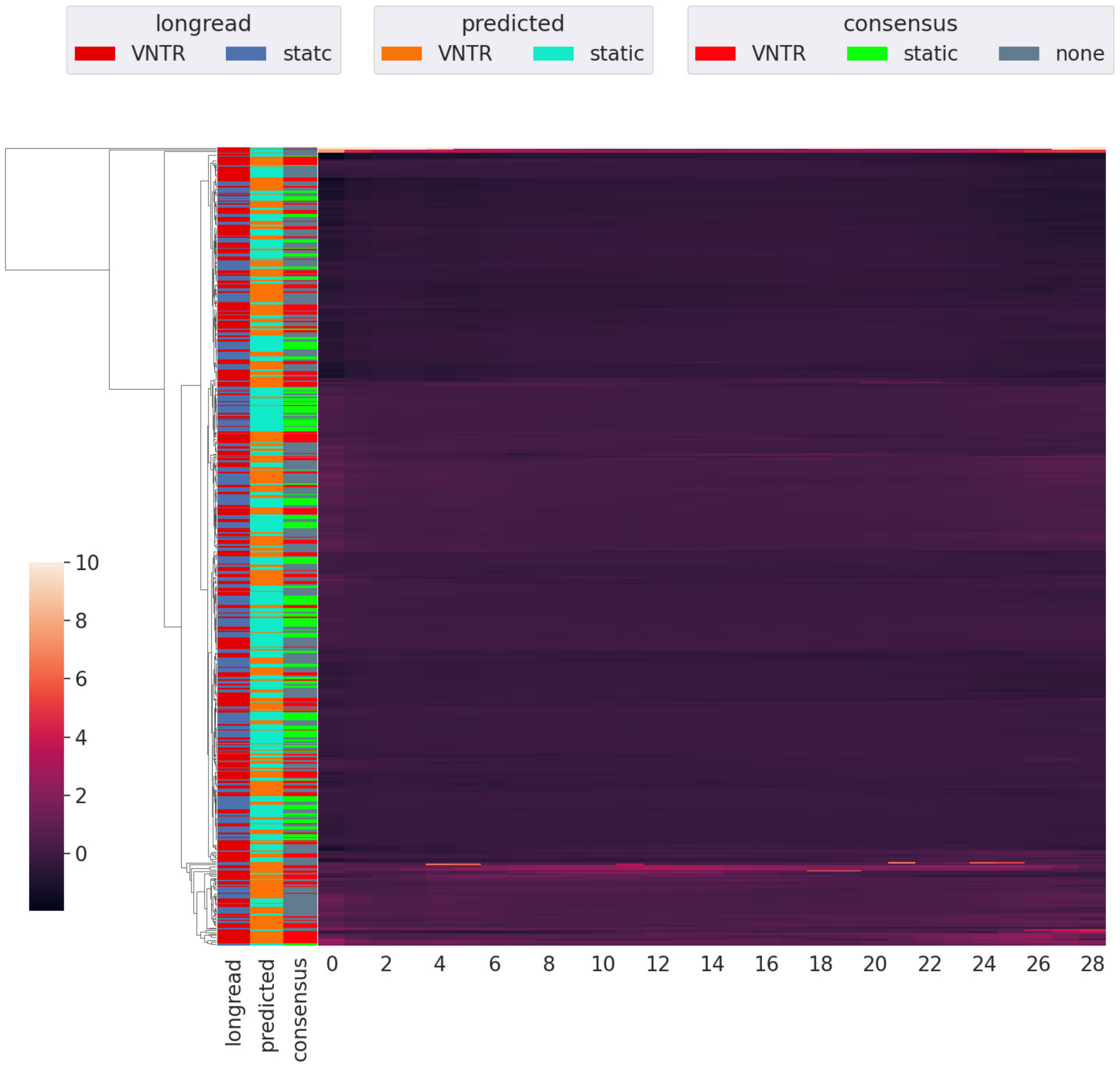


Raw MAD pileup

**Fig. S29.** Full linkage hierarchical clustering map of MAD profiles over raw pileup in $17$ genomes, sample of $200$ per classifier prediction truthgroup.

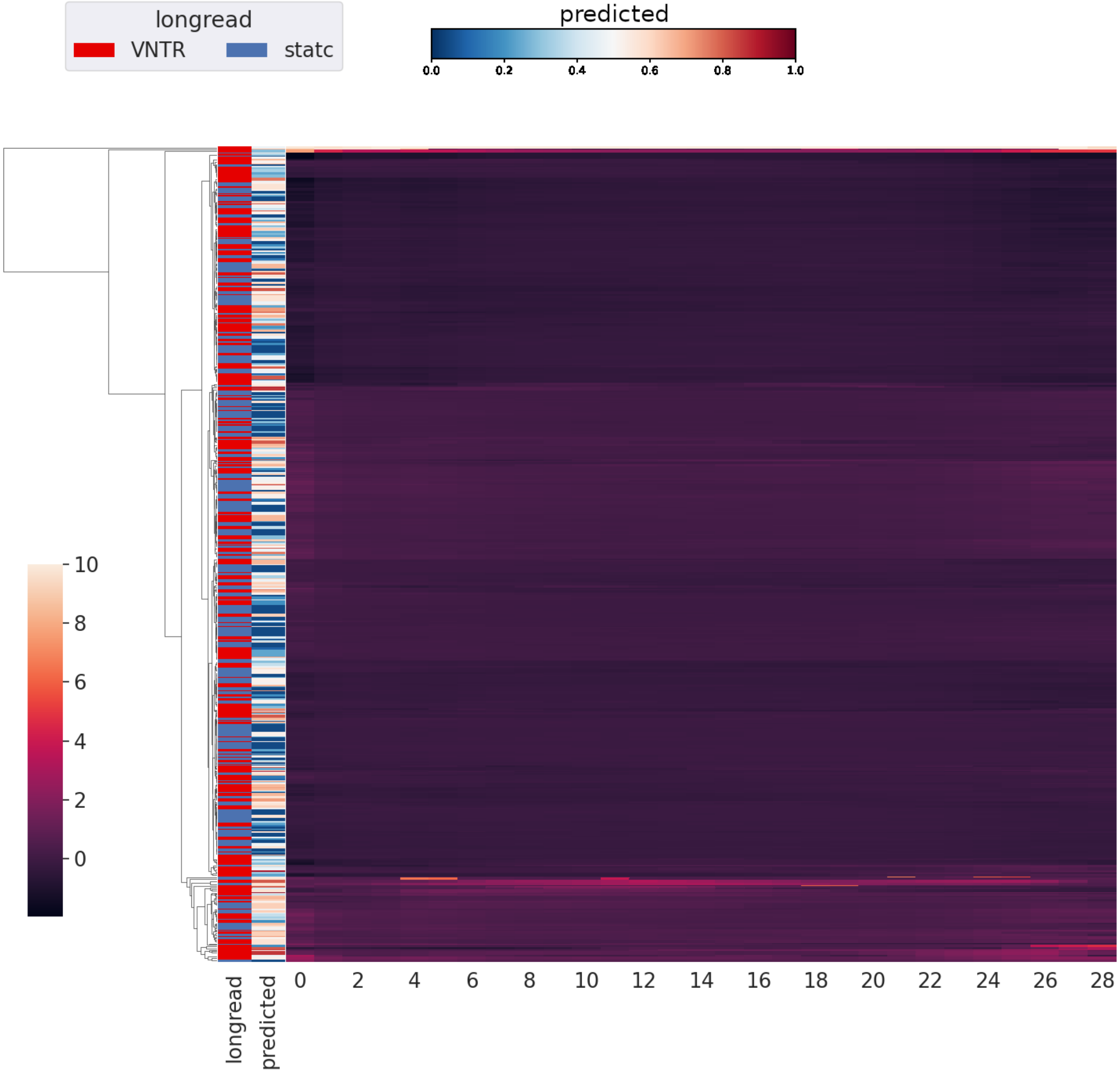


Raw MAD pileup, continuous

**Fig. S30.** Continuous predicted probability labeling of gull linkage hierarchical clustering map of MAD profiles over raw pileup in $17$ genomes, sample of $200$ per classifier prediction truthgroup.

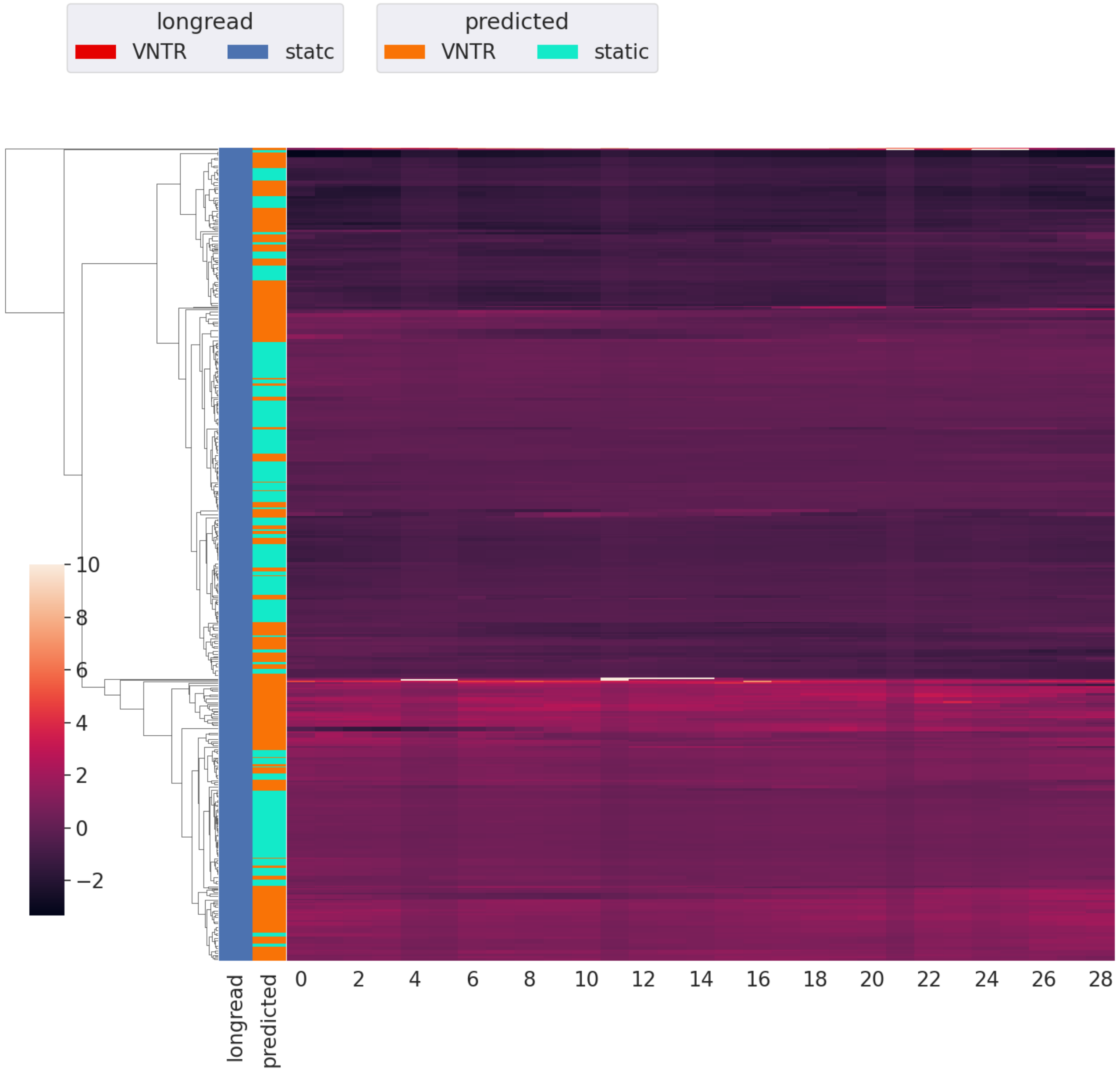


Raw MAD, static subset

**Fig. S31.** Longread labeled static TRs in full linkage hierarchical clustering map of MAD profiles over raw pileup in 17 genomes, sample of 200 per classifier prediction truthgroup.

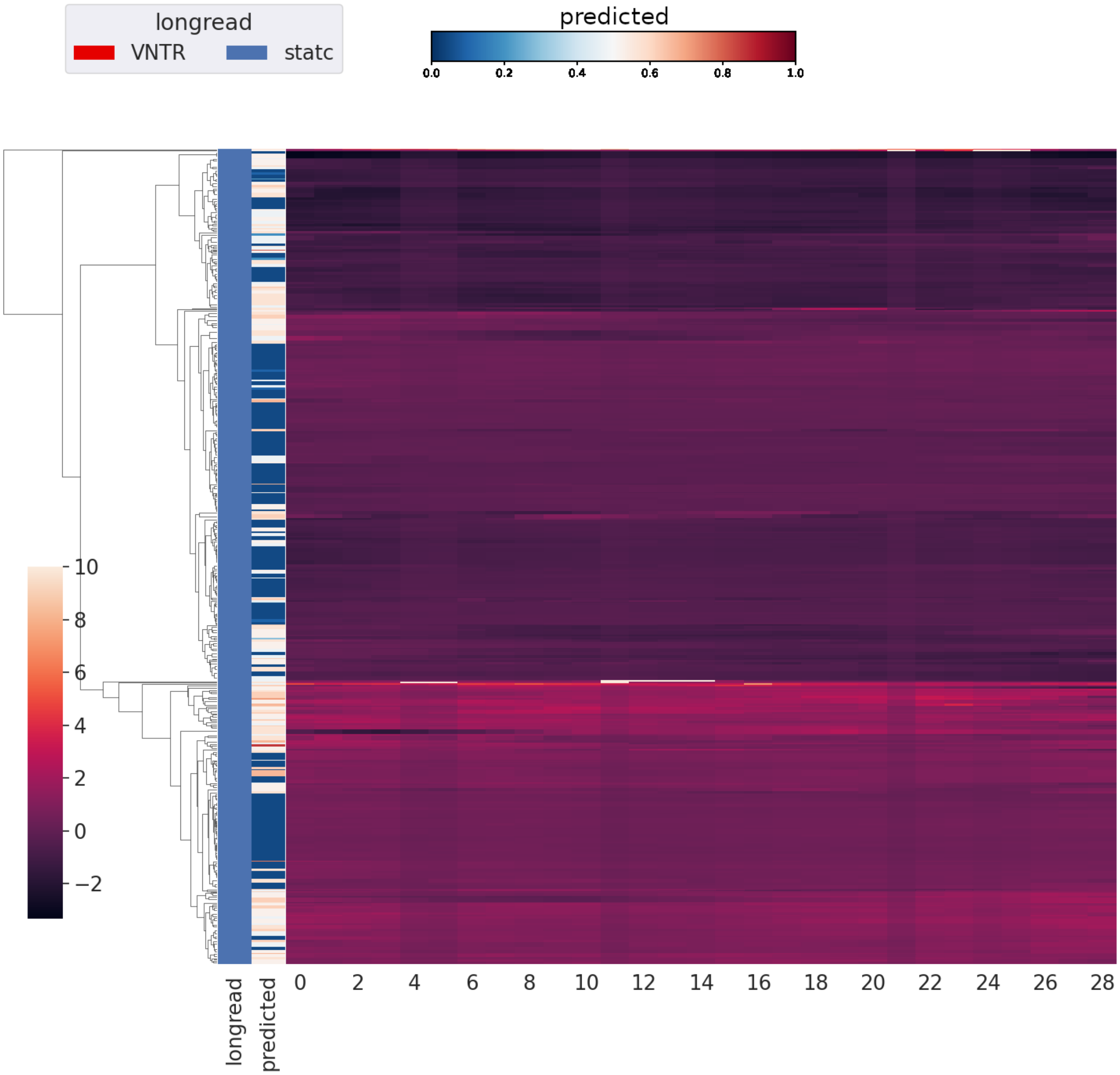


Raw MAD, static subset, continuous

**Fig. S32.** Continuous predicted probability labeling of longread labeled static TRs in full linkage hierarchical clustering map of MAD profiles over raw pileup in $17$ genomes, sample of $200$ per classifier prediction truthgroup.

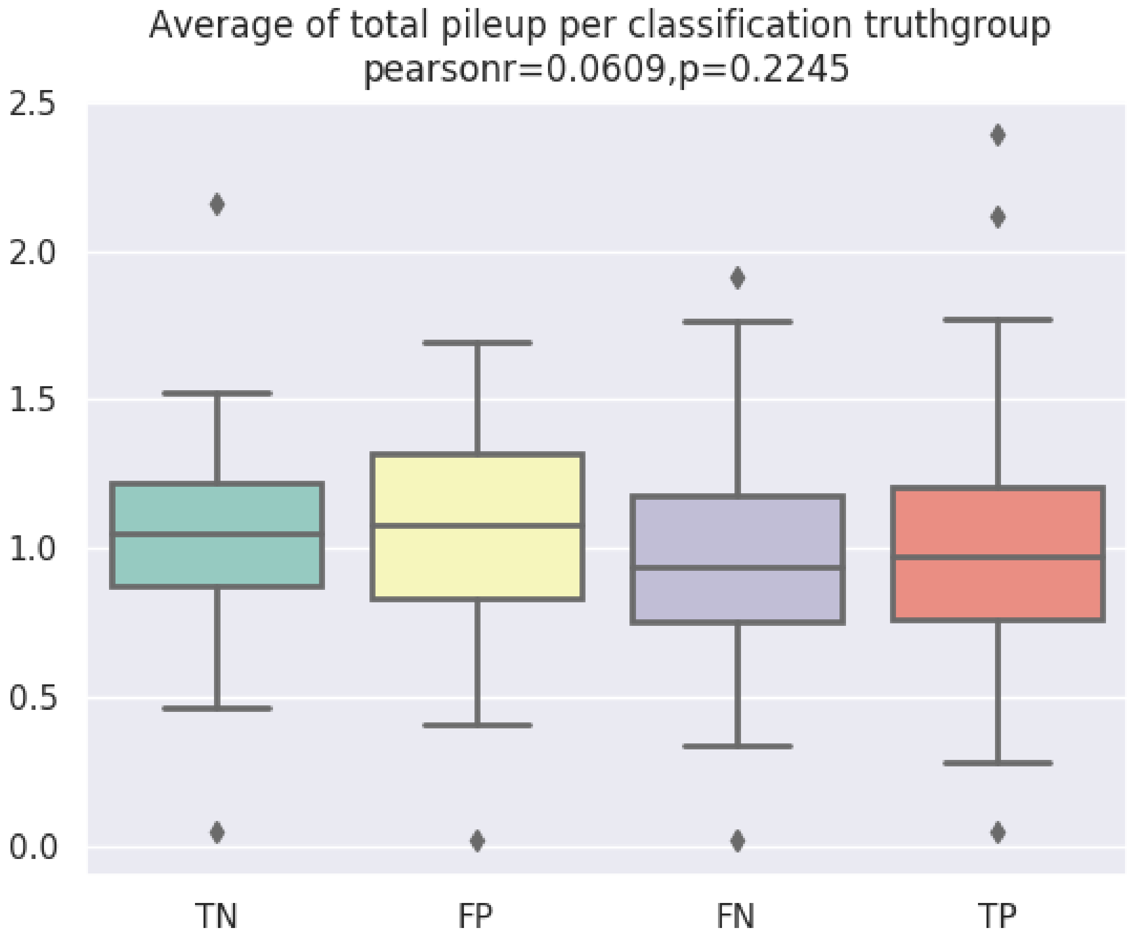


**Fig. S33.** Boxplot of average of total pileup per truth group of classifier prediction. Averages center around $1$ because of coverage normalisation (local pileup divided by average pileup in genome). The binary Pearson's r correlation is given for the FP group versus the TN group (candidates).

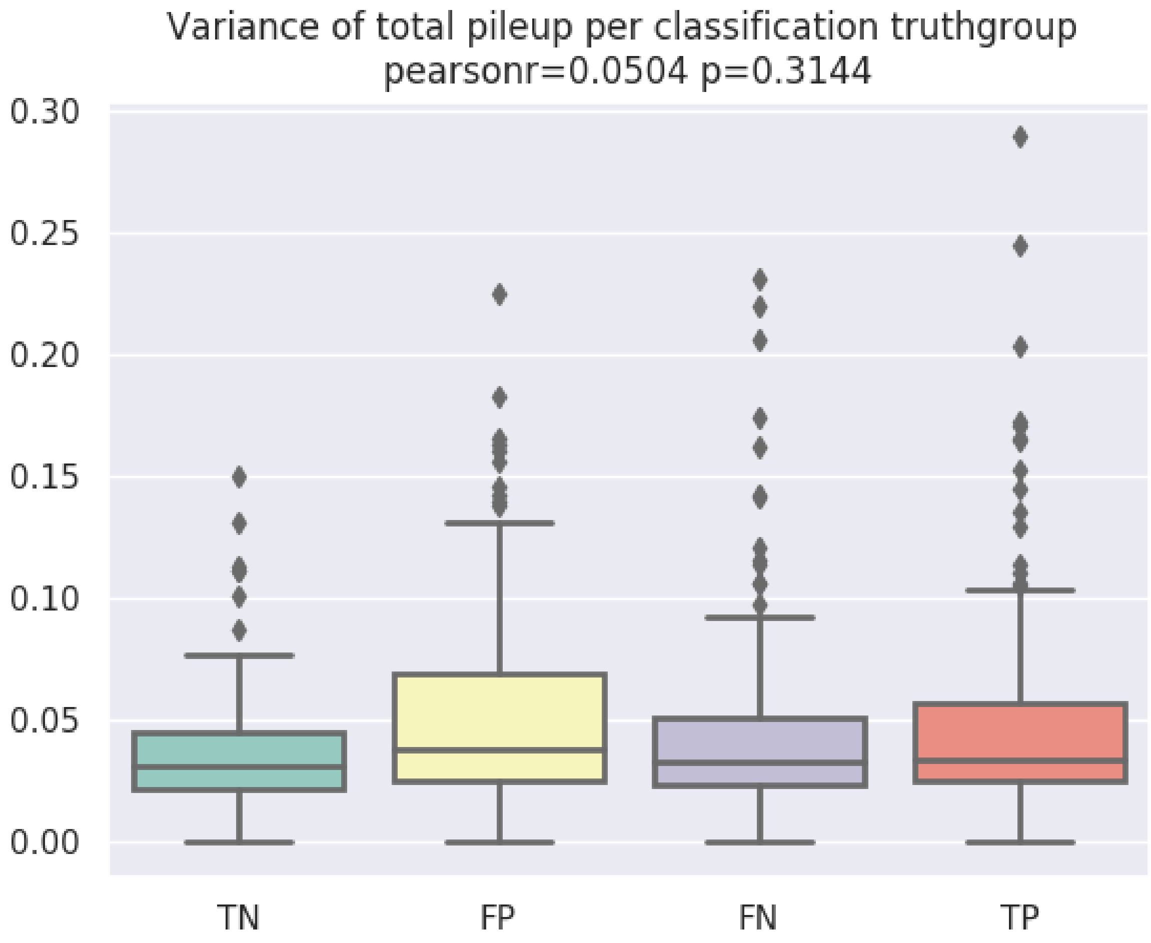


**Fig. S34.** Boxplot of variance in total pileup per truth group of classifier prediction. The binary Pearson's r correlation is given for the FP group versus the TN group (candidates).

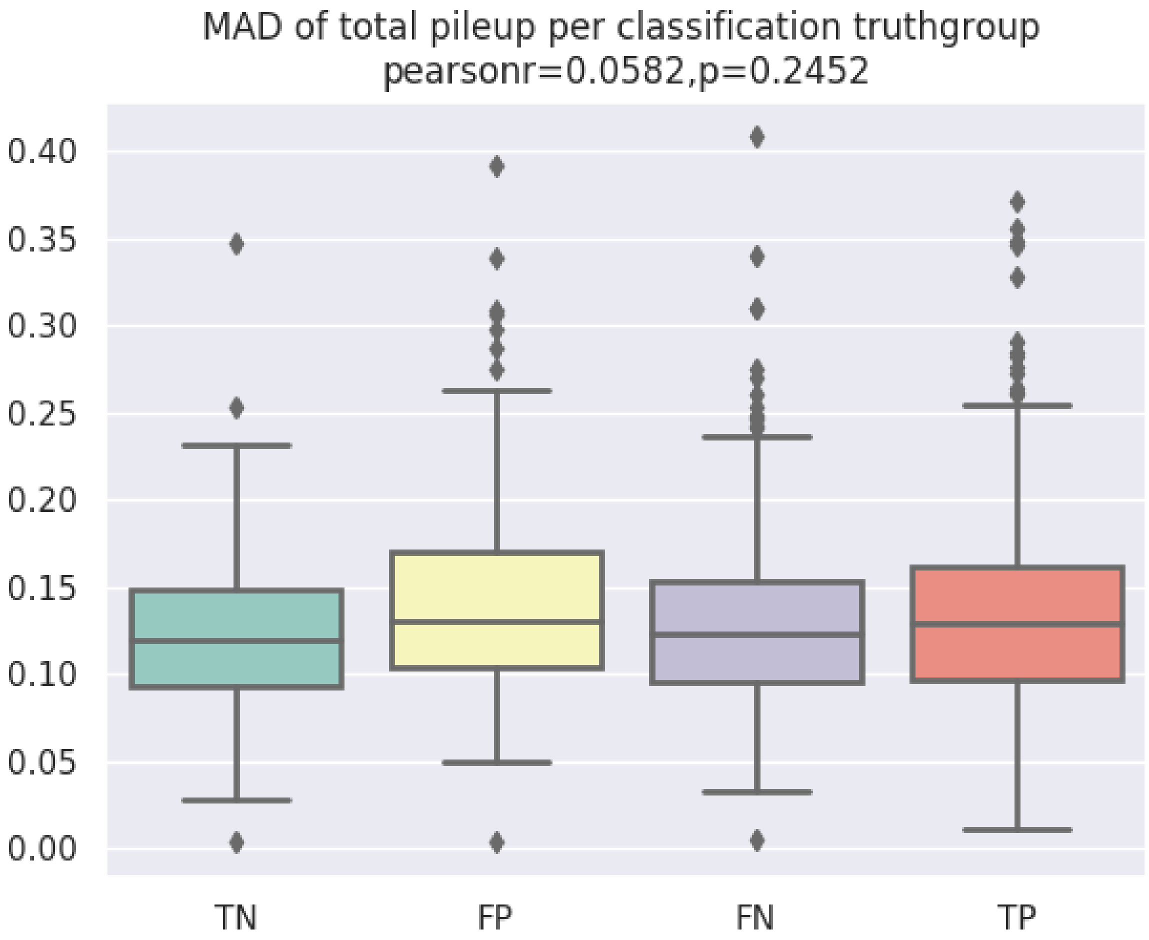


**Fig. S35.** Boxplot of MAD in total pileup per truth group of classifier prediction. The binary Pearson's r correlation is given for the FP group versus the TN group (candidates).

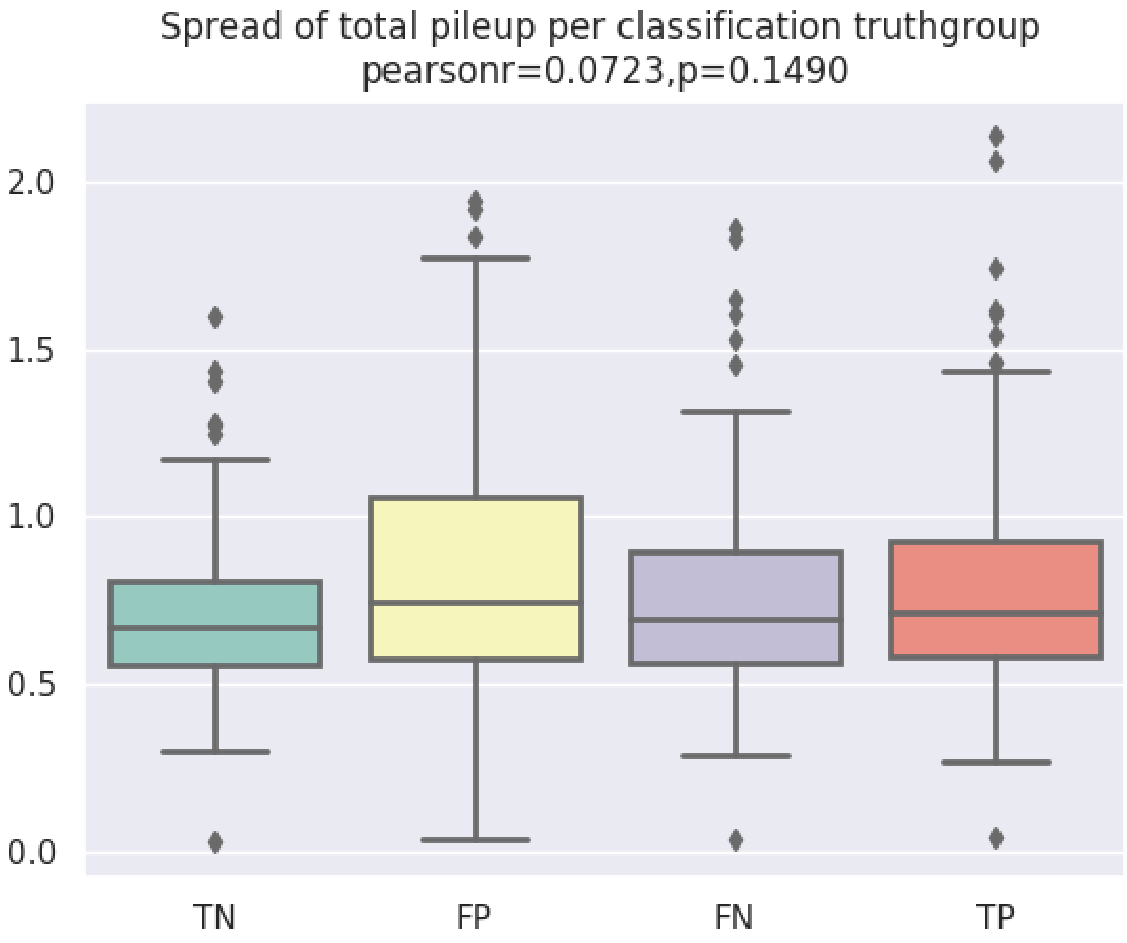


**Fig. S36.** Boxplot of spread in total pileup per truth group of classifier prediction. The binary Pearson's r correlation is given for the FP group versus the TN group (candidates).

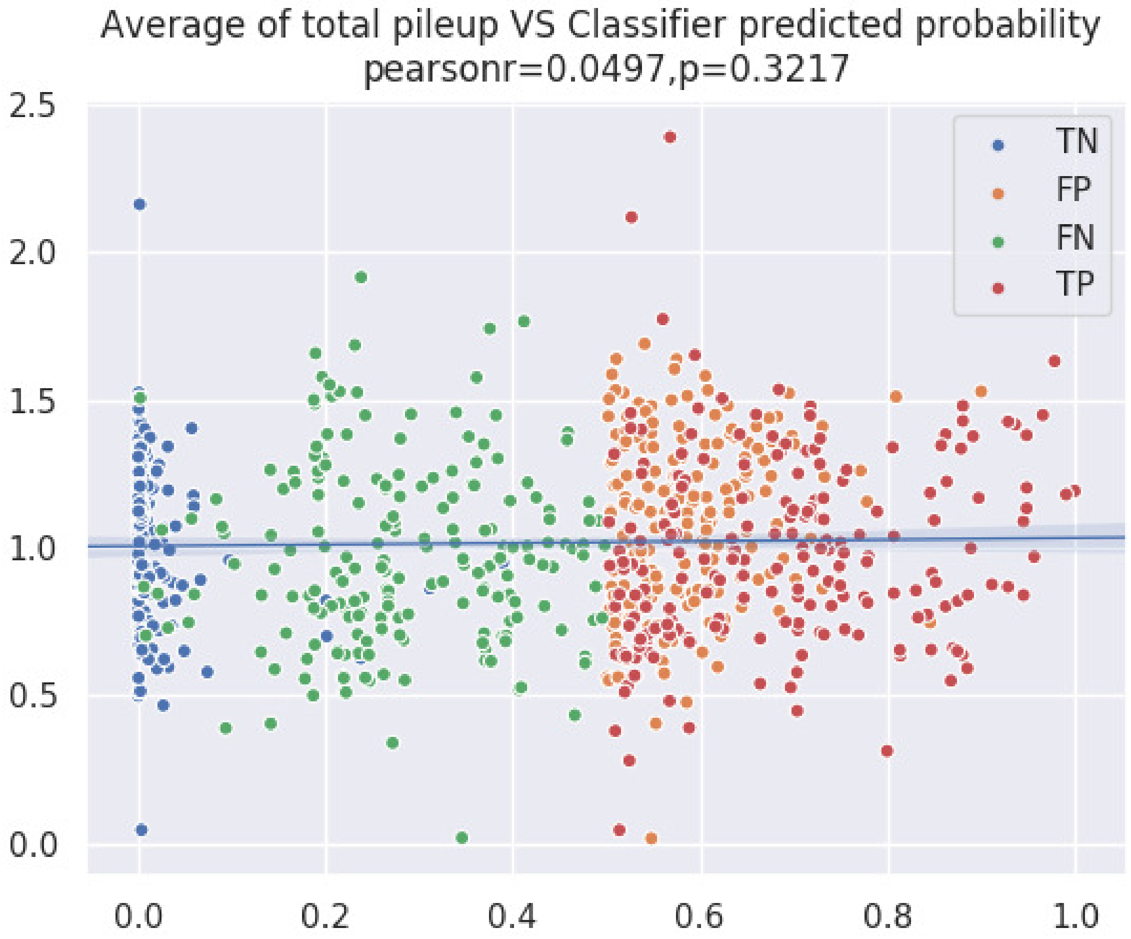


**Fig. S37.** Scatterplot of variance in total pileup per truth group of classifier prediction. The Pearson's r correlation is shown for the given axes and the blue line plotted is the linear regression.

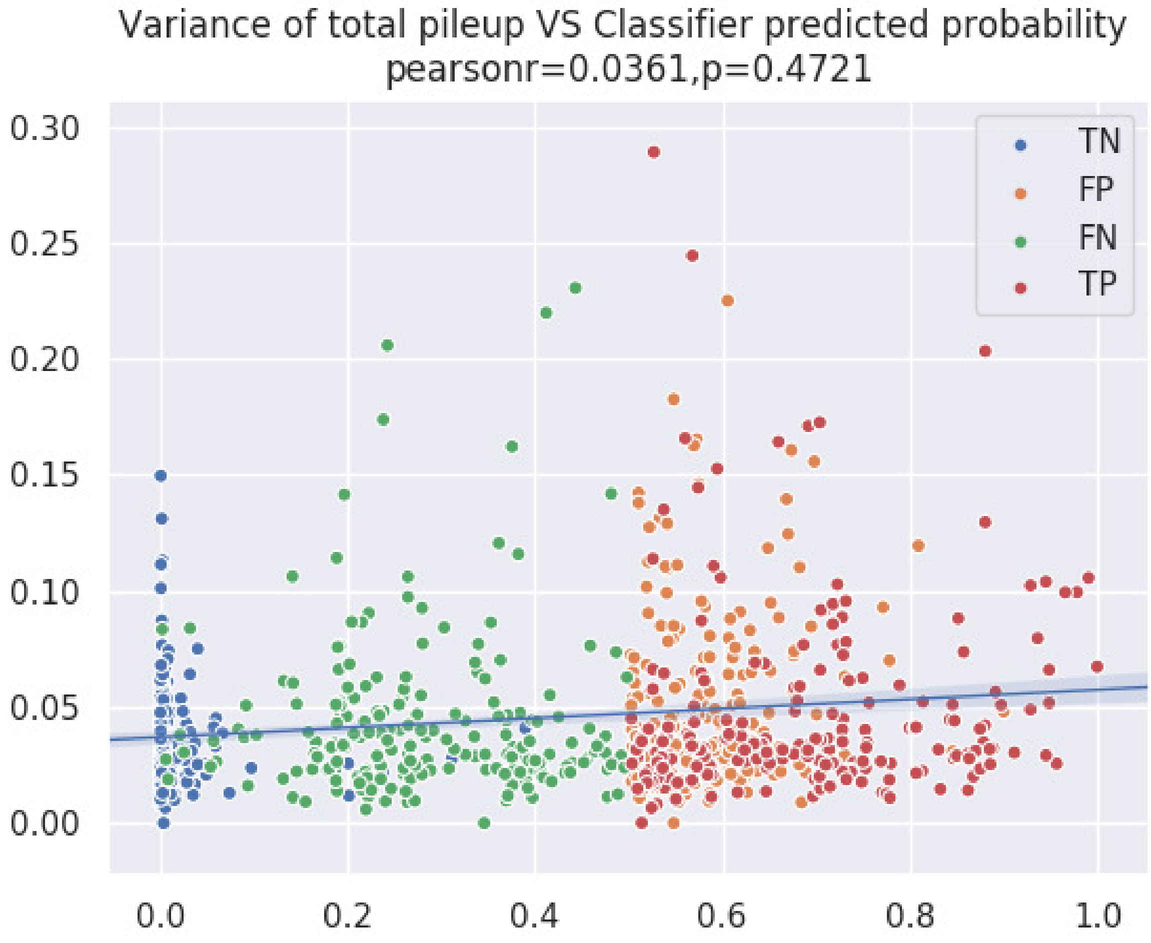


**Fig. S38.** Scatterplot of variance in total pileup per truth group of classifier prediction. The Pearson's r correlation is shown for the given axes and the blue line plotted is the linear regression.

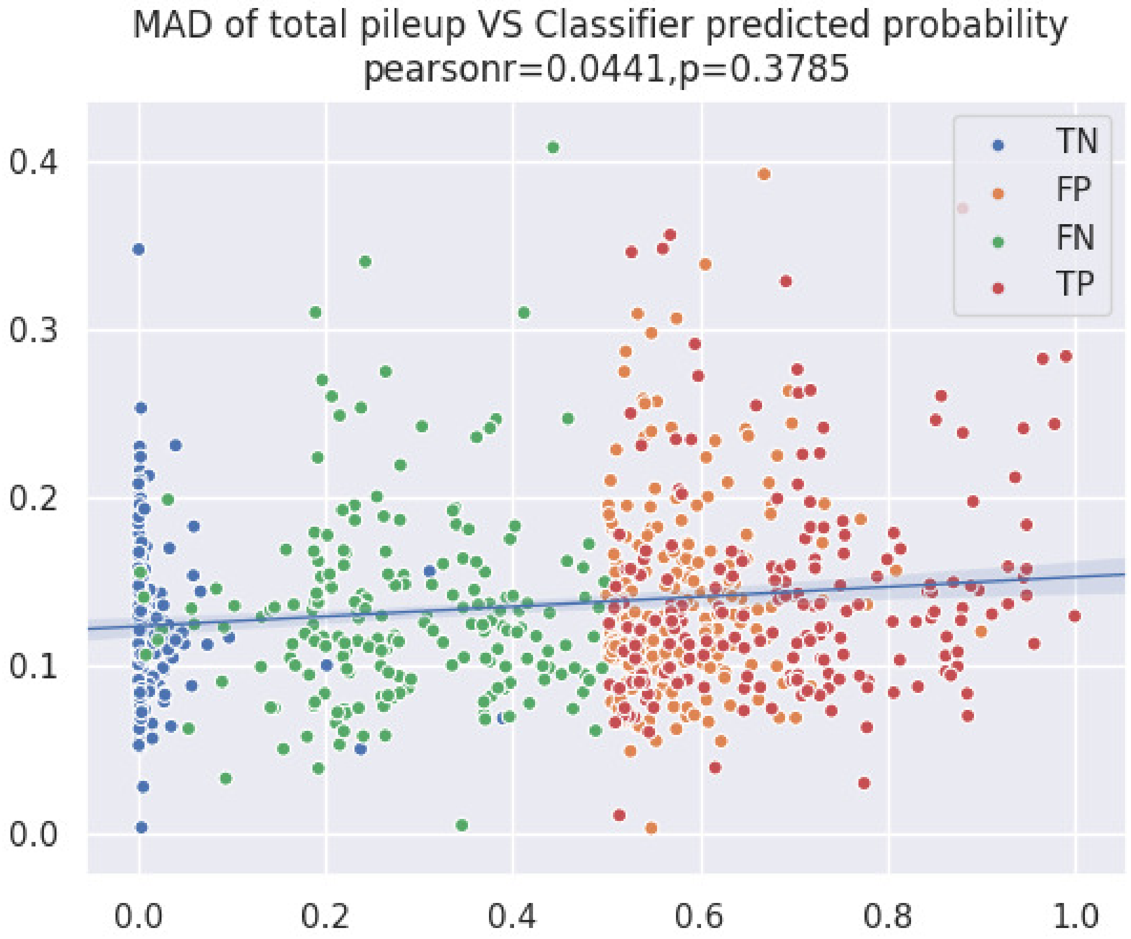


**Fig. S39.** Scatterplot of MAD in total pileup per truth group of classifier prediction. The Pearson's r correlation is shown for the given axes and the blue line plotted is the linear regression.

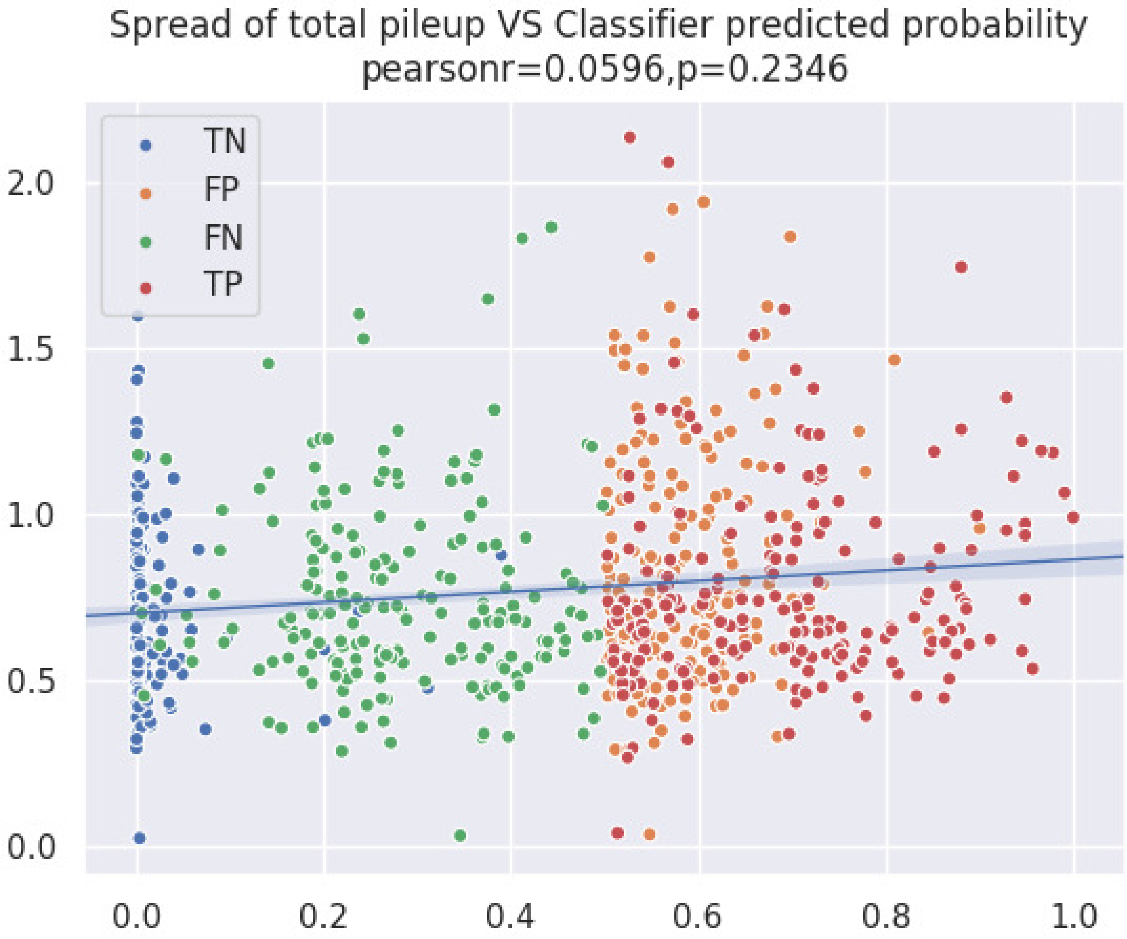


**Fig. S40.** Scatterplot of Spread in total pileup per truth group of classifier prediction. The Pearson's r correlation is shown for the given axes and the blue line plotted is the linear regression.

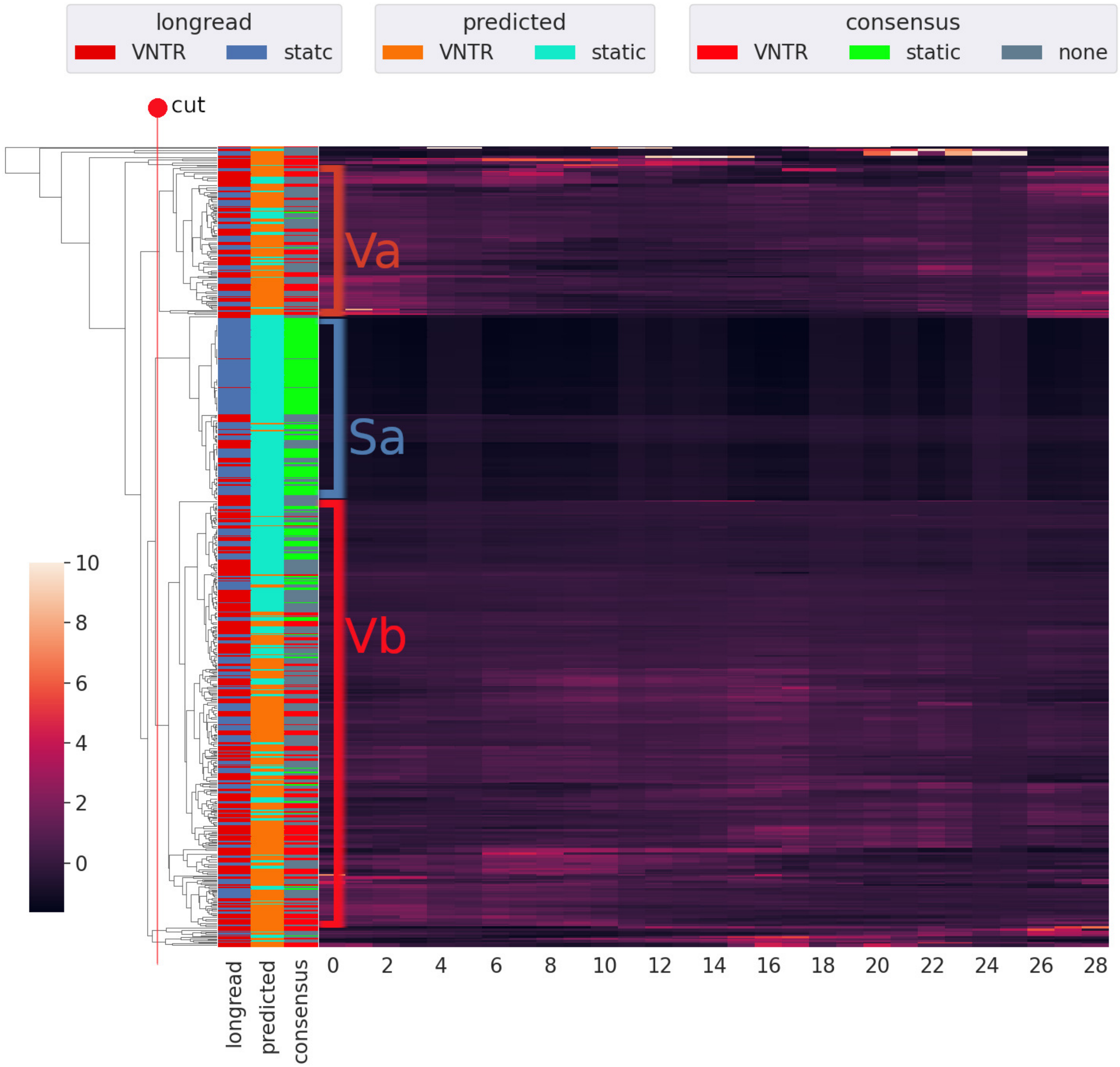


**Fig. S41.** Duplicate of Figure S21 with cut of clustering into semi pure agreed labeling in clusters.

**Table S1. Counts of label composition and FP enrichment in subclusters of Figure S41**

| Group | Total | Longread VNTR | Longread static | VNTR Agreement | static Agreement | FP in group | FN in group | TN in group | TP in group |
|---|---|---|---|---|---|---|---|---|---|
| Sa | 235 | 49 | 186 | 0 | 1 | 4 | 49 | 182 | 0 |
| Va | 193 | 103 | 90 | 0.9420289855 | 0.05797101449 | 83 | 41 | 7 | 62 |
| Vb | 552 | 328 | 224 | 0.7154811715 | 0.2845188285 | 156 | 157 | 68 | 171 |
